# RecPD: A Recombination-Aware Measure of Phylogenetic Diversity

**DOI:** 10.1101/2021.10.01.462747

**Authors:** Cedoljub Bundalovic-Torma, Darrell Desveaux, David S. Guttman

**Affiliations:** Department of Cell & Systems Biology, University of Toronto, Toronto, Ontario, Canada; Centre for the Analysis of Genome Evolution & Function, University of Toronto, Toronto, Ontario, Canada

**Author notes:** Corresponding author (DSG).

## Abstract

A critical step in studying biological features (e.g., genetic variants, gene families, metabolic capabilities, or taxa) underlying traits or outcomes of interest is assessing their diversity and distribution. Accurate assessments of these patterns are essential for linking features to traits or outcomes and understanding their functional impact. Consequently, it is of crucial importance that the metrics employed for quantifying feature diversity can perform robustly under any evolutionary scenario. However, the standard metrics used for quantifying and comparing the distribution of features, such as prevalence, phylogenetic diversity, and related approaches, either do not take into consideration evolutionary history, or assume strictly vertical patterns of inheritance. Consequently, these approaches cannot accurately assess diversity for features that have undergone recombination or horizontal transfer. To address this issue, we have devised RecPD, a novel recombination-aware phylogenetic-diversity metric for measuring the distribution and diversity of features under all evolutionary scenarios. RecPD utilizes ancestral-state reconstruction to map the presence / absence of features onto ancestral nodes in a species tree, and then identifies potential recombination events in the evolutionary history of the feature. We also derive a number of related metrics from RecPD that can be used to assess and quantify evolutionary dynamics and correlation of feature evolutionary histories. We used simulation studies to show that RecPD reliably identifies evolutionary histories under diverse recombination and loss scenarios. We then apply RecPD in a real-world scenario in a preliminary study type III effector protein families secreted by the plant pathogenic bacterium *Pseudomonas syringae* and demonstrate that prevalence is an inadequate metric that obscures the potential impact of recombination. We believe RecPD will have broad utility for revealing and quantifying complex evolutionary processes for features at any biological level.

**AUTHOR SUMMARY:** Phylogenetic diversity is an important concept utilized in evolutionary ecology which has extensive applications in population genetics to help us understand how evolutionary processes have distributed genetic variation among individuals of a species, and how this impacts phenotypic diversification over time. However, existing approaches for studying phylogenetic diversity largely assume that the genetic features follow vertical inheritance, which is frequently violated in the case of microbial genomes due to horizontal transfer. To address this shortcoming, we present RecPD, a recombination-aware phylogenetic diversity metric, which incorporates ancestral state reconstruction to quantify the phylogenetic diversity of genetic features mapped onto a species phylogeny. Through simulation experiments we show that RecPD robustly reconstructs the evolutionary histories of features evolving under various scenarios of recombination and loss. When applied to a real-world example of type III secreted effector protein families from the plant pathogenic bacterium *Pseudomonas syringae,* RecPD reveals that horizontal transfer has played an important role in shaping the phylogenetic distributions of aa substantial proportion of families across the *P. syringae* species complex. Furthermore, we demonstrate that the traditional measures of feature prevalence are unsuitable as a metric for comparing feature diversity.

## INTRODUCTION

The modern genomics era has provided unprecedented opportunities for identifying and quantifying the impact of genetic variants underlying traits of interest and furthering our understanding of the fundamental evolutionary processes driving the emergence, distribution, and fate of these variants. A critical step in studying genetic variants is assessing their distribution in a population and the overall diversity of the genetic factors underlying the trait. Accurate assessments of these patterns of diversity are essential for linking them to traits and understanding their functional impact. Consequently, it is critical that we have metrics that can accurately quantify diversity under any evolutionary scenario, including complex distributions brought about through recombination or horizontal gene transfer.

Prevalence, which is simply the proportion of individuals carry a feature of interest, and related ecological diversity metrics such as *richness* (i.e., the total number of features) and *diversity* (i.e., the number of features weighted by their prevalence, e.g., Simpson Index, Shannon Entropy, Hill Numbers) are some of the most common metrics used by researchers to compare the distributions of features of interest either within or between sampled populations [1–6]. In this context we use the term *feature* as a placeholder to refer to variation at essentially any biological level, such as the presence or absence of nucleotide or amino acid variants, gene families, species, operational taxonomic units (OTUs), metabolic capacities, phenotypic traits, or even different gene expression levels. Although prevalence can distinguish between the overall presence or absence of a feature of interest, it is agnostic to the evolutionary history of the organisms carrying that feature. Consequently, prevalence and related non-phylogenetic measures can be confounded by complex or unbalanced phylogenetic patterns, complex evolutionary histories, and biased sampling.

Phylogenetic diversity (PD) metrics have been developed that incorporate measures of evolutionary relatedness among individuals into the prevalence-based diversity metrics discussed above. In general, PD metrics are calculated by summing the branch lengths from a common ancestor of a selected group of descendants in a phylogenetic tree representing the total genetic diversity of the sampled population [1–3]. Two of the most widely used phylogenetic diversity metrics are Faith’s Phylogenetic Diversity [2] and UniFrac [7]. Faith’s PD calculates the sum of the phylogenetic tree branch lengths of all those branches that span the descendants that share the feature of interest. UniFrac measures the phylogenetic difference, or partitioning, between two feature distributions by calculating the proportion of branch lengths in a phylogenetic tree that leads to all the descendants that uniquely carry either feature of interest. Both Faith’s PD and UniFrac can work with cladograms that only represent the evolutionary branching patterns, or phylograms, where the branch lengths are proportional to evolution time and divergence. Additionally, both Faith’s PD and UniFrac not only incorporate organism relatedness in their calculation, but also implicitly allow for the loss the feature along phylogenetic lineages (e.g., by pseudogenization); thereby, making them excellent metrics for vertically inherited traits. To date, phylogenetic diversity metrics have largely been applied to the study of taxonomic diversity, and have been useful for identifying habitats which possess the greatest maximum biodiversity of a particular taxa/species to prioritize conservation efforts [8, 9], and comparing community diversity between sampling locations or under changing environmental conditions over time (e.g., calculating the beta-diversity of microbiomes) [4, 5, 10].

While phylogenetic diversity metrics have proven to be extremely useful, they all share a crucial underlying assumption that genetic diversity is vertically inherited, which limits their utility in the study of individual genetic loci. They can account for the loss of a feature but have no way to correct for non-vertical evolutionary processes such as recombination or horizontal gene transfer. While the assumption of vertical transmission is robust for many systems and studies, it absolutely is not appropriate when studying most microbial systems, where the horizontal transmission of genetic material is an important source of genetic novelty, functional innovation, and rapid adaptation.

Here we describe RecPD, a recombination-aware phylogenetic diversity measure, which addresses the inability of other phylogenetic diversity metrics to account for recombination and horizontal transfer. RecPD maps the distribution of a feature of interest onto a species tree and then employs a variety of ancestral state reconstruction approaches to infer the evolutionary histories of gain and loss which may have given rise to an observed distribution of that feature. Through simulation studies we show that RecPD can accurately account for diverse evolutionary scenarios involving recombination, which are ignored by currently available phylogenetic diversity measures. We derive a number of RecPD-based metrics that summarize the impact of horizontal transfer, and then show the utility of the approach when analysing the distribution of type III secreted effector protein families carried by the plant pathogenic bacterium *Pseudomonas syringae*.

## RESULTS

### Development of RecPD

The development of RecPD was inspired by the need to understand the distribution of bacterial gene families, so we will discuss the methods from this context although the method is transferable to any other features of interest (e.g., genetic variants, metabolic capabilities, or taxa). We begin with a phylogeny of strains on which we will map the acquisition, loss, descent, and divergence of a gene family of interest. In most circumstances, this phylogeny will be based on the core genome (i.e., those genes found in all strains of the species) and be constructed from the concatenated sequences of core genes. For simplicity, we will refer to this as the species tree. We also have presence / absence distribution of a gene family of interest that varies among the strains in the study set. The goal of RecPD is to determine the phylogenetic diversity of the gene family by reconstructing its evolutionary history on the species tree, accounting for potential horizontal acquisition events that may have occurred.

### Step 1: Assignment of gene family ancestral states on the species phylogenetic tree

To reconstruct the putative lineages where a gene family has arisen during the evolutionary history of a bacterial species, we first begin with ancestral state reconstruction of the gene family. In this case ancestral states considered will be a binary category of gene family presence/absence. To achieve this task, we devised a novel nearest-neighbour (NN) ancestral reconstruction approach (**Fig 1A**) that begins by identifying which strains in the study set carry the gene family of interest. Based on this, the tips of the species tree are assigned a state of presence or absence for the gene family. Next, each internal/ancestral node is examined and assigned to one of three possible states based on supporting information from the closest-related tips descended from it: 1) ‘present’ if the nearest-neighbour, i.e., closest related, descendant tips of the node both possess the gene family; 2) ‘absent’ if the gene family is absent in the nearest-neighbour descendant tips; 3) ‘split’ if only one nearest-neighbour descendant tip possesses the gene family, which may indicate potential gain/recombination or loss events.

**Fig 1.**
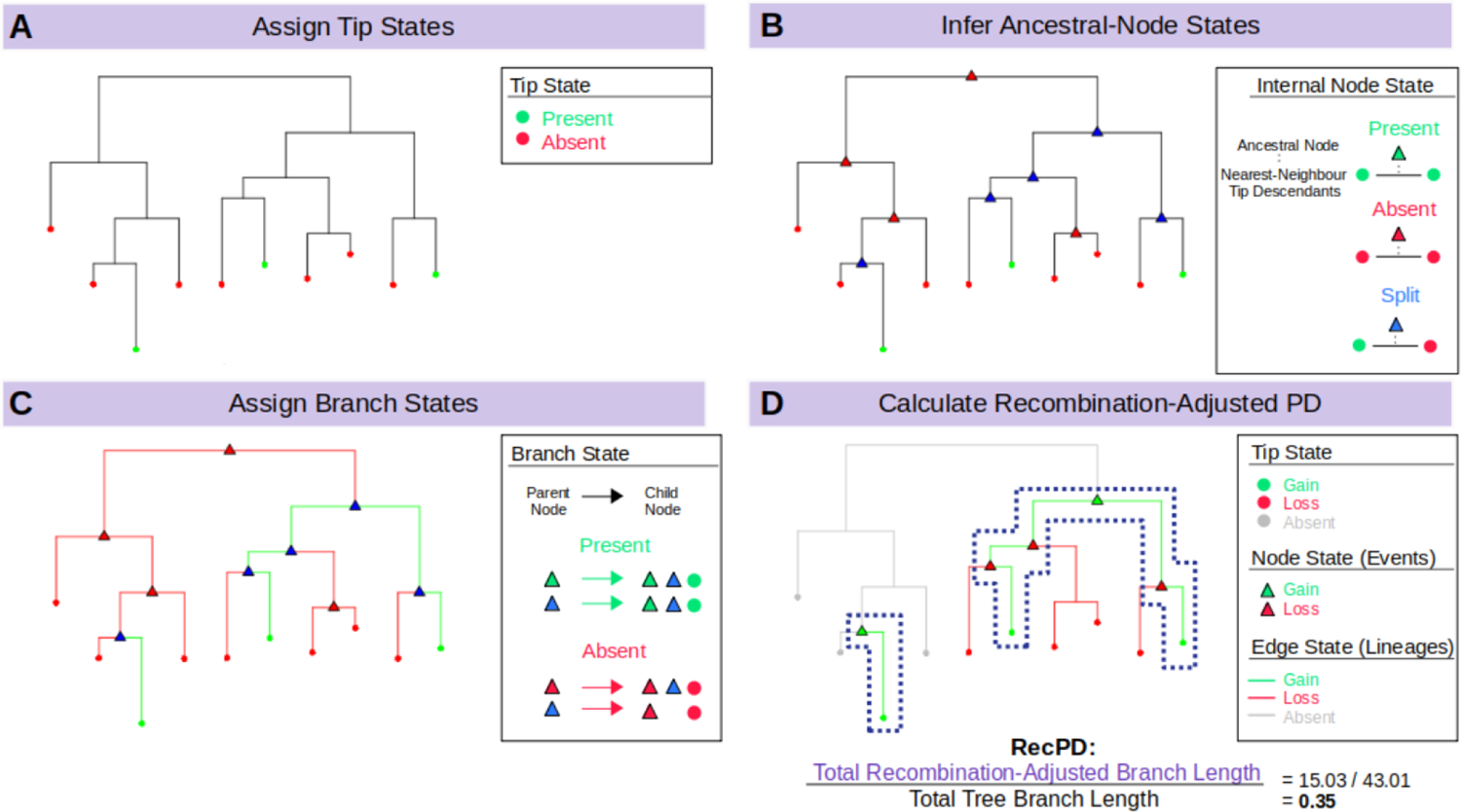
Outline of the RecPD Methodology. (A) Tip states are assigned to a given species phylogenetic tree based on presence (green), or absence (red) of the gene family of interest. (B) Ancestral node states are inferred using one of there ancestral state reconstruction methods and assigned to either presence (green), absence (red), or split (blue) states, the latter which indicate potential gain/loss events. (C) Branches are assigned to presence states if they join consecutive presence or split nodes/tips (green), otherwise they are assigned to absence (red). (D) Gene-family lineages are identified, and split state nodes are assigned to gains (green) or losses (red). Branches descended from the phylogenetic tree root node where no ancestral presence nodes were identified are assigned to absence (grey). RecPD is then calculated as the sum of gene family lineage branch-lengths normalized by the total branch-lengths of the phylogenetic tree.

Our framework also allows the incorporation of other popular approaches for ancestral state reconstruction, such as most-parsimonious reconstruction (MPR) [11] and maximum-likelihood ancestral character estimation (ACE) [12]. The goal of MPR is to find the overall set of internal node states which results in the fewest number of state changes, e.g., most-parsimonious, between ancestral and descendant nodes. The ACE approach was devised as an improvement on MPR, which incorporates branch-length information and inferred gain and loss rates from which the likelihood of given state for each internal node determined (i.e., from the likelihoods of the states of its descendants).

As the reconstruction of ancestral states is not a trivial task [13, 14], it is important to emphasize that reconstructed ancestral states can vary depending on the approach used, so it is useful to state the potential strengths and limitations inherent to each. The NN approach may produce spurious evolutionary histories of ancestral gene family loss followed by reacquisition depending on the frequency of gain and loss in different lineages. MPR is less liable under these scenarios but does not incorporate divergence between ancestral and descendant nodes. Therefore, it may miss certain internal nodes where a state might be present or could find equally parsimonious scenarios resulting in ambiguous state assignments. In the case of ACE, modelling gains and losses as separate processes may resolve ambiguities in some scenarios, but reliably estimating these rates largely depends on the sampling of the given species phylogeny. State assignments will tend to become more uncertain as the amount of divergence between ancestral nodes increases. A combination of approaches could be used in theory to find consensus assignment, although this method has not yet been developed.

### Step 2: Identification of gene family lineages

Ancestral node state assignments made in the previous step are then consolidated to delineate the species lineages and ancestral phylogenetic tree branches where the gene family is likely to have arisen (**Fig 1B**). This is done by examining all the branches of the species tree and assigning a presence state to branches where ancestral-child node states where the gene family has also been predicted to be present, otherwise a given branch is assigned as an absence state. Some branches may also include nodes where a split/ambiguous state has been assigned as the result of potential gain or loss events occurring in descendant node lineages. In this case, branches are selected for inclusion if the ancestral or child node is assigned to the presence state, or both are split states/ambiguous.

After branch-state identification, the final node states are consolidated, with gain and loss event nodes highlighting the particular internal nodes where the gene family appears to have been gained or lost in subsequent descendant lineages respectively (**Fig 1C**). In addition, the states of tip nodes descended from each gain or loss lineage are updated accordingly.

### Step 3: Calculation of RecPD

After the corresponding gain and loss lineages for a given gene family distribution has been determined, its recombination-adjusted phylogenetic diversity, RecPD, is calculated by summing the total branch-lengths of the gain lineages divided by the sum of the total branch-lengths of the species tree (**Fig 1D**).

### Step 4: RecPD-Derived Metrics

In addition to RecPD, we can also calculate additional metrics based on the ancestral mapping of the gene family onto the species tree. These metrics can be divided into two classes that either assess the overall topological structuring of gene families within the species tree or summarize the evolutionary events influencing gene family lineages (i.e., those lineages of the species tree where the gene family is predicted to be present) inferred by RecPD. The first class of topology-based metrics include: *Span*, which measures how much of the maximum possible species diversity is realized for the gene family of interest and is akin to the Faith’s PD metric (S1 Fig); and *Clustering*, which measures the extent to which the gene family lineages identified by RecPD partition among subclades of the species tree, which may be more or less closely related (S2 Fig). While Span and Clustering are conceptually very similar, the former uses branch lengths, while the latter is a cladistic measure that only considers clade structure. The second class of evolutionary event-based metrics include: *Longevity*, which is the median normalized evolutionary distance since a gene family was initially gained across species lineages (S3 Fig); and *Lability*, the normalized number of gain, loss, and re-acquisition event nodes occurring across species lineages for a given gene family (S3 Fig).

We also developed a comparative metric for directly quantifying the extent of shared ancestry (i.e., correlation) between different gene families, called RecPDcor. For a pair of gene family distributions, RecPDcor is calculated as the sum of the ancestral lineages branch-lengths where they co-occur divided by the sum of branch-lengths from the union of their ancestral lineage reconstructions (S4 Fig). This is in essence the branch-length weighted Jaccard similarity between ancestral gene family lineages.

### RecPD accounts for potential recombination events from randomized gene family phylogenetic distributions

As a preliminary exploration of our RecPD method, we present an idealized test-case scenario using a randomly generated species tree of 10 tips and assessed the potential impact of recombination inferred from examining all possible gene family presence/absence patterns. We calculated Faith’s PD for each gene family phylogenetic pattern to serve as a baseline evolutionary scenario considering only vertical descent, gene family loss, and no recombination. Fig 2A shows that RecPD was affected by the choice of the ancestral state reconstruction method used (NN, MPR, or ACE). When compared to Faith’s PD, ACE tends to overestimate PD, which is likely due to increasing uncertainty in state assignments for the deepest ancestral nodes the species tree. In contrast, both the NN and MPR methods consistently predict lower PD values than Faith’s, particularly for gene family distributions found in at least half of the species in the tree, with MPR predicting the greatest degree of recombination (Fig 2B & C).

**Fig 2.**
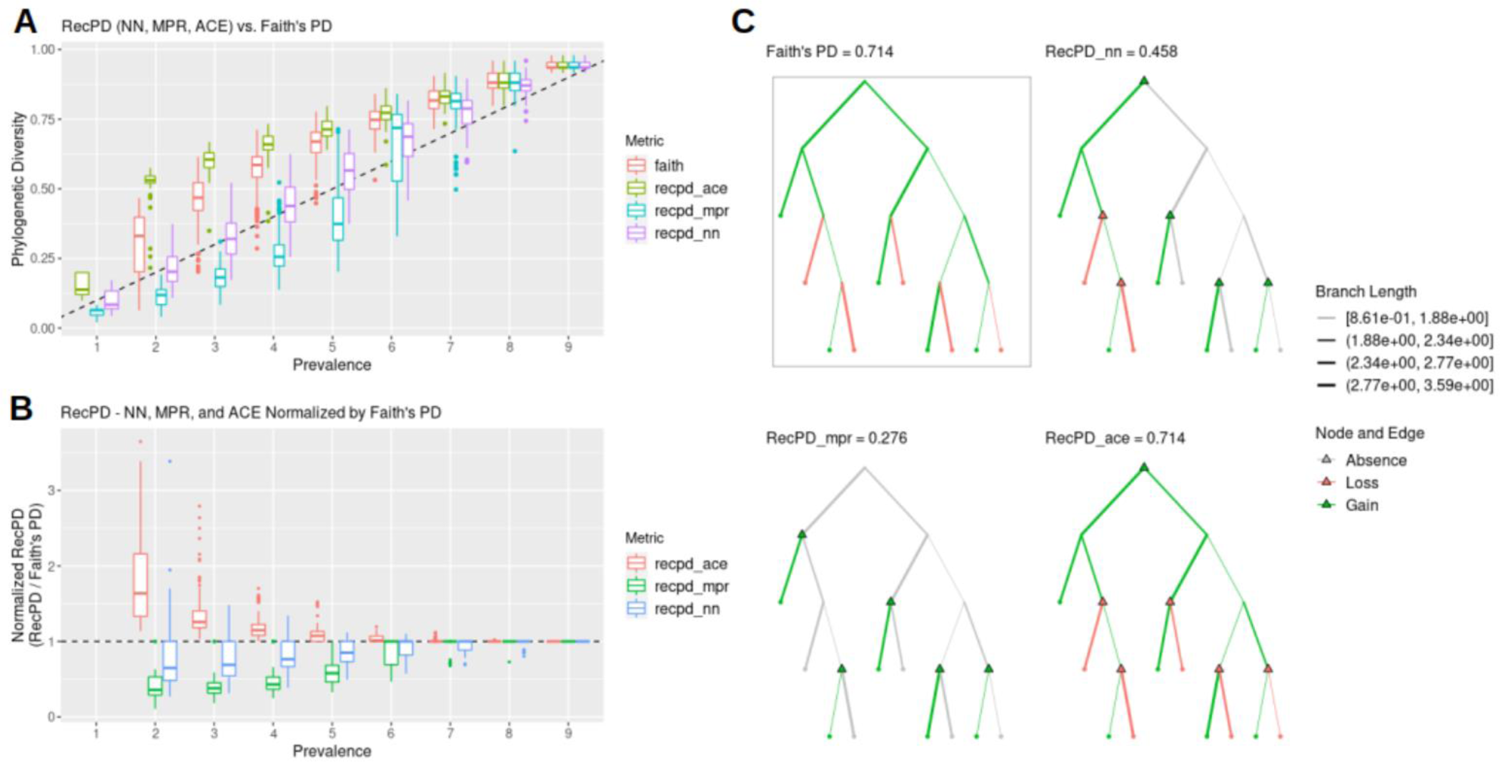
RecPD using the NN, MPR, and ACE ancestral reconstruction approaches compared to Faith’s PD. (A) RecPD using three ancestral state reconstruction methods and Faith’s PD distributions binned by gene family prevalence. Results shown correspond to all 1022 possible randomized gene-family distributions mapped onto a tree of ten tips. (B) RecPD normalized by Faith’s PD for three ancestral state reconstruction methods binned by gene-family prevalence. Results shown correspond to all 1022 possible randomized gene-family distributions mapped onto a tree of ten tips. (C) Example gene family distribution of prevalence = 5 illustrating differences in inferred evolutionary events using different methods: Faith’s PD = 5 losses, 0 gains (boxed), RecPD(NN) = 2 losses, 4 gains, RecPD(MPR) = 0 losses, 5 gains, and RecPD(ACE) = 5 losses, 0 gains.

Therefore, RecPD can serve to identify potential gene family distribution patterns which do not conform to a strictly vertical pattern of inheritance. These results also hold for randomly generated species trees of greater size (S5 Fig).

### Nearest-neighbours ancestral state reconstruction accurately captures recombination events of simulated gene family evolutionary histories

In the real world, we almost never know the true evolutionary histories of a gene family. Although ancestral reconstruction methods are valuable as a means for reconstructing evolutionary histories, more thorough analyses of genomic data are required for support. Therefore, we simulated gene family histories on randomly generated species trees using a Poisson process to model recombination and loss (Table 1, S6 Fig). With these ‘known’ gene family histories we evaluated how accurately the different ancestral state reconstruction approaches, NN, MPR, and ACE, perform in identifying recombination events under diverse evolutionary scenarios, e.g., loss-recombination balanced, loss dominant, or recombination dominant.

**Table 1.** Gene family evolutionary history simulations: parameters used.

| Tree Size<br>(Ntips) | # of Trees<br>(N_trees) | Longevity<br>Rate<br>(long_r) | Loss Rate<br>(death_r) | Recombina-<br>tion Rate<br>(recomb_r) | # Rate<br>Parameter<br>Sets | Simulations<br>per Set<br>(N_iter) | # of<br>Simulations |
| --- | --- | --- | --- | --- | --- | --- | --- |
| 10 | 10 | 0,1 | 0,2,4,6,8,10 | 0,2,4,6,8,10 | 72 | 25 | 5894 |
| 50 | 10 | 0,1 | 0,2,4,6,8,10 | 0,2,4,6,8,10 | 72 | 25 | 8173 |
| 100 | 10 | 0,1 | 0,2,4,6,8,10 | 0,2,4,6,8,10 | 72 | 25 | 8147 |

To determine the accuracy of RecPD and each ancestral reconstruction approach (NN, MPR, and ACE) we calculated the ratio between the calculated RecPD values and the known PD based on the simulated distribution. As before, Faith’s PD was also calculated as a baseline comparison assuming no recombination. The results of our simulation using a tree of ten tips (Fig 3) show that Faith’s and RecPD using ACE result in similar error rates, largely overestimating the PD of gene families, recapitulating results shown previously (mean error Faith = 0.46 ± 0.68; mean error ACE = 0.57 ± 0.78). Conversely, RecPD using MPR underestimated gene family PD (error MPR = −0.18 ± 0.24). Surprisingly, our newly devised NN method was shown to be the most accurate (mean error = 0.083 ± 0.28) in correctly reconstructing gene family histories evolved under high-recombination scenarios. These results were consistent across different evolutionary scenarios (S7 Fig) and for other simulations using randomly generated species trees of ranging from 50 to 100 tips, with the observed error decreasing with increasing tree size (S8 Fig).

**Fig 3.**
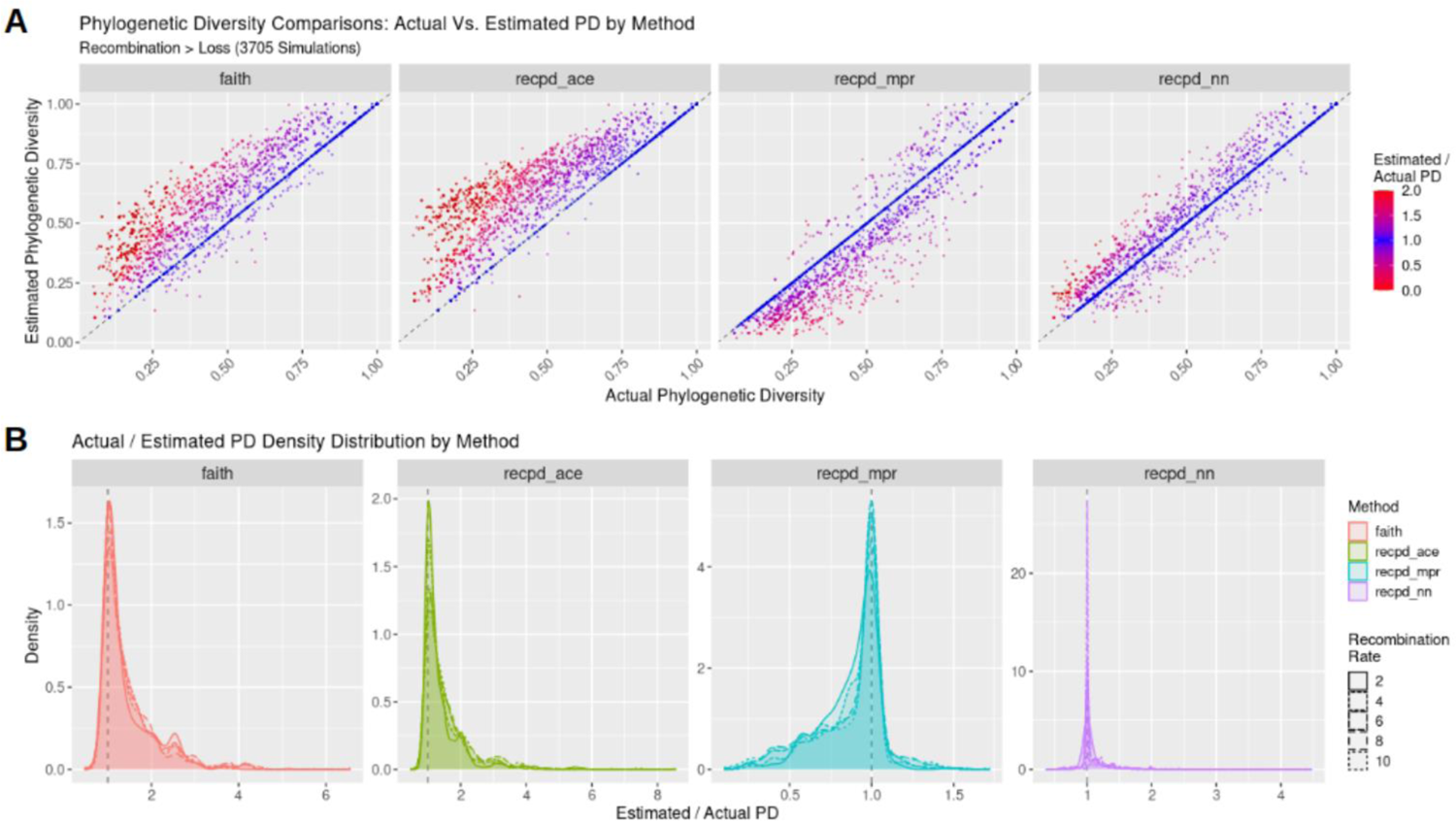
RecPD NN, MPR, and ACE vs. Faith’s PD for simulated gene family distributions. Simulations of gene family evolution comparing actual phylogenetic diversity to estimated diversity using Faith’s PD and RecPD with three different ancestral reconstruction methods (see Table 1 for parameters used and number of simulations run). (A) Scatterplots of estimated PD values against actual PD values by method. (B) Corresponding density plot distributions. Results are shown for recombination predominant rate regime on randomly generated trees with 10 tips.

### RecPD identifies gene family distributions with shared and unrelated evolutionary histories

We next explored the use of RecPD in the context of identifying gene families with shared evolutionary histories. From the RecPD NN ancestral reconstructions for all randomized gene family distributions generated in our first analysis, we calculated the RecPDcor values of their recombination adjusted evolutionary histories, resulting in 521,731 unique pairwise comparisons (note that identical distributions were excluded). In the case of gene family distributions with identical prevalence, we observed that RecPDcor values can vary considerably (Fig 4), particularly for distributions of lower prevalence, reflective of their greater possibility of having evolved through recombination (see Fig 2). Similarly, RecPDcor values tended to be lower for gene family distributions with greater relative differences in prevalence, which is to be expected as the result of limited overlap in evolutionary histories (S9 Fig).

**Fig 4.**
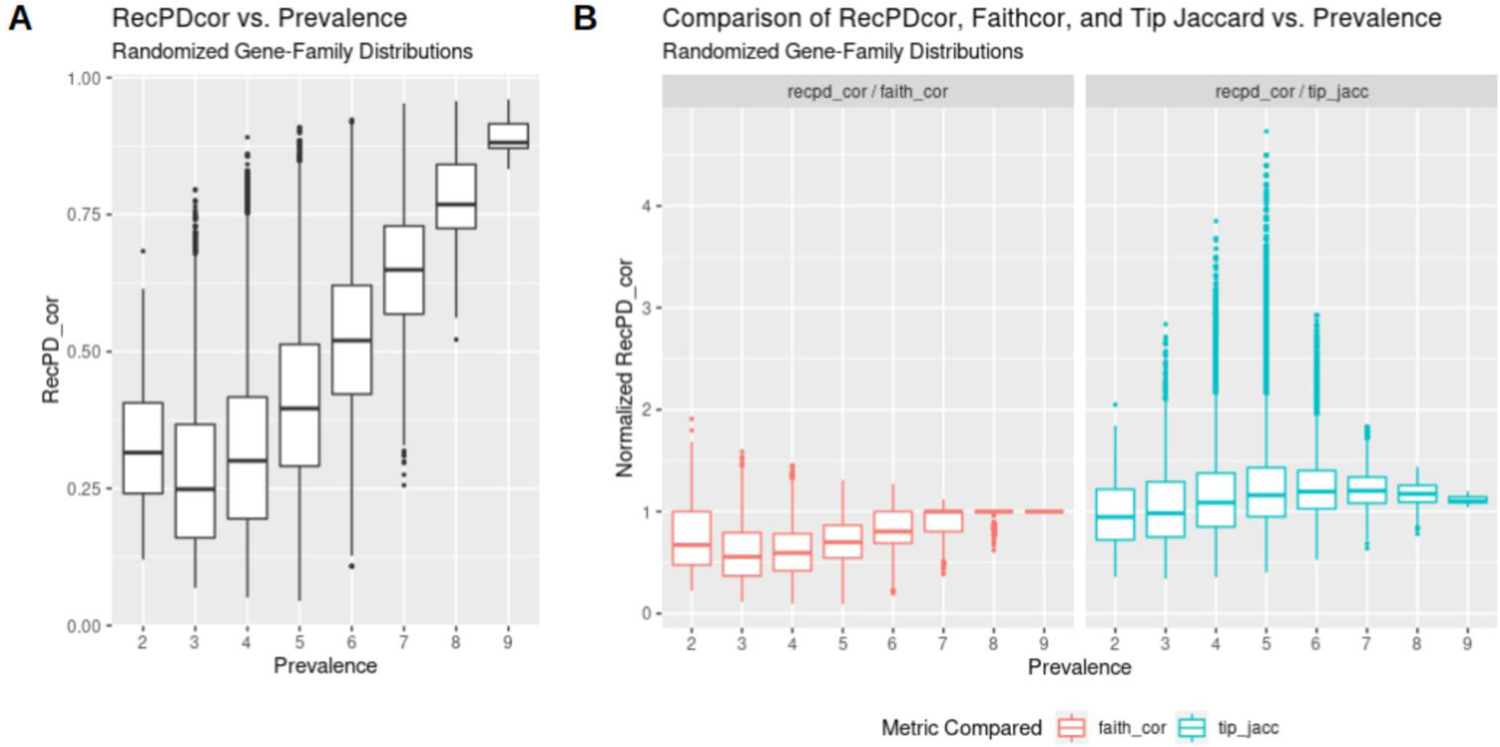
Pairwise gene family evolutionary history correlations using RecPDcor. (A) RecPD correlation (RecPDcor) values for randomized gene family distributions vs. prevalence; (B) RecPDcor normalized by Faith’s PD based branch-length weighted Jaccard similarity (recombination-agnostic) and tip presence and absence Jaccard similarity (phylogeny-agnostic).

We also compared RecPDcor against other correlation metrics to examine the effect of recombination in determining gene family co-occurrence (Fig 4B). The two metrics employed were a Faith’s PD-based (faith_cor) branch-length weighted Jaccard similarity, which ignores recombination and assumes vertical ancestry of gene families, and Jaccard similarity of co-occurrence among species/tips of the tree (tip_jaccard), which ignores gene family ancestries. It was observed that the correlations of gene family distributions tended to be overestimated when not accounting for potential recombination, while ignoring their evolutionary history altogether resulted in their underestimation. In the latter case, by comparing the differences between RecPDcor and tip Jaccard similarities, we could identify several instances where tip Jaccard either missed or identified spuriously correlated gene families (Fig 5).

**Fig 5.**
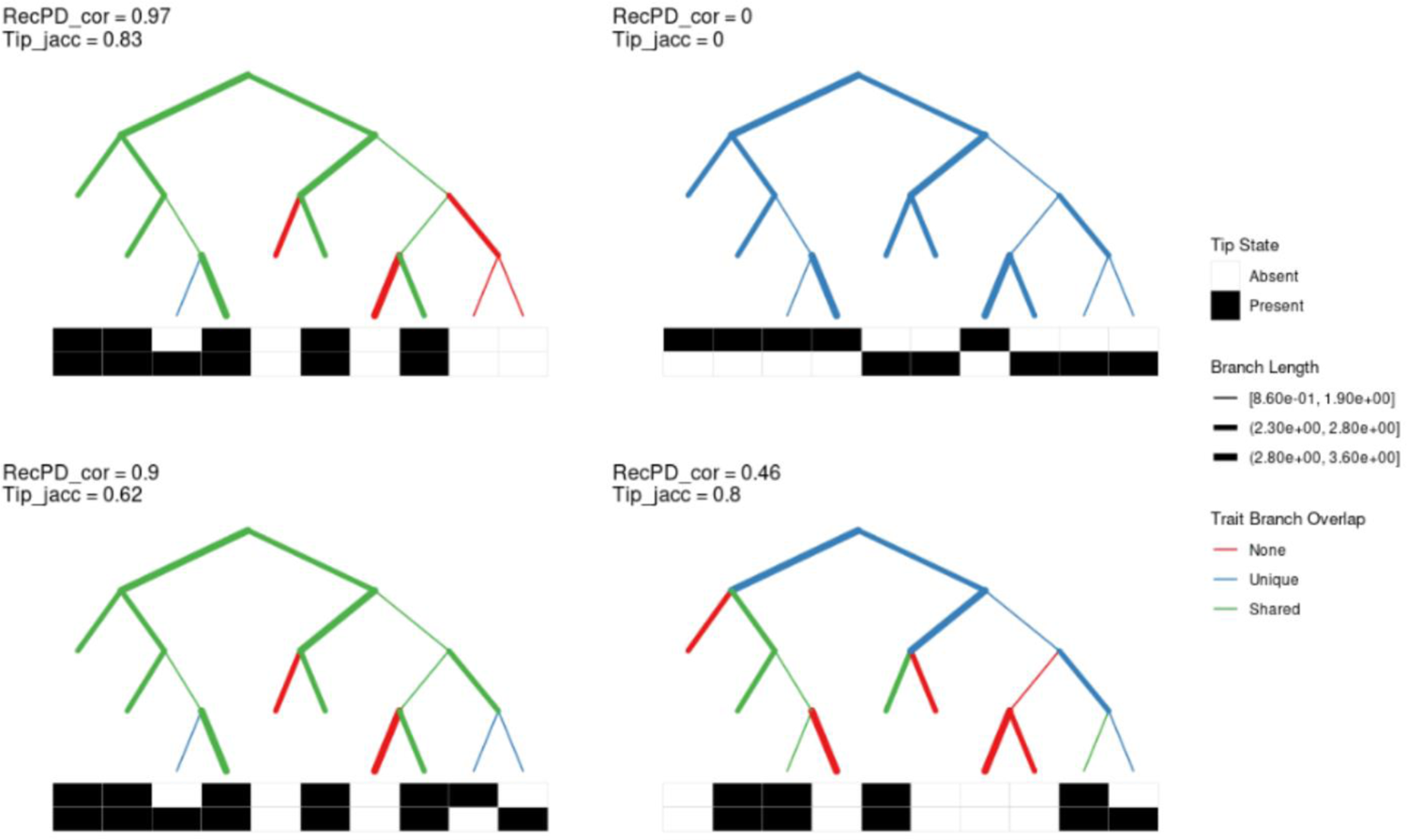
RecPDcor vs. tip Jaccard similarity example. Top panels – Distributions of two features (black = present, white = absent) arrayed against a species tree. RecPDcor and tip Jaccard similarities both identify correlated and anti-correlated gene families. Bottom panels – Distributions where RecPDcor reveals correlated gene family distributions where tip Jaccard does not, and where tip Jaccard overestimates correlation of gene families with distinct evolutionary histories. Tree topologies are represented as cladograms with branches of equal length; actual branch-lengths are indicated by branch-thickness as indicated in the legend. Branches are coloured according to overlap between RecPD-inferred gene family lineages.

### RecPD characterization of Pseudomonas syringae type III secreted effector families

*Pseudomonas syringae* is a highly diverse phytopathogenic bacterial species complex that includes over 60 pathogenic varieties that cause many agronomically important crop diseases [15–18]. Decades of research have established *P. syringae* as an important model for the study of host-pathogen interactions. One factor that makes *P. syringae* a particularly adept phytopathogen is its use of a type III secretion system and diverse repertoires of type III secreted effector proteins (hereafter effectors), which have evolved to promote disease by disrupting host immunity and cellular homeostasis [19, 20]. In turn, plant hosts have evolved a layer of immunity that triggers when receptors recognize the presence or activity of pathogen effectors [19, 21–23]. As a result, the outcome of any particular host-pathogen interaction, and pathogen host specificity in general, largely depends on the specific effectors carried by the pathogen and the specific immune receptors carried by the host. These interactions have led to a co-evolutionary arms race and the accumulation of extensive effector and immune diversity. The *P. syringae* species complex has at 70 characterized effector families, most of which include numerous diverse alleles that have evolved through both vertical and horizontal evolutionary processes [15]. There is huge diversity in the suites of effectors carried by individual *P. syringae* strains, with most strains carrying ∼30 effectors (±9 stderr) [15].

We applied RecPD to a previously published dataset of 529 representative effector alleles distributed among the 70 effector families identified from a collection of 494 sequenced *P. syringe* strains [15, 24]. *P. syringae* strains are classified into phylogroups based their placement in a core genome phylogenetic analysis, with phylogroups varying in overall size and diversity. We mapped effector alleles onto the core genome (i.e., species) tree and found wide variation in the prevalence and distribution of families (Fig 6A) [15]. Relatively few effector families are widely conserved across the *P. syringae* species complex, and effector families of similar levels of prevalence can be distributed in very different ways across the phylogroups, suggesting the importance of recombination and loss in effector evolution [15, 24].

**Figure 6.**
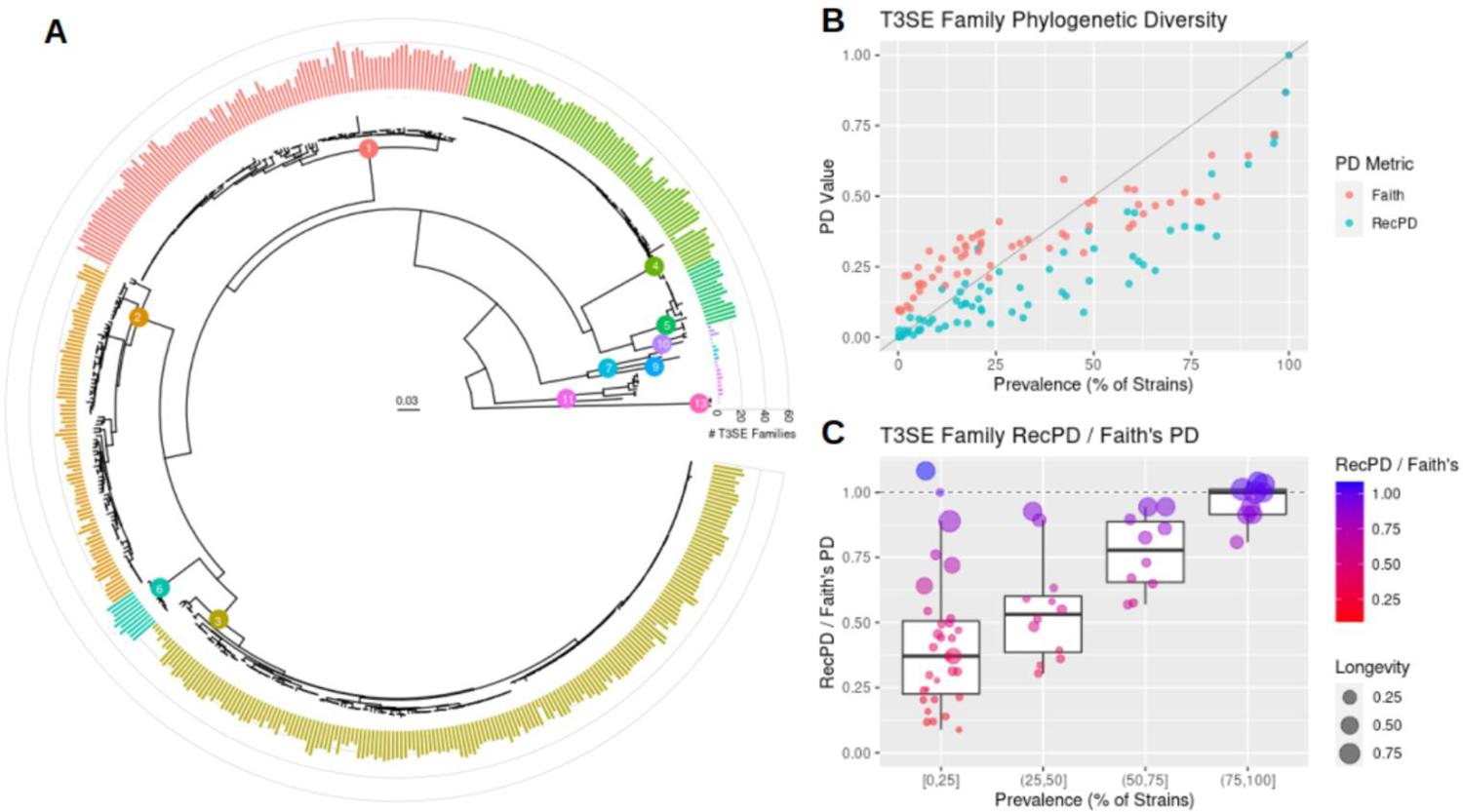
The Phylogenetic Distribution of *P. syringae* effector families. (A) Core genome phylogeny of the *P. syringae* species complex, with internal tree nodes indicating *P. syringae* phylogroups. The outer-ring barplot shows the total number of distinct effector families carried by each strain and coloured according to strain phylogroup designation. (B) Plot of effector family prevalence against RecPD(NN) and Faith PD for all 70 effector families. (C) Effector family RecPD values normalized by Faith’s PD, binned by effector family prevalence. The point size indicates effector family longevity.

We used RecPD to gain a better understanding of effector evolutionary diversity. Not unexpectedly, we find that effector phylogenetic diversity is positively correlated with prevalence, however we note that relying on prevalence alone gives a misleading view of the phylogenetic distribution of effectors as a whole (Fig 6B). Notably, effector family RecPD values were lower than might be expected from effector prevalence because phylogenetic diversity, unlike prevalence, takes into account the structuring of gene families across evolutionarily distinct bacterial clades and the different degrees of evolutionary diversity within each. This is of particular importance when certain clades might be under sampled, and others might consist of many nearly identical strains (as in observed for the current *P. syringae* dataset). Furthermore, we also note the importance of considering the effects of recombination when calculating phylogenetic diversity. In the instance of lower-prevalence effector families (found in less than ∼ 25% of strains), Faith’s PD values tend to be larger than expected based on the observed prevalence, possibly due to impact of horizontal gene transfer that distributes many of these families across multiple phylogroups (i.e., deeper common evolutionary ancestry including branches with greater evolutionary divergence). The impact of horizontal transfer is much more evident from the RecPD values, which are consistently lower than the corresponding Faith’s PD values. This is made even more clear when we normalize the PD values by calculating the ratio of RecPD to Faith’s PD. Since Faith’s PD assumes strictly vertical ancestry, families with a high ratio (∼ 1) can be interpreted as evolving by largely vertical evolutionary processes, while a low ratio supports extensive horizontal transmission. We found that the extent of recombination can vary widely for effector families, even when they have very similar prevalence (Fig 6C). The longevity (median normalized evolutionary distance since an effector family was gained) also appears to be correlated to the RecPD - Faith’s PD ratio. Taken together, this analysis supports the hypothesis that the majority of effector families have experienced extensive horizontal transfer and have been acquired relatively recently during the evolutionary history of *P. syringae*.

To concretely illustrate the impact of horizontal transfer and utility of RecPD, we highlight two examples. The first is the effector families HopS and HopAW (Fig 7A), which have nearly identical prevalence, with HopS found in 114 strains and HopAW found in 116 strains, but dramatically different distributions and inferred evolutionary histories. The Faith’s PD values for the two families are 0.152 and 0.198 for HopS and HopAW, respectively. HopS, with a RecPD value of 0.164, appears to largely follow vertical descent from an early acquisition event in the evolutionary history of *P. syringae* and is highly conserved in phylogroup 1 and 6 strains. In contrast, HopAW, with a RecPD value of 0.048, appears to have been gained and lost at numerous times throughout the *P. syringae* phylogroups. The second example is the effector families HopH and HopBN (Fig 7B) which have different prevalence (206 vs. 78 strains for HopH and HopBN respectively), but nearly identical RecPD values of ∼ 0.16. Their corresponding Faith’s PD values are 0.31 and 0.34, indicating similar overall distribution across the *P. syringae* species phylogeny despite their difference in prevalence. However, by visually comparing their reconstructed evolutionary histories it can be seen that HopH and HopBN have followed dramatically different evolutionary trajectories. The RecPD derived metrics also paint a very different evolutionary picture for each family, as shown in Table 2.

**Fig 7.**
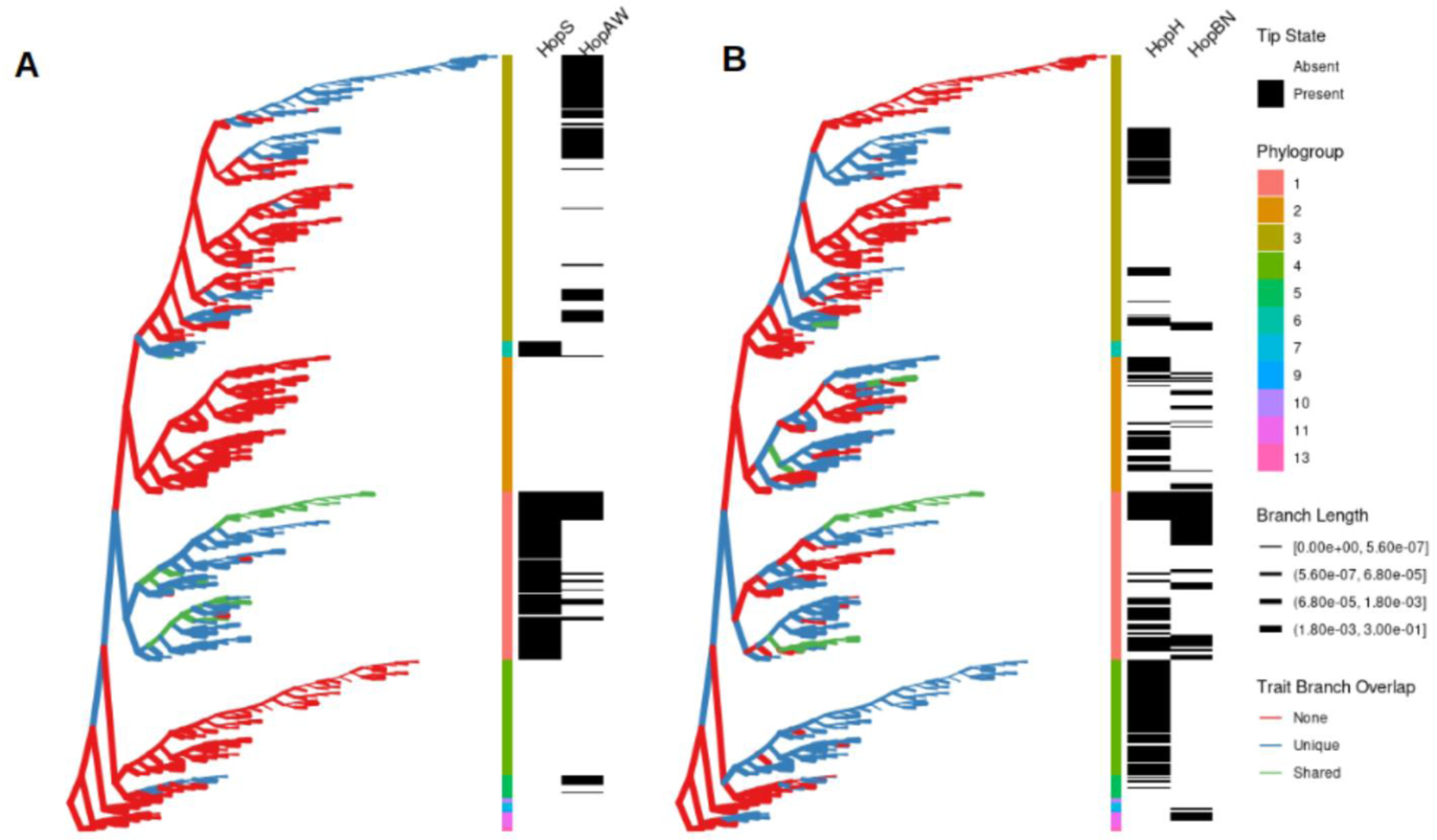
RecPD inferred effector family histories. Example pairs of effector family distributions mapped onto the *P. syringae* core-genome phylogeny. (A) Effector families HopS and HopAW show similar prevalence (HopS = 114 and HopAW = 116) but different RecPD values (HopS = 0.399 and HopAW = 0.198). (B) Effector families HopH and HopBN show different prevalence HopH = 206 and HopBN = 78) but similar RecPD values (0.16). Tree topologies are represented in a ‘willow tree’ format, with branches set to equal length, and actual branch-lengths indicated by branch-thickness. Branches are coloured according to overlap between RecPD-inferred gene family lineages.

**Table 2.** RecPD and associated metrics for selected effector families shown in Fig 7.

| **T3SE**<br>**Effector**<br>**Family** | **Prevalence** | **Faith's PD** | **RecPD(NN)** | **Span** | **Clustering** | **Longevity** | **Lability** |
| --- | --- | --- | --- | --- | --- | --- | --- |
| HopS | 114 | 0.152 | 0.164 | 0.225 | 0.991 | 0.574 | 0.015 |
| HopAW | 116 | 0.198 | 0.048 | 0.269 | 0.896 | 0.004 | 0.147 |
| HopH | 206 | 0.313 | 0.160 | 0.369 | 0.937 | 0.031 | 0.115 |
| HopBN | 78 | 0.342 | 0.161 | 0.400 | 0.818 | 0.017 | 0.073 |

## DISCUSSION

Ecological diversity metrics, such as phylogenetic diversity, have been used to gain valuable insight into the complexity of ecological communities. These metrics can be used to quantify diversity both within and between species or communities (alpha and beta diversity respectively), and to assess richness (i.e., how many), divergence (i.e., how different), and regularity (i.e., how uniform). Put another way, these metrics assess the sum, mean, and variance of the phylogenetic differences among organisms [1]. Despite their tremendous utility, all these metrics have a common underlying assumption that raises concerns about their applicability for the majority of life on Earth. Specifically, they assume that organisms under study have descended from a common ancestor strictly through vertical descent. While this may be a reasonable assumption for eukaryotes, it is certainly less valid for bacteria and archaea, where horizontal gene transfer can occur within and between species and dramatically influence their adaptive capabilities.

We developed our recombination-aware phylogenetic diversity metric RecPD to provide a framework for understanding ecological and evolutionary diversity that is robust to the presence of horizontal gene transfer and recombination. RecPD utilizes ancestral state reconstruction to infer the evolutionary histories of features of interest (e.g., gene families, metabolic or phenotypic traits, taxa, etc.) in the context of a given phylogeny. It then identifies evolutionary gains and losses of these features on the tree and quantifies the diversity for only that fraction of the species tree where the feature is present. It then enables the calculation of a number or related metrics that quantify the amount of recombination that has occurred in the history of the feature. In general, RecPD provides an intuitive metric for comparing the diversity and impact of recombination on a large number of features. It also provides a way to identify lineages that have gained a feature of interest via horizontal transfer, which may otherwise be difficult to determine with features that do not have a clear pattern of descent.

An important step in calculating RecPD is the reconstruction of ancestral states to infer gain and loss events. RecPD can use either of the two well-established ancestral state reconstruction approaches that use distinct modelling frameworks, e.g., parsimony (MPR) and maximum likelihood (ACE). We also introduce a novel approach that is based on nearest-neighbouring states (NN). When evaluating the accuracy of each ancestral reconstruction approach we found that MPR and ACE led to under- and over-estimation of the phylogenetic diversity of simulated gene family histories, respectively, reflective of the assumptions and limitations particular to each modelling framework. We found that the NN ancestral state reconstruction approach provided the most accurate reconstructions, and importantly, performed robustly under evolutionary scenarios of elevated recombination and loss. In addition, we also showed that correlation of ancestral lineages could serve as a useful extension of traditional genomic-context approaches to assess potential functionally linked gene families [25, 26]. However, we caution that the reconstructed gene family histories are still best-guesses given the data at hand, and predicted horizontal transfer events should be used as a starting point for validation using other methods, e.g., conservation of genomic neighbourhood, association with mobile genetic elements, GC content, nucleotide diversity, or species-gene tree topological concordance [27].

The utility of RecPD is clear when analyzing gene families such as *P. syringae* effectors that function as both virulence factors and immune elicitors. These effectors are subject to strong selective pressures, frequent horizontal transfer, and pseudogenization. In a preliminary study of *P. syringae* effectors, we demonstrate that RecPD provides greater insights into effector diversity and evolution than non-phylogenetically aware methods.

In addition to quantify the impact of recombination, RecPD may also be of value in analytical approaches that need to control for population structure, such as genome wide association studies (GWAS). Bacterial GWAS approaches are heavily dependent on population structure corrections that control for the evolutionary history of the sample [28–34]. The ability of RecPD to identify and quantify recombination may increase the power of these statistical methods and provide an interesting avenue for future development. In general, RecPD has great potential for quantifying diversity and assessing the contributions of vertical and horizontal modes of evolution, which is of critical importance for understanding the processes driving the evolution of many bacterial and archaeal gene families

## METHODS

### RecPD Development and Implementation

All code development, final implementation of RecPD analyses, simulation experiments and figures presented in this work was performed in RStudio (R version 4.0.2) [35]. The ape library [36] was used for basic phylogenetic tree import and processing tasks, generation and visualization of random trees used in methods development section, and ancestral reconstruction using MPR and ACE. Faith’s PD was calculated using the pd() function from the picante library [37]. The ggtree library [38] was used for final phylogenetic tree visualizations. Code for running RecPD can be found in **S1 File**.

### Gene family evolutionary simulation

Simulation gene-family history evolution was performed using a poisson process to model gene family recombination and loss events onto a provided species tree phylogeny (code supplied in **S2 File**). The method is motivated by ideas from the modelling of birth-death phylogenetic trees [39], similar approaches used for simulating microbial gene-tree phylogenies [40], and the simulation of bacterial genomes and phenotypic evolution on phylogenetic trees [33]. In essence, it models the evolution of a trait (gene-family, variant, or locus) through loss and recombination occurring along the different lineages of a provided species phylogenetic tree. The tree can be either ultra-metric or non-ultrametric, with branch-lengths representing either time since emergence from a common ancestor, or a molecular evolutionary distance (average expected nucleotide/amino acid substitutions per site).

The model requires two parameters, specifying the rate exponents for each type of event and their values can be scaled according to the maximum root-to-tip death of the phylogenetic tree (in our case this value is scaled to 1):

- Extinction/Death/Loss Rate (Er): losses of a locus/state
- Recombination Rate (Rr: gains of a locus/state from one species lineage to another

These rates can be thought of as a summary of the evolutionary selective pressures acting to maintain a locus/state or its selective advantage helping to propagate it. Note that these rates remain constant throughout the evolutionary history of the species tree, however in reality they are likely to vary under different population bottlenecks or changing environments.

Using these rates the poisson interarrival time distribution for any event (loss or transfer) can be calculated using the exponential distribution with the rate parameter equal to the sum of the extinction and recombination rates: P(ti) = (Er + Rr) * exp(-(Er + Rr)*ti), where exp is the exponential function and ti is the given inter-arrival time between successive events.

Another important feature of this model is that probabilities of events occurring at a given inter-arrival time:

- Extinction Probability: Er / (Er + Rr)
- Recombination Probability: Rr / (Er + Rr)

In our modelling procedure, first the emergence time of the trait is randomly drawn from a uniform distribution and then randomly assigned to a species lineage existing at that time (trait origination event). Next, the occurrence of events is modelled using a poisson process by randomly drawing a sequence of interarrival times from the inverse cumulative probability function of the exponential distribution: e_t = -log(1-P(i))/(Er + Rr + Lr)), where i is a random variable sampled from a uniform distribution taking values from [0–1].

The inter-arrival time sequence is cumulatively summed and then added to the emergence time of the first event (cumsum(e_t) + birth_time) which gives a sequence of the the event occurrence times to be randomly mapped upon the species tree lineages. Note, only those event occurrence times time of emergence for the locus/state until the time when the species tips are observed (= 1) are considered.

Iterating successively through the event timings, a number betweeen [0–1] is randomly drawn from a uniform distribution (prob_event) and used to determine whether the given event is a loss or recombination:

- Extinction/Loss: if prob_event <= Er / (Er + Rr); otherwise
- Recombination: prob_event > Er / (Er + Rr)

In addition, a locus/state longevity rate (Lr) parameter can be incorporated, which will result in the inter-arrival event distribution of exp(-(L + Er + Rr)), but also add the possibility of no events occurring in the evolution of the trait:

- No Event (Trait State Maintained): if prob_event <= Lr / (Lr + Er + Rr); otherwise
- Extinction/Loss: prob_event <= Er / (Lr + Er + Rr); otherwise
- Recombination: prob_event <= Rr / (Lr + Er + Rr)

If the event is a trait loss, species lineages possessing the locus/state at the given time are extracted and assigned as a loss event, and all descendant lineages occurring after the event time are also assigned as losses. If the event is a recombination, then a species lineage lacking the locus/state at the given time (if it exists) is randomly selected and assigned as a locus gain event, and its descendants are assigned as a locus gain event, in distinction to the initial locus/state origination event. In effect this generates a locus/state distribution presence/absences for the tips of the species tree, as well as the ancestral evolutionary histories of these traits. There may also be tips which never possessed the locus in their evolutionary history (absence). Using this approach we can examine how frequently locus/state distributions overlap between different evolutionary regimes, e.g. loss dominated vs. recombination dominated.

### *Pseudomonas syringae* type III effector family analysis

Data for *P. syringae* effectors including NCBI genomic accession numbers for 494 *P. syringae* strains used for effector identification, associated strain metadata, and classified effector family sequences originates from a previously published study [15], and can be downloaded from the supplementary material provided therein. Genomic assemblies used were generated using the protocol outlined in [15]. Annotation of genome assemblies was performed using prokka (version 1.14.16) [41], pangenome analysis and core-genome nucleotide alignment was produced using PIRATE (version 1.0.4) [42], from which a core-genome phylogenetic tree was generated using IQ-TREE (version 1.6.12) [43]. All software was run in linux. Generation of effector presence/absence matrices, RecPD analyses, and phylogenetic tree visualization and annotation were performed in R.

## Supporting information

RecPD R code

RecPD Tutorial

Evolved Trait Histories

## ACKNOWLEDGEMENTS

We would like to thank all members of the Guttman and Desveaux labs for their insight input to this project. The work was support by a Natural Sciences & Engineering Research Council Discovery Grant to D.S.G.

## Author Contributions

CBD, conceptualization, data curation, formal analysis, methodology, software, validation, visualization, writing, editing DD, conceptualization, review, editing DSG, conceptualization, funding acquisition, methodology, project administration, resources, supervision, writing, review, editing.

## SUPPORTING INFORMATION

**S1 Fig.**
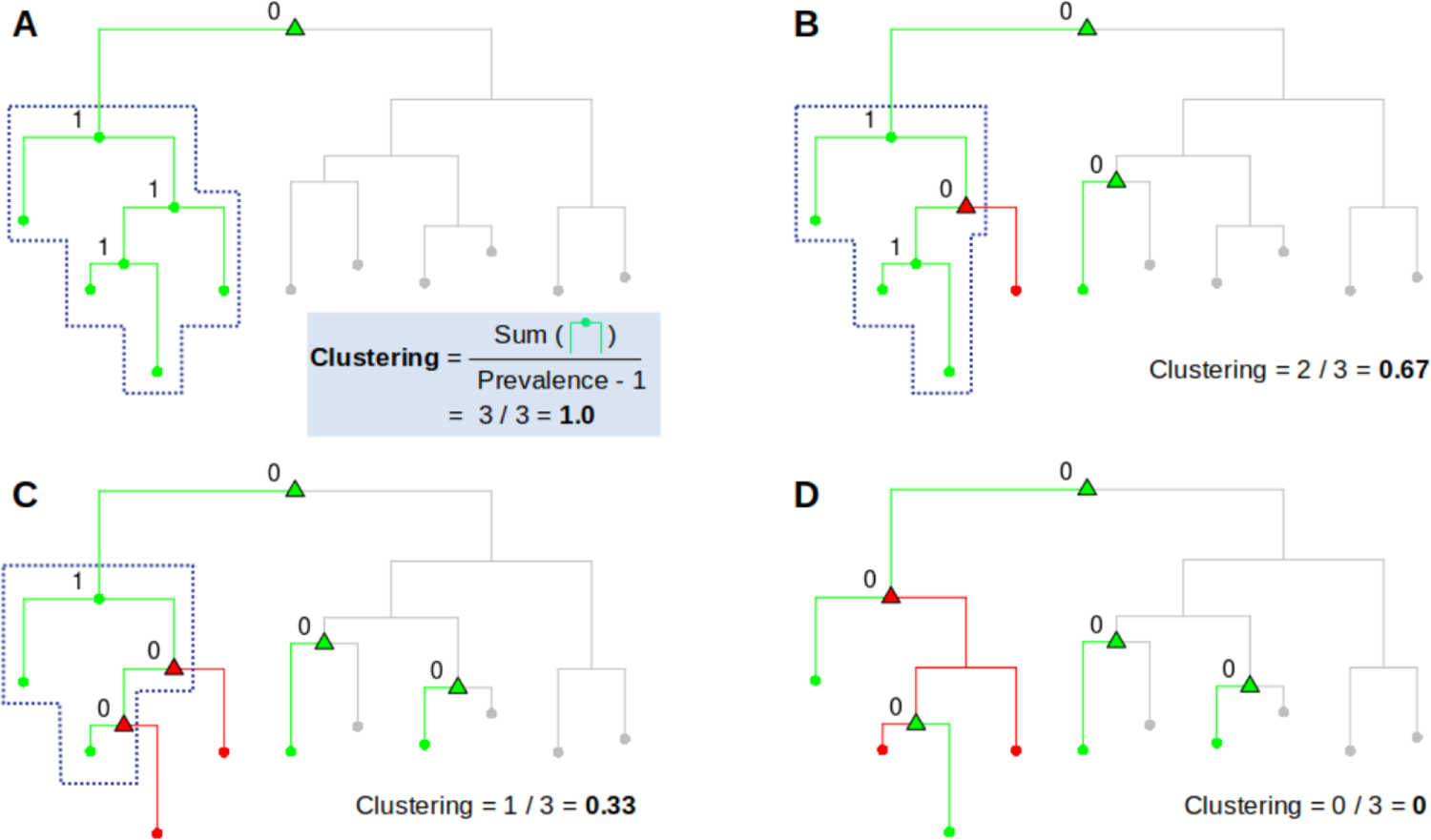
Illustration of RecPD Clustering metric calculation with example distributions. Clustering is calculated from the sum of the number of internal presence state nodes identified, normalized by the maximum clustering possible based on tip prevalence.

**S2 Fig.**
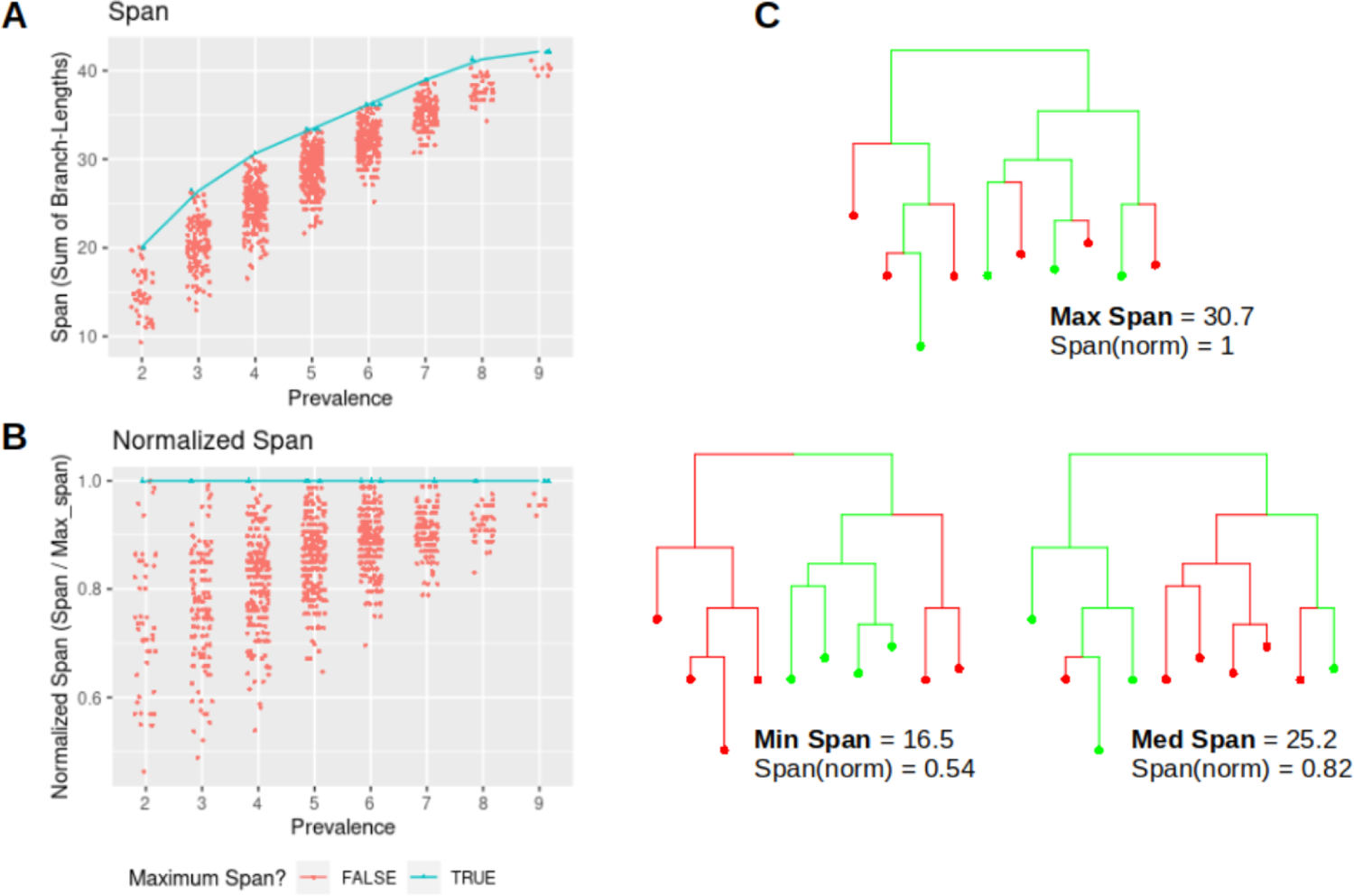
Illustration of RecPD Span metric calculation with example distributions. (A) Span is calculated by summing of branch-lengths joining tips in the phylogenetic tree possessing a gene family with a given level of prevalence. (B) Normalizing by the maximum possible sum of branch-lengths found at the same level of prevalence. (C) Example gene family distributions of prevalence = 4 mapped onto a tree of 10 tips, with maximum, minimum and medium span values.

**S3 Fig.**
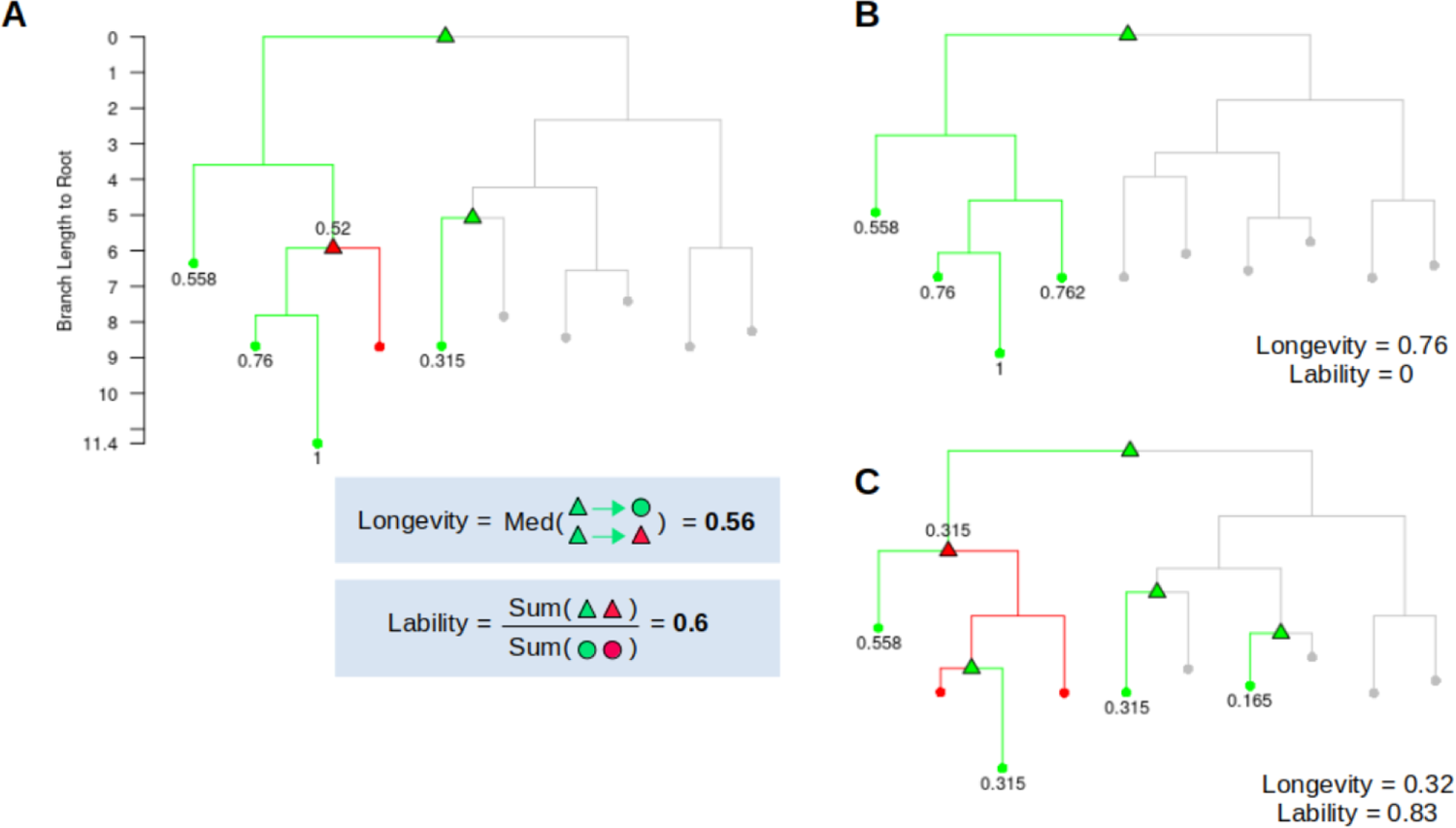
Illustration of RecPD Longevity and Lability metric calculation with example distributions. Longevity is calculated as the median branch-lengths of ancestral gain to loss internal nodes and presence state tips, normalized by the maximum root-to-tip distance of the phylogenetic tree. Lability is the corresponding sum of ancestral gain and loss nodes identified for each RecPD reconstructed gene-family lineage divided by the total number of gained and lost tips. Panels A – C show gene-family distributions of prevalence = 4 mapped onto a tree of 10 tips having approximately equal Longevity and Lability (A), High Longevity and Low Lability (B) and low Longevity and High Lability (C).

**S4 Fig.**
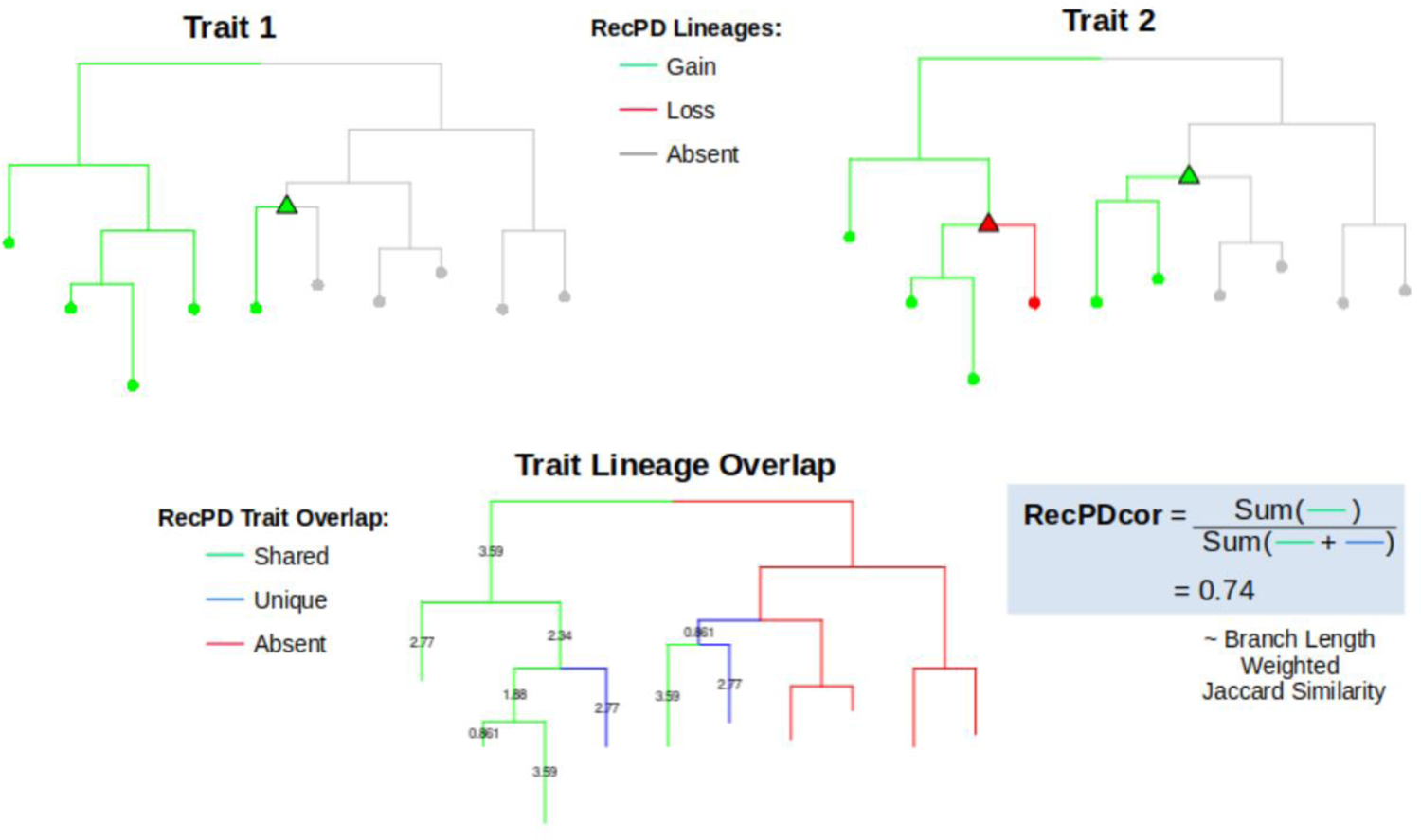
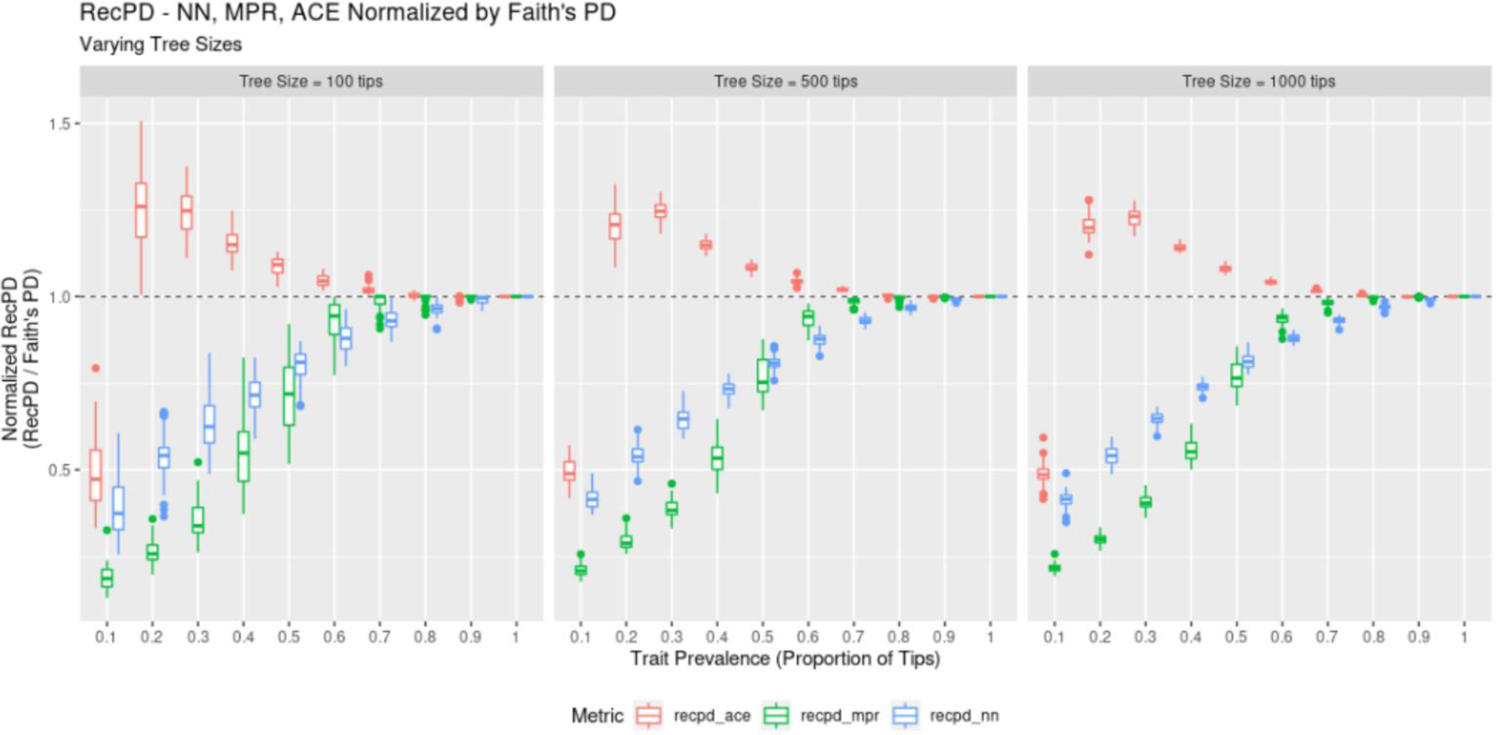
Illustration of RecPDcor metric calculation with example distributions. A pair of RecPD gene family reconstructions are merged into a consolidated phylogenetic tree with mutually present (green), unique (blue), and mutually absent (red) ancestral branches identified. RecPDcor is then calculated as the sum of branch-lengths mutually present branches divided by the total sum of mutually present and unique branches.

**S5 Fig.**
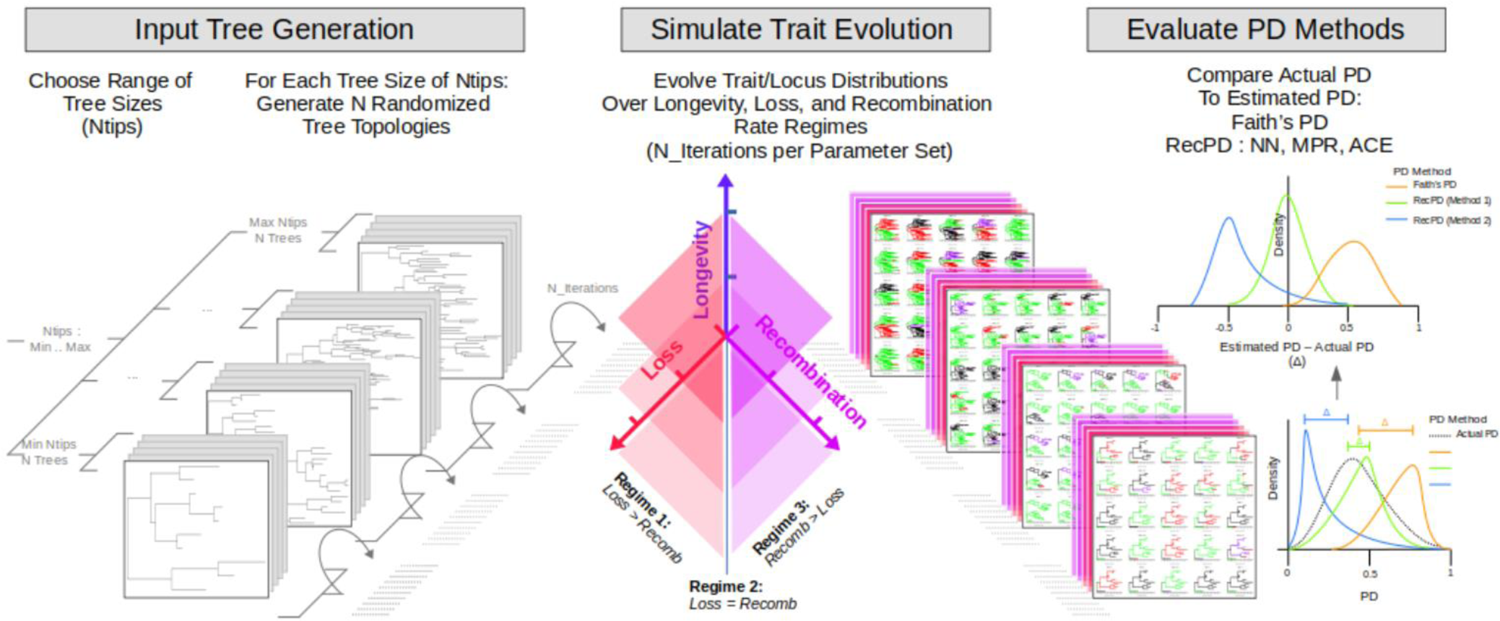
RecPD using the NN, MPR, and ACE ancestral reconstruction approaches compared to Faith’s PD: random gene family distributions for trees of 100, 500, and 1000 tips. 50 random gene-family distributions were generated at each level of prevalence indicated, resulting to 451 distributions in total for each tree.

**S6 Fig.**
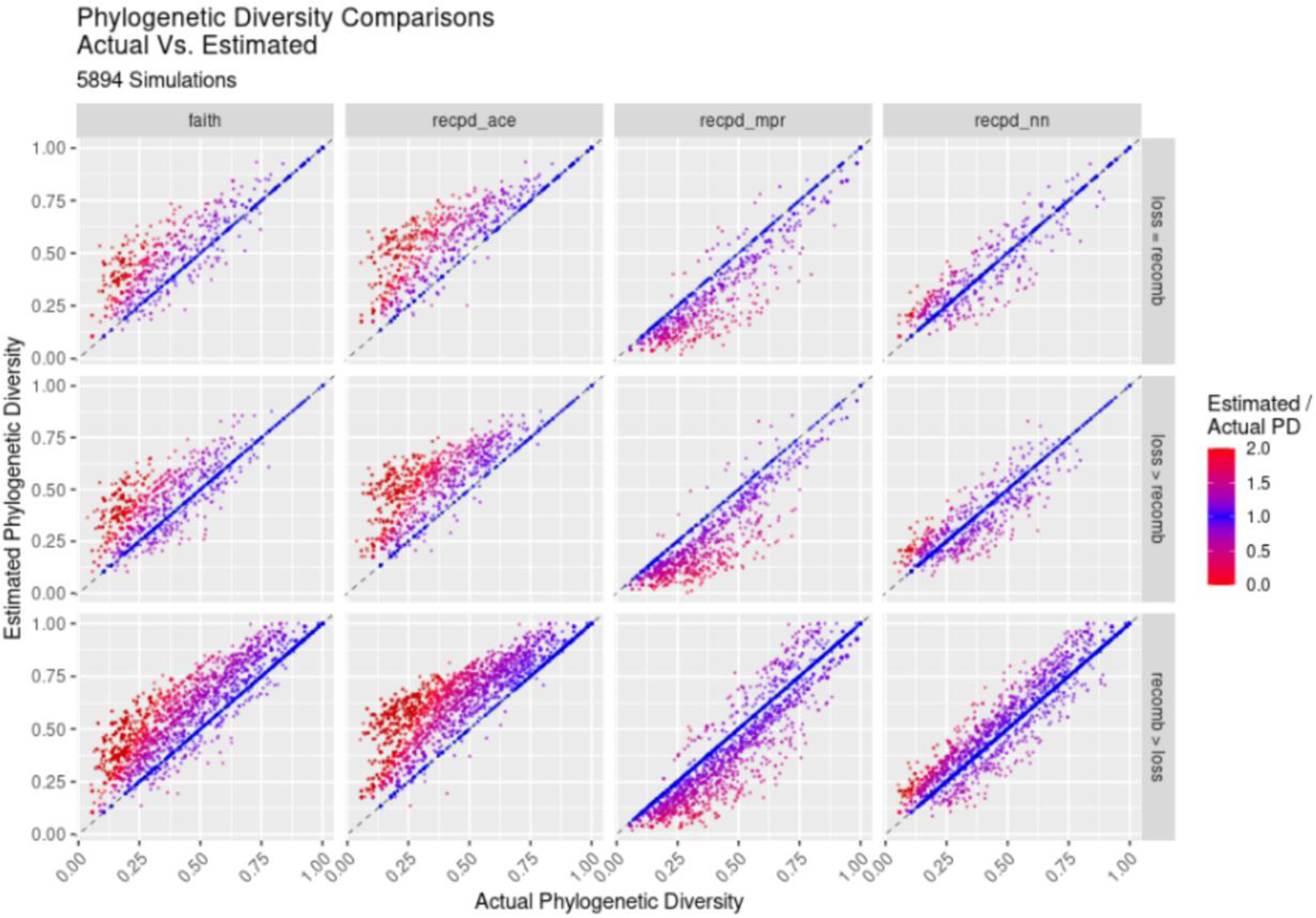
Gene family evolutionary history simulation - outline of simulation experiment protocol.

**S7 Fig.**
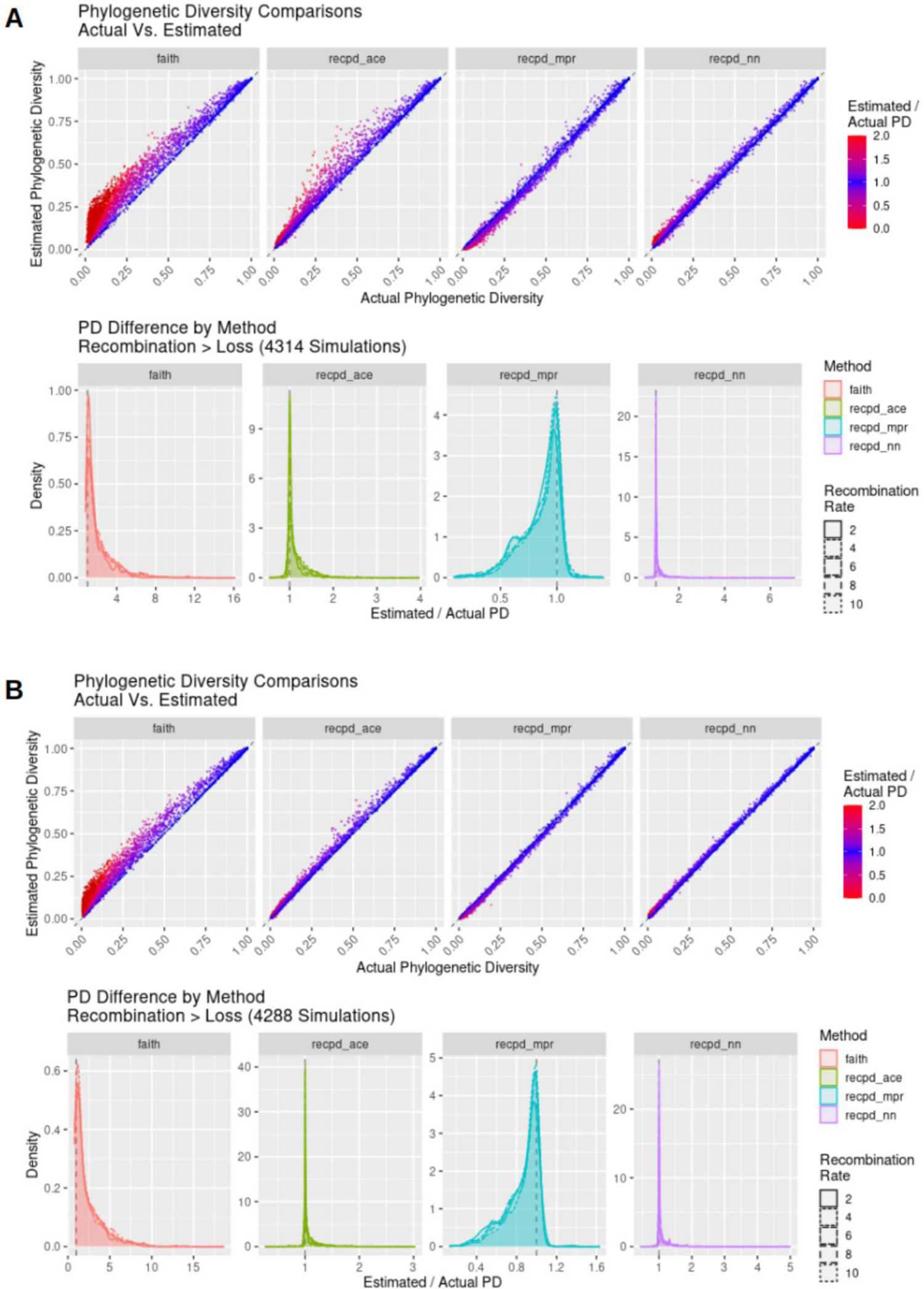
RecPD NN, MPR, and ACE vs. Faith’s PD – estimated / actual PD for evolved gene family distributions by rate regime.

**S8 Fig.**
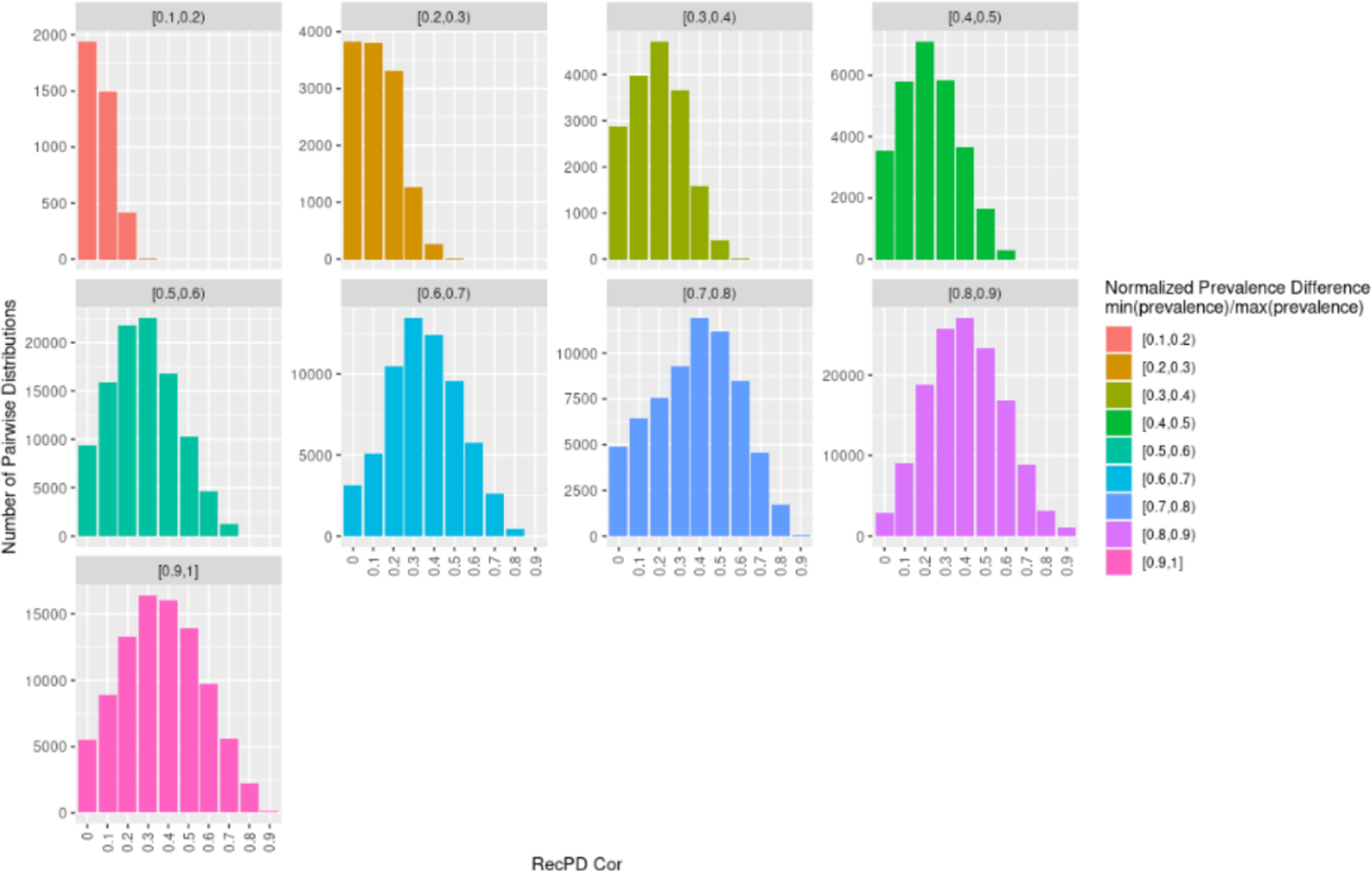
RecPD NN, MPR, and ACE vs. Faith’s PD – estimated / actual PD for evolved gene family distributions by rate regime. (A) Trees with 50 tips. (B) Trees with 100 tips. **S9 Fig.** RecPDcor by prevalence difference. Facets represent the distribution of RecPDcor values binned by the normalized prevalence differences, min(prevalence) / max(prevalence), of each pairwise randomized gene-family distribution comparison compared. Note, normalized prevalence difference = 1 indicate distributions with identical prevalence. Results correspond to a test-case of all possible 1022 gene-family distributions mapped onto a tree of 10 tips.

---

S1 File. R code for calculation of RecPD and associated metrics and tutorial.

S2 File. R code for gene family evolutionary history simulations.

