## Supplementary material for "RecPD: A Recombination-Aware Measure of Phylogenetic Diversity": RecPD R code: recpd_functions.R

```
#recpd_functions.R - all the code for implementing recpd analyses.
#*Requires subfunctions from the 'ape' function to access and process the phylogenetic tree.


#recpd_calc() - main wrapper function for performing RecPD calculations and branch state identification,
#and other cladistic and evolutionary distance based metrics:
recpd_calc <- function(tree, tab, option=c('nn', 'mpr', 'ace'), params=list(nn_select=c('min', 'median'), model_type=c('ER', 'ARD'), kappa=1, lik_cut=0.99), calc=FALSE){
  #Input arguments:
  
  #tree - the species tree
  #tab - a presence absence matrix in dataframe format: rows are a given trait/locus distribution, columns are the corresponding species names matching the tip labels of the tree
  ##Note: all species in the presence/absence matrix (t) should be in the tree
  
  #eff - the locus/state distribution to calculate RecPD
  #node_sp - the list of nearest neighbour descendant tips for each node in the tree
  ##Note: node_sp will be moved into the branch_select_nn() function, and the function will be reconfigured to iterate through all rows of the presence / absence matrix, except for a specific row at a time (eff will also be removed).
  
  #method - the method for assigning internal node states - 3 options: 
  #1) 'nn' : nearest-neighbours approach (by default)
  #2) 'mpr' : Most Parsimonious Reconstruction (using the MPR() function from the ape package)
  #3) 'ace' : Maximum Likelihood Ancestral Character Estimation (using the ace() function from the ape package).
  #Note that the state ancestral likelihood measurements (marginal likelihoods) are known as empirical bayesian posterior probabilities (see: https://lukejharmon.github.io/ilhabela/instruction/2015/07/03/ancestral-states-2/)
  #4) a bayesian approach? anc.Bayes() from the phytools package? This appears to assume that states are continuous??
  
 
  #Additional parameters are supplied using the params list:
  #If option = 'nn':
  #nn_select = choice of evolutionary distance cutoff to select nearest-neighbour nodes - 'min' for minimum pairwise distance, 'median' for median pairwise distance.
  ##If option = 'ace':
  #model_type -  can choose between rates equal ('ER') or all rates different ('ARD') models.
  #kappa - exponent for transforming tree branch-lengths.
  
  #calc - TRUE/FALSE if you want to perform topology or evolutionary metric calculations.
  
  #If calc == TRUE, get the maximum spanning distributions here, to prevent rerunning multiple times 
  #for multiple trait distributions - will waste time, especially for larger species trees.
  if(calc == TRUE) max_span <- get_max_span(tree)
  
  
  #Output list:
  res <- list()
  
  #Make sure that the presence/absence tab is in the proper format, i.e. a matrix with row indices:
  #Remember to add row and column names accordingly. Even if Tab is just a named array, make sure the names are turned into column names.
  if(is.null(nrow(tab))){
    tab <- matrix(tab, nrow=1, ncol=length(tab), dimnames=list(make.names(1), names(tab)))
  }
  
  for(i in 1:nrow(tab)){
    #Make sure tips in the presence absence matrix are ordered according to the tip ordering in the phylogenetic tree:
    #Names of columns in t must match the tree!
    #Have to have some error checking to make sure that names match...
    #Resulting array should have strain names....
    t <- tab[i,tree$tip.label]
    
    #Initialize an array to store the states of terminal tip and internal nodes
    node_state <- NULL
    
    #Assign states based on nearest neighbour descendant tips of a given node
    
    #Option 1: Nearest neighbour descendants:
    if(option[1] == 'nn'){
      #Only generate the species-tree interal-node nearest-neighbour tip list once! 
      if(i == 1){
        #Check if any input parameters have been changed:
        nn_select <- 'min'
        if(!is.null(params$nn_select[1])) nn_select <- params$nn_select[1]
        
        node_sp <- nn_nodes(tree, option=nn_select)
      }
      node_state <- node_state_nn(tree, t, node_sp)
      
    }
    #Option 2: Most Parsimonious Reconstruction (from ape package)
    else if(option[1] == 'mpr'){
      node_state <- node_state_mpr(tree, t)
      
    }
    #Option 3: Maximum Likelihood Ancestral Character Estimation *from ape package)
    else if(option[1] == 'ace'){
      
      #Update input parameters for ace, if provided
      if(i == 1){
        #defaul parameters:
        model_type <- 'ER'
        kappa <- 1
        lik_cut <- 0.99
        
        #Check if any of the model paramaters have been changed:
        if(!is.null(params$model_type[1])) model_type <- params$model_type[1]
        if(!is.null(params$kappa)) kappa <- params$kappa
        if(!is.null(params$lik_cut)) lik_cut <- params$lik_cut
      }
      
      #ace will only work if there is at least one presence or absence state on the tips:
      if(sum(t) < length(t)){
        
        node_state <- node_state_ace(tree, t, model_type, kappa, lik_cut)
      }
      else{
        node_state <- array(rep(1, Ntip(tree)+Nnode(tree)), dimnames=list(1:(Ntip(tree) + Nnode(tree))))
      }
    }
    
    #Assign branch states based on ancestral node state reconstruction:
    branch_state <- branch_state_assign(tree, node_state)
    
    #Merge descendant and ancestral node states together to identify putative gain and loss nodes:
    #also output all of the updated state annotations (gain, loss, absence)
    #for tips, branches, and internal nodes, for both plotting and additional calculations.
    node_state_merge <- node_ident5(tree, node_state, branch_state)
    
    pd <- sum(tree$edge.length*branch_state)/sum(tree$edge.length)
    
    
    ##Perform other calculations?:
    
    span <- NULL
    cluster <- list(NULL, NULL)
    evo <- list(NULL, NULL, NULL, NULL)
    
    if(calc){
      
      #Topology:
      ##Span:
      span <- span_calc(tree, t, max_span)
      
      ##Clustering (has to take the entire set of annotated internal node states):
      cluster <- cluster_calc(tree, t, node_state, branch_state)
      
      #Evolutionary:
      ##Longevity, lability, gain and loss times.
      evo <- longevity_calc(tree, node_state_merge)
    }
    
    res[[rownames(tab)[i]]] <- list('metrics' = list(recpd=pd, 
                     span=span, 
                     cluster=cluster[[2]],
                     # cluster_n=cluster[[1]],
                     # cluster_b=cluster[[2]],
                     longevity=median(evo$longevity), #Note, longevity is taken as the median of trait lineage branches, leading from gain nodes to both presence state tips and descendant loss nodes (if present). 
                     lability=evo$lability3 #,
                     # lability1=evo$lability1,
                     # lability2=evo$lability2,
                     # lability3=evo$lability3,
                     # gain_times=evo$gain_times, 
                     # loss_times=evo$loss_times
                     ), 
                'node_state' = node_state_merge, 
                'ns_old' = list(tip_state=node_state[(1:Ntip(tree))],
                            branch_state=branch_state,
                            node_state=node_state[-(1:Ntip(tree))]))
  }
  return(res)
}

#branch_state_assign() - branch state annotation function based on the ancestral state reconstruction
#of neighbouring internal nodes of the species phylogenetic tree.

branch_state_assign <- function(tree, node_state){
  branch_state <- NULL
  
  for(j in 1:nrow(tree$edge)){
    e <- tree$edge[j,]
    e1 <- e[1]
    e2 <- e[2]
    
    s1 <- node_state[e1]
    s2 <- node_state[e2]
    
    #If the branch ancestor node is the root, only include it if its node state is present (=1)?
    #Call branches present if ancestor and child nodes are both assigned as presence, split, or a combination of presence/split:
    #Call branches absent if ancestor and child nodes are both assigned as absence, split->absence, or absence->split.
    
    #Only keep branches leading to split nodes if its parent is a gain, and vice versa?
    
    if(s1 != 2 & s1 == s2){
      branch_state <- append(branch_state, as.numeric(s1))
    }
    # else if(s1 != 2 && s2 == 2){
    #   branch_state <- append(branch_state, 1)
    # }
    else if(s1 != 0 && s2 != 0){
      branch_state <- append(branch_state, 1)
    }
    else{
      branch_state <- append(branch_state, 0)
    }
  }
  
  return(branch_state)
}


#node_ident5() - consolidation of previously generated phylogenetic tree ancestral state node state assignments;
#Assigns split state nodes to gain or loss events, if they correspond to ancestor - descendant branch state changes, 
#branch state ancestor to branch state descendant:
#gain node = absent to present,
#loss node = present to absent.

#Also merge any loss or gain nodes of idential state which occurr within the same phylogenetic tree lineages.

node_ident5 <- function(tree, node_state, branch_state){
  #Get the tip states:
  tip_state <- node_state[1:Ntip(tree)]
  
  #Get the internal node states:
  node_state <- node_state[-(1:Ntip(tree))]
  
  #Get the edge matrix of the tree:
  edge <- tree$edge
  
  
  #Identify state change nodes based on changes of ancestral and descendant branches of each node.
  #Go through the edge matrix, for each parent node, get its ancestral edge, 
  #and check if the branch states change, if they do then keep that node as a state change.
  
  #Define an array for storing updated node states:
  ns_merge <- array(rep(NA, Nnode(tree)), dimnames=list((Ntip(tree) + 1):(Ntip(tree) + Nnode(tree))))
  
  
  for(i in 1:nrow(edge)){
    #Get parent and child nodes for each edge:
    p <- edge[i,1]
    c <- edge[i,2]
    #Exclude the root:
    if(p != Ntip(tree) + 1){
      #Get the ancestral node:
      a <- Ancestors(tree, p, 'parent')
      
      #Find the ancestral edge:
      j <- which(edge[,1] == a & edge[,2] == p)
      
      #Compare branch states of the ancestral and decendant branches of the parent node:
      
      #If the state changes, then assign the node state to:
      #Node = Gain if ancestral branch is absent (0), and descendent branch is present (1) = gain lingeage branches
      #Node = Loss if ancestral branch is present (1), and descendent branch is absent (0) = loss lineage branches
      
      if(branch_state[i] == 1 & branch_state[j] == 0) ns_merge[as.character(p)] <- 1
      if(branch_state[i] == 0 & branch_state[j] == 1) ns_merge[as.character(p)] <- 0
    }
  }
  
  #Keep only the nodes where states change:
  rm <- which(is.na(ns_merge))
  ns_merge <- ns_merge[-rm]
  
  #Also check if the root node is assigned to an absence state. Then check descendant nodepaths from the root which 
  #are all absent and assign to -1 state, i.e. absent lineages.
  
  r <- which(edge[,1] == Ntip(tree) + 1)

  if(branch_state[r[1]] == 0 | branch_state[r[2]] == 0){
    #Get all nodepaths:
    np <- nodepath(tree)
    
    #Traverse the nodepaths from the root, and identify any nodes on absent lineages, assign them to -1:
    bs_abs <- list()
    
    for(i in 1:length(np)){
      x <- np[[i]]
      #Get the edge indices for consecutive nodes in each nodepath:
      j <- which(edge[,1] %in% x[-length(x)] & edge[,2] %in% x[-1])
      
      bs <- branch_state[j]
      #Find the consecutive absent state edge indices, 
      #i.e. where does the branch state change to a gain?
      k <- which(bs[-length(bs)] != bs[-1])
      
      if(bs[1] == 0){
        if(length(k) != 0){
          bs_abs[[i]] <- j[1:k[1]]
        }
        else{
          bs_abs[[i]] <- j
        }
      }
      
      #Also assign any tips to absent states if they appear on a absent lineage
      #Descended directly from the root:
      if(bs[length(j)] == 0 & length(k) == 0){
        tip_state[i] <- -1
      }
    }
    
    #Assign the identified branches to absent state:
    branch_state[unique(unlist(bs_abs))] <- -1
  }
  
  #Add the root node if at least one descendant branch has been assigned as presence:
  r_d <- which(edge[,1] == Ntip(tree) + 1)
  
  if(length(which(branch_state[r_d] == 1)) != 0){
    ns_merge <- append(array(1, dimnames=list(Ntip(tree) + 1)), ns_merge)
  }
  
  return(list(tip_state=tip_state, 
              branch_state=branch_state,
              node_state=ns_merge)
         )
}


##nn_nodes() - Identify Nearest Neighbour Tips Descended From Each Internal Node of the Phylogenetic Tree.
#The Preliminary Step For Nearest-Neighbours Ancestral State Reconstruction (required by the node_state_nn()).

#For each internal node of the tree, identifying the subset of tips of each child-node descendant,
#then ompare the pairwise phylogenetic distances between each tip subset, and select those tips which are:
#1) Equal to the minimum pairwise distance, or 2) <= median pairwise distance.

nn_nodes <- function(tree, option=c('min', 'median')){
  #Get the pairwise tip distance matrix
  #dist <- cophenetic.phylo(tree)
  
  #Get the distances between tips and nodes:
  dist <- dist.nodes(tree)
  
  #Get internal node indices
  nodes <- Ntip(tree) + seq(1, Nnode(tree))
  node_sp <- list()
  
  for(n in nodes){
    #Get descendant children nodes for each internal node
    descend <- Descendants(tree, n, type='children')
    
    i <- which(descend <= Ntip(tree))
    
    #Check if descendant nodes are tips:
    #If nodes include 2 tips:
    if(length(i) > 1){
      
      t1 <- descend[1]
      t2 <- descend[2]
      
      close_t <- descend 
      
    }
    #If only one descendant is a tip, get the descendant tips of the other child internal node
    else if(length(i) == 1){
      #If nodes include 1 tip:
      t1 <- descend[i]
      
      #Get the descendant tips of the other child descendant node:
      t2 <- unlist(Descendants(tree, descend[-i], type='tips'))
      
      #Get their pairwise evolutionary distances:
      d_sub <- dist[t1, t2]
      
      #Find the closest tip to the child descendant node:
      if(option[1] == 'min'){
        close_t <- c(t1, as.numeric(names(d_sub)[which(d_sub == min(d_sub))]))
      }
      #Or, find the pairs of tips <= the 25%-tile (?median) pairwise distance:
      else{
        close_t <- c(t1, as.numeric(names(d_sub)[which(d_sub <= median(d_sub))]))
      }
      
    }
    #If both descendant nodes are internal-nodes, get the descendant tips for each:
    else{
      #If nodes are internal, find the nearest neighbour tips between the descendant nodes:
      t1 <- unlist(Descendants(tree, descend[1], type='tips'))
      t2 <- unlist(Descendants(tree, descend[2], type='tips'))
      
      #Get their parwise evolutionary distances:
      d_sub <- dist[t1, t2]
      
      #Take the closest related tips:
      if(1){
        
        #Find the closest tip pair to the child descendant node:
        if(option[1] == 'min'){
          m_ind <- which(d_sub == min(d_sub), arr.ind=TRUE)
          #close_t <- as.numeric(c(rownames(d_sub)[m_ind[1,1]], colnames(d_sub)[m_ind[1,2]]))
        }
        #Or, find the pairs of tips <= the 25%-tile (?median) pairwise distance:
        else{
          m_ind <- which(d_sub < median(d_sub), arr.ind=TRUE)
          #close_t <- as.numeric(c(rownames(d_sub)[unique(m_ind[,1])], colnames(d_sub)[unique(m_ind[,2])]))
          
        }
        close_t <- as.numeric(c(rownames(d_sub)[unique(m_ind[,1])], colnames(d_sub)[unique(m_ind[,2])]))
      }
      else{
        #Or choose the closest related pairs of strains between descendant nodes.
        m_ind <- apply(d_sub, 1, function(x) which(x == min(x)))
        close_t <- as.numeric(c(rownames(d_sub)), unique(as.numeric(colnames(d_sub)[m_ind])))
      }
    }
    
    #Order the pair of tips based on increasing distance to the given node:
    #Sort tips by increasing distance to their parent node:
    d <- dist[n, close_t]
    
    #Store closest related descendant species of a node:
    node_sp[[as.character(n)]] <- tree$tip.label[close_t[order(d)]] #list(close_t, tree$tip.label[close_t[order(d)]], sort(d))
  }
  
  return(node_sp)
}


#Ancestral Reconstruction Functions of Internal Node Trait States,
#Used For Branch Selection to Calculate RecPD and Associated Metrics:

#node_state_nn() - The Default / Original Implemenatation Based on Nearest-Neighbours:
node_state_nn <- function(tree, t, node_sp){
  
  #Define a function to assign states to internal nodes:
  assign_state <- function(j, node_sp, presabs){
    close_sp <- as.character(unlist(node_sp[j]))
    
    #Assign split states based on the states of nearest neighbour descendant nodes:
    ##Version 1: look at the closest related pair of strains descended from each node,
    #ensuring that they come from different child nodes. If they both have the same state, then
    #assign the parent node to that state, otherwise annotate the parent node as a split (potential gain/loss event).
    #Version 2: only look at the first, closest descendant of the parent node.
    s <- sum(presabs[1,close_sp])

    state <- 0
    #If all tips are present state:
    if(s == length(close_sp)){
      state <- 1
    }
    else if(s != 0){
      state <- 2
    }
    
    return(state)
  }
  
  #Presence / absence state of locus/trait
  presabs <- ifelse(t != 0, 1, 0)
  
  #return(presabs)
  
  #An array to store the states of internal nodes and tips
  node_state <- NULL
  
  n <- 1:(Ntip(tree) + Nnode(tree))
  
  for(i in n){
    if(i <= Ntip(tree)){
      node_state <- append(node_state, presabs[1,tree$tip.label[i]])
    }
    else{
      node_state <- append(node_state, assign_state(as.character(i), node_sp, presabs))
    }
  }
  names(node_state) <- n
  return(node_state)
  
}


#node_state_mpr() - Ancestral Node State Reconstruction Using Most Parsimonious Reconstruction:
#*requires the MPR() function from the 'ape' package.
node_state_mpr <- function(tree, t){
  #Presence / absence state of locus/trait
  c <- ifelse(t[tree$tip.label] != 0, 1, 0)
  names(c) <- 1:Ntip(tree)
  
  #Add the internal node labels to the tree, replacing 
  
  
  #Use the Most Parsimonious Reconstruction (MPR) function in the ape package to assign ancestral states for a given T3SE to all nodes in the tree.
  #The MPR algorithm will produce a matrix of nodes (in their order of appearance in the tree?) and their corresponding lower and upper value character state prediction for each node. 
  #Therefore, in the case of a binary discrete character, e.g. presence = 1, absence = 0:
  #- Nodes with [1,1] indicate that the ancestral state of the is predicted to be present.
  #- Nodes with [0,0] indicate the ancestral state of the trait is predicted to be absent.
  #- Nodes with [0,1] indicate an ambiguous ancestral state.
  
  #Note: the tree must be unrooted.
  
  #The states assigned according to the appearance of parent nodes in the edge matrix of the unrooted tree:
  #Note, have to supply an unrooted tree but also designate a tip to use as a root. Given that midpoint rooted and chronograms (usually) do not have a tip to designate as a root, so the resulting node-state mappings have to be remapped to the original tree to be visualized properly.
  #To make this process easier by preserving the original clade structuring of the tree, employ the following trick:
  
  #Replace node labels (usually bootstrap supports) with node IDs,
  #So output from mpr() is intelligible.
  tree$node.label <- (Ntip(tree)+1):(Ntip(tree) + Nnode(tree))
  
  #Add an artificial outgroup, attaching it to the root node of the original tree, then root the new tree on it, which will preserve the :
  r_tree <- bind.tip(tree, 'outgroup', edge.length=1)
  r_tree <- root(r_tree, 'outgroup')
  
  r <- 'outgroup'
  
  #Make all branch lengths the same length.
  r_tree$edge.length <- rep(1, length(r_tree$edge.length))
  
  #Add the outgroup to the original tip presence/absence state array:
  t2 <- c(t, `outgroup`=0)
  
  #Now perform MPR:
  parsim <- MPR(ifelse(t2[r_tree$tip.label] != 0, 1, 0), unroot(r_tree), r)
  
  #Assign node labels to the node parsimony state array according to their appearance in the rooted tree (note: in some imported trees, the original node labels assigned by MPR are the node bootstrap support values, but generally they will be in the order of the node numbers of the tree).
  
  #Convert the table into an array, assigning states based on correspondence of lower and upper state predictions.
  node_state <- NULL
  node_state <- apply(parsim, 1, function(x) if(x[1] == x[2]){return(x[1])}else{return(2)})
  
  #Assign internal node IDs to the node_state array
  names(node_state) <- rownames(parsim)
  
  #Also append the presence/absence states of the tips:
  node_state <- append(c, node_state)
  
  #Check if there are any presence state tips without ancestral nodes
  #annotated: will occur for trait distributions of low prevalence, 
  #or if the trait appears in a clade where states are largely absent.
  #If there are, assign as a split (= 2).
  
  a <- Ancestors(tree, which(c == 1), type='parent')
  
  i <- which(node_state[as.character(a)] == 0)
  
  if(length(i) != 0) node_state[as.character(a)[i]] <- 2
  
  #return(node_state)
  
  # #The only apparent difference between the edge matrices of the unrooted trees used for MPR and the original tree is that the original root node will be removed and set to a new node leading to the outgroup and its nearest descendant clades.
  # 
  # #To map internal nodes of the rooted tree to the original tree, we have to remove the first and last edges in the former, which will be the branch leading from the new root to the old, and from it to the artifically added outgroup, respectively.
  # 
  # #Make an array to map rooted to original tree internal nodes:
  # node_lab <- array(unique(tree$edge[,1]), dimnames=list(unique(r_tree$edge[c(-1,-nrow(r_tree$edge)),1])))
  # 
  # #The first node_state from MPR will correspond to the root of the original tree, for which a state is not assigned (set to -1):
  # node_state[1] <- -1
  # 
  # #Relabel the node_state array to match the internal nodes in the original tree:
  # names(node_state) <- node_lab#[names(node_state)]
  # names(node_state)[1] <- Ntip(tree) + 1
  # 
  # 
  # #Also add the tip node presence and absence states
  # node_state <- append(c, node_state)
  # 
  # #Since the Node state of the root is absent (-1), assign it a state (1, 0, 2) based on the states of its descendant nodes:
  # d <- Descendants(tree, Ntip(tree)+1, type='children')
  # 
  # if(node_state[d[1]] == node_state[d[2]]){
  #   node_state[Ntip(tree) + 1] <- node_state[d[1]]
  # }
  # else{
  #   node_state[Ntip(tree) + 1] <- 2
  # }
  # 
  # #Note, for many instances (low prevalence), all internal nodes will be annotated as absent, 
  # #particularly if presence tips do not share a common ancestor, or if the presence-tip is found in a clade 
  # #where the majority of tips lack the state. In these instances where there are no ancestral node assignments for presence tips, 
  # #assign them as splits(?):
  # 
  # a <- unique(Ancestors(r_tree, which(t2 == 1), type = 'parent'))
  # i <- which(node_state[a] == 0)
  # 
  # if(length(i) != 0){
  #   node_state[a[i]] <- 2
  # }
  
  # if(sum(node_state[(Ntip(tree)+1):length(node_state)]) == 0 & sum(node_state[1:Ntip(tree)]) != 0){
  #   a <- Ancestors(r_tr, which(t2 == 1), type = 'parent')
  #   
  #   node_state[a] <- 2
  # }
  
  return(node_state)
}


#node_state_ace() -  Ancestral Node State Reconstruction Using ML (Ancestral Character Estimation):
#*Requires the ace() function from the 'ape' package.
node_state_ace <- function(tree, t, model_type='ER', kappa=1, lik_cut=0.9){
  #lik_cut - a cutoff for selecting states based on the scaled state likelihood of each node. 
  #This will effect the corresponding node state assignments, if it is too high, all nodes will be splits.
  
  #Presence / absence state of locus/trait
  c <- ifelse(t[tree$tip.label] != 0, 1, 0)
  names(c) <- 1:Ntip(tree)
  
  #ace will throw an error if there are 0 or negative length branches.
  #Convert all of these edges to a minimum edge length. Let's say 1e-10
  if(length(which(tree$edge.length == 0)) != 0){
    tree$edge.length[which(tree$edge.length == 0)] <- 1e-9
  }
  
  #Results will be quite different if branch lengths are all set to 1, and RecPD appears to be close to the other approaches
  #tree$edge.length <- rep(1, length(tree$edge.length))
  
  #Equal Rates assumed for state transitions?
  if(model_type == 'ER'){
    ace_res <- ace(ifelse(t[tree$tip.label] != 0, 1, 0), tree, type='discrete', method="ML", model='ER', marginal=TRUE, kappa=kappa)
  }
  else if(model_type == 'ARD'){
    #Unequal Rates?
    #The 'model' argument allows you to specify the number of different rate categories (integer) to describe transitions between states. Rates are estimated using ML using the best fit-model, and 
    #Note that given the size of the tree and locus/state tip distribution, some rates (e.g. gains) might not be estimated. Changing the scaling of the branch-lengths (kappa) will result in a multiple rates being estimated, although this means that the estimated rates have to be rescaled. This rescaling also changes the resulting likelihood estimates as well.... The intuition is that by raising the branch-lengths of a tree by a certain exponent to approximate speciational change leads to better ancestral character estimates (This applies also to the ER model).
    
    ace_res <- ace(ifelse(t[tree$tip.label] != 0, 1, 0), tree, type='discrete', method="ML", model='ARD', marginal=TRUE, kappa=kappa)
  }
  
  
  #Ancestral state scaled likelihoods:
  anc <- ace_res$lik.anc
  
  #An array for storing node states
  node_state <- NULL
  
  #Assign presence, absence, or split state to nodes if their 
  #corresponding state likelihoods are >= lik_cut, <= 1-lik_cut, or not, respectively.
  for(i in 1:nrow(anc)){
    if(anc[i,1] >= lik_cut){
      node_state <- append(node_state, 0)
    }
    else if(anc[i,1] <= 1-lik_cut){
      node_state <- append(node_state, 1)
    }
    else{
      node_state <- append(node_state, 2)
    }
  }
  
  #Nodes in sequence:
  n_ord <- seq(Ntip(tree)+1, Ntip(tree)+Nnode(tree))
  
  names(node_state) <- n_ord
  
  #Add the tip node states
  node_state <- append(c, node_state)
  
  return(node_state)
}


###
#Additional Functions
###

#Span:
#For a given trait distribution with prevalence (P):
#Recalculate the PD by summing all tree branch lengths between tips possessing a trait and the tree root; 
#Then divide by the sum of branch lengths for the maximum spanning locus distribution with the same P. 

#get_max_span() - Finds The Maximum Spanning Trait Distributions (Based on Their Total Sum of Branch Lengths) For A Given Phylogenetic Tree:
#*Note that span is not adjusted for potential recombination.
get_max_span <- function(tree){
  
  #Start with a full tree (P = Ntip),
  #Remove a single tip and its accompanying ancestral branch, and recalculate the sum of branch lengths.
  #Select the tip which results in the lowest reduction in the sum of branch lengths to find the maximum spanning distribution for P = Ntip -1. 
  #Use this tip distribution to repeat the step above until reaching 2 tips.
  
  np <- nodepath(tree)
  dn <- dist.nodes(tree)
  
  b_tot <- sum(tree$edge.length)
  
  n <- 1:Ntip(tree)
  
  tmp <- np
  
  s_res <- list()
  for(p in (Ntip(tree)-1):2){
    #How much does the total tree branch lengths change when removing the giving tip:
    #Also have to keep track of the previous nodes which have been removed.
    #However, to initialize the span distribution, this could go in two ways:
    #Start from P-1 presence tips, find the single tip which has the smallest distance to the root (only considering distance to ancestral node ignores if the tip is present in a highly divergent clade), and remove it along with its nodepath; use this procedure to iteratively find the next tip to remove.
    #Or take all the nodepaths of every tip, 
    
    #Get the resulting nodepaths remaining in the tree after removing a given tip:
    j <- lapply(1:length(tmp), function(x) unique(unlist(tmp[-x])))
    
    #For each removed tip,
    #use the remaining nodepaths to calculate the total tree branch lengths:
    d <- unlist(
      lapply(j, 
             function(x){
               k <- which(tree$edge[,1] %in% x & tree$edge[,2] %in% x)
               sum(tree$edge.length[k])
             }
      )
    )
    
    #Select the tip to remove which reduces the total tree branch length the least:
    rm <- order(d, decreasing=TRUE)[1]
    
    tmp <- tmp[-rm]
    
    s_res[[as.character(p)]] <- list(max_span=d[rm], tips=sapply(tmp, function(x) rev(x)[1]))
  }
  
  #Return the maximum spanning distribution and sum of branch lengths by prevalence:
  return(s_res)
}


#span_calc() - Calculates The Span For A Given Trait Distribution Found On A Species Phylogeny Based On
#The Sum Of Branch Lengths Joining All the Tree Tips In Which The Trait is Found, Normalized By The 
#Potentially Maximum Spanning Trait Distribution With The Same Prevalence (calculated by get_max_span()).
span_calc <- function(tree, t, max_span){
  #Get the prevalence of the given trait/locus:
  p <- sum(t)
  
  #Get the nodepaths of the tree:
  np <- nodepath(tree)
  
  #Calculate the sum of branch lengths for the given locus distribution
  #Note, presence/absence matrix should be sorted by the tip label order in the tree:
  t <- which(t != 0)
  
  if(length(t) != 0 & p != 1){
    
    n <- unique(unlist(np[t]))
    
    i <- which(tree$edge[,1] %in% n & tree$edge[,2] %in% n)
    
    br_p <- sum(tree$edge.length[i])
    
    return(br_p/max_span[[as.character(p)]]$max_span)
  }
  else{
    if(p == 1){
      return(0)
    }
    else if(p == Ntip(tree)){
      return(1)
    }
  }
}

####
#Clustering:
####

#cluster_calc() - Calculate The Branch-State Clustering Coefficient For A Trait Distribution: 
#For Each Internal Node of the Phylogenetic Tree (Excluding Root) Identified As A Gain State by RecPD:
#Calculate the Proportion of Descendant Branches (or Tips) Which Are Also Annotated As Present State, 
#Then Sum These Together and Normalize by The Maximum Clustering Value Expected For A Trait Distribution With 
#The Same Prevalence.

cluster_calc <- function(tree, t, ns, bs){
  #ns - node_states predicted using a given ancestral state reconstruction method (includes split nodes).
  #bs - branch_states. 
  
  clust_b <- 0
  clust_n <- 0
  
  clust_coeff_n <- 0
  clust_coeff_b <- 0
  
  #Descendant tips of nodes:
  desc_t <- Descendants(tree, type='tips')
  
  #Descendant children of nodes:
  desc_c <- Descendants(tree, type='children')
  
  #Extract Internal Nodes:
  n <- as.numeric(names(ns[-(1:(Ntip(tree)))]))
  
  #First step for calculating clustering: identify potential internal nodes which delimit presence state clades (not gains),
  #And calculate the proportion of presence state tips descended from them. 
  #Any internal nodes not matching this criterion are assigned 0.
  
  #Two ways of selecting presence state clades:
  #1)Based on the proportion of descendant presence state tips from each internal node:
  #2)If the descendant branches of the node are assigned gain/presence states based on ancestor -> descendant node state relationships.
  
  #return(desc_c)
  #return(lapply(desc_t, function(x) mean(t[as.character(x)])))
  
  if(length(n) != 0){

    #Check if the descendant branches of a given node are presence state:
    db <- sapply(n, function(i){
      sum(bs[which(tree$edge[,1] %in% i & tree$edge[,2] %in% desc_c[[i]])])
    })
    
    #Calculate the proportion of presence state tips descended from each node in the gain lineage?
    #clust_b <- sapply(1:length(n), function(i) ifelse(db[i] == 2, sum(t[desc_t[[n[i]]]])/length(desc_t[[n[i]]]), 0))
    
    #Otherwise, just tally whether or not the descendant branches are both present states (=1), or split (=0)
    clust_b <- sapply(1:length(n), function(i) ifelse(db[i] == 2, 1, 0))
    names(clust_b) <- n
    
    #Note for node based clustering, include split state tips?
    clust_n <- sapply(n, function(i) ifelse(ns[as.character(i)] != 0, sum(t[desc_t[[i]]])/length(desc_t[[i]]), 0))
    
  }
  else{
    j <- rep(0, length(n))
  }
  
  #Calculate the two kinds of global clustering (by branch state gain lineages, vs. node state gain lineages...) 
  #coefficients for comparison, and normalize by the maximum clustering possible, 
  #assuming that each internal node of a clade leads to two presence state lineages: (Prevalence - 1) / (# Nodes):
  
  #New version, sum the proportion of presence state tip descendants per internal node.
  clust_coeff_n <- sum(clust_n) / Nnode(tree)
  
  if(sum(t) != 1) clust_coeff_b <- mean(clust_b) / ((sum(t)-1)/(Nnode(tree)))
  
  
  #Old version:
  #Note, that only internal nodes with only present tip descendants (== 1) are included.
  #This will ignore instances of nodes where descendant lossess have occured!

  #clust_coeff_n <- (length(which(clust_n == 1))/length(clust_n)) / ((sum(t)-1)/(Nnode(tree)))
  #clust_coeff_b <- (length(which(clust_b == 1))/length(clust_b)) / ((sum(t)-1)/(Nnode(tree)))
  
  ##There is also another thing to consider, which is partitioning, or the number of distinct clades the trait appears in.
  #For one needs to identify the earliest ancestral nodes (descendants from the root) where the trait is first gained, and the proportion of presence state tips descended from them.
  
  return(list(clust_coeff_n=clust_coeff_n,
              clust_coeff_b=clust_coeff_b,
              clust_n=clust_n,
              clust_b=clust_b))
}


#longevity_calc() - Calculate Longevity and Lability of Traits Based on Trait Lineages Identified by Ancestral State Reconstruction:
longevity_calc <- function(tree, ns){
  ts <- ns$tip_state
  bs <- ns$branch_state
  ns <- ns$node_state
  
  np <- nodepath(tree)
  
  
  #Also add the root node if it has a presence branch joining it:
  r <- which(tree$edge[,1] == Ntip(tree) + 1)
  r_br <- which(bs[r] == 1)
  
  if(length(r_br) != 0 & !as.character(Ntip(tree) + 1) %in% names(ns)){
    ns <- append(1, ns)
    names(ns)[1] <- Ntip(tree) + 1
  }
  
  n <- as.numeric(names(ns))
  
  #Get potential gain and loss nodes:
  gain_n <- which(ns == 1)
  loss_n <- which(ns == 0)
  
  #return(list(gain_n, loss_n))
  
  #Initialize output variables:
  longevity <- 0
  lability <- 0
  gain_times <- NULL
  loss_times <- NULL
  
  #Get the ancestral gain nodes for each presence tip:
  if(length(gain_n) != 0){
    ##Longevity:
    #For each presence tip, find the ancestral nodepath leading to their ancestral gain tip, 
    #i.e. identify where the branch state changes from presence to absence, or if it leads directly to the root.
    
    #Then calculate the sum of branchlength distances  and normalize by the maximu of all tip to root node branch
    #lengths, to get the trait longevity distribution.
    #Like RecPD, except multiple branches can be counted, and they are taken for each individual locus, 
    #to later come up with a mean, or distribution...
    
    dist <- dist.nodes(tree)[1:Ntip(tree), -(1:Ntip(tree))]
    #dist <- dist.nodes(tree)
    
    #longevity:
    
    #For presence tips, get sum of consecutive presence state branches.
    #Define a function to do this (should only be done once, outside of this function, as a helper):
    path_state <- function(x){
      x <- rev(x)
      i <- sapply(1:(length(x)-1), function(i){
        which(tree$edge[,2] %in% x[i] & tree$edge[,1] %in% x[i+1])
      })
      
      b <- bs[i]
      j <- which(b[1:(length(b)-1)] != b[2:length(b)])[1]
      
      if(is.na(j)) j <- length(b)
      
      return(sum(tree$edge.length[i[1:j]])/max(dist[,1]))
      #return(list(j, bs, r_tr$edge.length[i], sum(r_tr$edge.length[i[1:j]])/max(dn) ))
    }
    
    longevity <- array(unlist(lapply(np[which(ts == 1)], path_state)), dimnames=list(which(ts == 1)))
    
    #The old longevity calculation with node states: will not work in the cases where loss nodes occur
    #on gain lineages.
    # longevity <- sapply(np[which(ts == 1)], function(x){
    #   i <- ns[as.character(rev(x))]
    #   i <- which(i == 1)[1]
    #   p <- x[1:i]
    #   #t <- as.character(rev(x)[1])
    #   #n <- as.character(rev(x)[i])
    #   e <- which(tree$edge[,1] %in% p & tree$edge[,2] %in% p)
    #   bl <- sum(tree$edge.length[e])
    # 
    #   #return(list(tip=x, time=bl/max(dist[,1]), nodes=c(x[i], x[1]), edge=p))
    #   return(bl/max(dist[,1]))
    #   })
    
    
    
    
    #Gain times:
    #When do gain nodes appear, relative to the root... normalized by the maximum root-to-tip distance.
    
    gain_times <- sapply(n[gain_n], function(x){
      g <- nodepath(tree, x, Ntip(tree)+1)
      #t <- as.character(rev(x)[1])
      #n <- as.character(rev(x)[i])
      e <- which(tree$edge[,1] %in% g & tree$edge[,2] %in% g)
      bl <- sum(tree$edge.length[e])
      return(bl/max(dist[,1]))
    })
    
    #return(longevity)
    
    #Lability:
    #Calculate lability as the normalized number of state changes occurring in the tree (excluding the root):
    #the sum of internal node gain and loss events, divided by the number of internal tree nodes where events could potentially occur;
    
    #If the root appears as a node-state change, remove it?
    lability1 <- ifelse(as.character(Ntip(tree)+1) %in% names(ns), length(ns) - 1, length(ns)) / (Nnode(tree) - 1)
    
    
    #Or divided by the total number of internal nodes of each trait lineage where events could occur:
    ##To do this, need to identify the gain lineages (from gain_n) and loss lineages (from loss_n), and get the total number of unique
    ##descendant internal nodes from each...
    
    #Get gain lineage subtree nodes:
    s_n <- c(n[gain_n], n[loss_n])
    
    #If no loss lineages are identified, remove them:
    s_n <- s_n[which(!is.na(s_n))]
    
    #Get descendant nodes of each node state change lineage. 
    #Note some gain/loss nodes may be descended from other ancestral gain/loss nodes, remove redundant nodes.
    st_n <- unlist(Descendants(tree, s_n, type='all'))
    st_n <- unique(st_n[which(st_n >= Ntip(tree) + 2)]) # +2 - remove root?

    #Distinguish the ancestral gain and loss lineage nodes from the descendant nodes of each gain or loss lineages,
    #those will be the actual node state change nodes:

    lability2 <- length(which(s_n %in% st_n))/length(st_n)
    
    
    #Or, just take the number of internal gain and loss event nodes and
    #divide by the number of gain and loss tips (excluding absences), descended from them.
    #Note, lability == 1 when all gain/loss states result directly from a gain/loss event, i.e. later occuring events,
    #while lower levels of lability indicates gains/losses occuring earlier in the tree.
    #and lability == 0 when all gain nodes are directly descended from an 
    #ancestral gain event (have to subtract each category by 1 if only a single gain node exists).
    
    if(length(gain_n) == 1){
      lability3 <- ((length(which(ns == 1)) - 1) + length(which(ns == 0)))/ length(which(ts != -1))
    }
    else{
      lability3 <- ((length(which(ns == 1))) + length(which(ns == 0)))/ length(which(ts != -1))
    }
    

    if(length(loss_n) != 0){
      
      #If any internal loss nodes are found, also add them to the longevity array.
      #Get the sum of branch lengths between ancestral gain -> loss node lineages.
      #and normalize by max tip to root distance.
      
      #Get the branch state distances from loss nodes to ancestral gains:
      #i <- as.numeric(names(ns[[2]]$node_state)[which(ns[[2]]$node_state == 0)])
      
      l <- lapply(n[loss_n], function(x) nodepath(phy=tree, from=Ntip(tree)+1, to=x))
      
      longevity <- append(longevity, array(unlist(lapply(l, path_state)), dimnames=list(n[loss_n])))
      
      
      # long_internal <- sapply(n[loss_n], function(x){
      #   i <- nodepath(tree, x, Ntip(tree)+1)
      #   g <- which(ns[as.character(i)] != 0)[1]
      #   #l <- which(ns[as.character(i)] == 0)[1]
      #   e <- which(tree$edge[,1] %in% i[1:g] & tree$edge[,2] %in% i[1:g])
      #   bl <- sum(tree$edge.length[e])
      #   return(bl/max(dist[,1]))
      #   #return(list(node=x, time=bl/max(dist[,1]), nodes=c(i[g], x), edge=i[1:g]))
      # })
      # 
      # longevity <- append(longevity, long_internal)
      
      #Loss times:
      #Similar to gain times, but is the distance from the internal loss node relative to the root:
      loss_times <- sapply(n[loss_n], function(x){
        l <- nodepath(tree, x, Ntip(tree)+1)
        #t <- as.character(rev(x)[1])
        #n <- as.character(rev(x)[i])
        e <- which(tree$edge[,1] %in% l & tree$edge[,2] %in% l)
        bl <- sum(tree$edge.length[e])
        return(bl/max(dist[,1]))
      })
    }
  }
  
  return(list(longevity=longevity, 
              lability1=lability1,
              lability2=lability2,
              lability3=lability3,
              gain_times=gain_times, 
              loss_times=loss_times))
  
}


####
#recpd_cor_calc() - Extension of Jaccard similarity of two trait branch lineage distributions, 
#based on the sum of branch lengths shared between two traits, normalized by the total sum of unique trait lineage branches.
#this version also outputs the unweighted branch state jaccard overlap, and jaccard overlap using only phylogenetic tree tips.
####

#Wrap this function to generate a pairwise correlation matrix, or binary list?

recpd_cor_calc <- function(tree, state_x, state_y){
  #state_x - recpd state annotations for trait distribution 1,
  #state_y - recpd state annotations for trait distribution 2.
  
  
  #br_x - recpd annotated branch states for trait distribution 1.
  #br_y - recpd annotated branch states for trait distribution 2.
  br_x <- state_x$branch_state
  br_y <- state_y$branch_state
  
  #ts_x - recpd annotated tip states for trait distribution 1.
  #ts_y - recpd annotated tip states for trait distribution 2.
  ts_x <- state_x$tip_state
  ts_y <- state_y$tip_state
  
  #tree - the species tree.
  
  #Identify which gain branches are present in both x and y trait lineages (states = 1),
  #and, which loss branches are absent in both x and y trait lineages (states = 0)?
  ##Note that absent branches (state = -1) are excluded.
  ol_b <- which(br_x == 1 & br_y == 1)# | br_x == 0 & br_y == 0)
  
 
  #Identify all unique branches which are present in x and y trait lineages (states = 1/1, 1/0, 0/1):
  tot_b <- which(br_x == 1 | br_y == 1)
  
  
  #Calculate RecPDcor, by taking the sum of shared gain and loss branch lengths between both trait lineages, 
  #divided by the sum of all tree branch lengths represented by either trait distribution lineage.
  #This is equivalent to a jaccard similarity of trait lineage branches, weighted by tree branch lengths.
  
  #Also, for comparison purposes, calculate the unweighted recpd trait lineage jaccard similarity.
  
  #Also calculate the tip gain/loss node jaccard overlap.
  ol_t <- which(ts_x == 1 & ts_y == 1)
  tot_t <- which(ts_x == 1 | ts_y == 1)
  
  rpd_cor <- list(recpd_cor = sum(tree$edge.length[ol_b])/sum(tree$edge.length[tot_b]),
                  recpd_jacc = length(ol_b)/length(tot_b),
                  tip_jacc = length(ol_t)/length(tot_t)
  )
  
  return(rpd_cor)
}


####
#Functions To Plot Out The Species Phylogenetic Tree With Trait Distributions and Their Node and Branch Ancestral State Annotations 
#Inferred using recpd_calc().
####

#plt_tree_f() - requires the 'ggtree' package.
plt_tree_f <- function(tree, br_l){
  tr <- as_tibble(tree)
  
  labs <- array(c('Absence', 'Loss', 'Gain', 'Split'), dimnames=list(c('-1', '0', '1', '2')))
  col <- array(c('#C0C0C0','#F8766D', '#00B81F', '#00C0B8'), dimnames=list(c('-1', '0', '1', '2')))
  
  lab <- as.character(br_l$branch_state)
  
  branchcol <- tibble(data.frame(parent=tree$edge[,1], node=tree$edge[,2]), branch_col=as.numeric(br_l$branch_state))
  
  nodecol <- tibble(node=c(1:Ntip(tree), as.numeric(names(br_l$node_state))), node_col=as.numeric(c(br_l$tip_state, br_l$node_state)))
  
  tr <- as_tibble(tree)
  
  tr <- full_join(tr, branchcol, by=c('parent' = 'parent', 'node' = 'node'))
  tr <- full_join(tr, nodecol, by=c('node'='node'))
  
  #Assign branch lengths to stroke width aesthetic:
  #Group branch lengths into bins based on quantiles.
  branch_bins <- cut(tr$branch.length, quantile(tr$branch.length, na.rm=TRUE), include.lowest=TRUE)
  b_size <- lapply(strsplit(as.character(levels(branch_bins)),  ','), function(x) as.numeric(sub('[\\(\\)\\[\\]]', '', x, perl=TRUE)))
  
  #Format the branch length bins to be more legible.
  bin_level <- str_match_all(levels(branch_bins), '([\\(\\[])(.*),(.*)(])')
  names(bin_level) <- levels(branch_bins)
  relab <- unlist(lapply(bin_level, function(x) paste0(x[2], formatC(as.numeric(x[3]), format='e', digits=2), ', ', formatC(as.numeric(x[4]), format='e', digits=2), x[5])))
  
  branch_bins <- factor(as.vector(relab[branch_bins]), levels=relab)
  
  #Set the branch lengths to a unit size.
  tr2 <- tr
  tr2$branch.length <- 0.001
  
  #Plot the tree with state mappings:
  gt <- ggtree(as.treedata(tr2), aes(color=factor(as.character(branch_col), levels=c('0', '1', '-1', '2')), size=as.numeric(branch_bins)), layout='slanted') + 
    #Tip gain/loss/absence:
    geom_tippoint(aes(subset=(!is.na(node_col) & grepl('\\w', label)), fill=factor(as.character(node_col), levels=c('0', '1', '-1', '2'))), size=1) +
    #Internal node gain/loss/absence:
    geom_nodepoint(aes(subset=!is.na(node_col), fill=factor(as.character(node_col), levels=c('0', '1', '-1', '2'))), shape=24, color='black', size=2) + 
    layout_dendrogram() +
    #Specify the aesthetics values:
    scale_size_continuous(name='Branch Length', 
                          labels=levels(branch_bins), 
                          range=c(0.25, 1.0), 
                          guide=guide_legend(order=1)) + 
    scale_color_manual(name='Node and Edge', 
                       labels=labs[sort(unique(as.character(tr$node_col)))], 
                       values=col[sort(unique(as.character(tr$node_col)))], 
                       na.translate=FALSE,
                       guide=guide_legend(order=2)) +
    scale_fill_manual(name='Node and Edge', 
                      labels=labs[sort(unique(as.character(tr$node_col)))], 
                      values=col[sort(unique(as.character(tr$node_col)))], 
                      na.translate=FALSE,
                      guide=guide_legend(order=2)) +
    theme(legend.position='right')
  
  #Have to do the same for absent state branches...
  
  #For older node mappings with split nodes:
  #Old split color: '#00C0B8', also have to add na.value='#00C0B8' to ensure consistent color mapping for trait distributions
  #where no splits occur, and set na.translate=TRUE.
  
  #Add The Y-Axis Which Gives Branch Length From Root To Tip?
  if(0){
    gt <- gt + 
      geom_treescale() +
      coord_flip() +
      scale_x_reverse() +
      theme(axis.line.y=element_line(color='black'),
            axis.text.y=element_text(),
            axis.ticks.y =element_line())
  }
  
  gt
}


#plt_tree_f_old() - The Equivalent Function As plt_tree_f(), but using plot.phylo from the ape package (base R graphics):
plt_tree_f_old <- function(tree, br_l, test='-a', branch.lengths=TRUE, axis=FALSE){
  #Test = Which mappings to add:
  #-a - none
  #t - tips
  #n - internal nodes
  #b - branches
  #a - all
  
  #branch.lengths = TRUE/FALSE - use tree branch lengths (TRUE), otherwise set all branch lengths to the same length.
  #axis = TRUE/FALSE - add a y-axis representing tree branch lengths to the root.
  
  #Node colorings by presence/absence of descendants
  
  n_col <- array(c('grey', 'red', 'green', 'blue'), dimnames=list(as.character(c(-1, 0, 1, 2))))
  
  br_col <- array(c('grey', 'red', 'green'), dimnames=list(as.character(c(-1, 0 ,1))))
  
  if('b' %in% test || 'a' %in% test){
    brcol <- br_col[as.character(br_l$branch_state)]
  }
  else{
    brcol <- rep('black', length(tree$edge.length))
    
  }
  
  #If branch.lengths == FALSE, 
  #Then set all of the tree edge lengths to the same size, 
  #and do not show the branch length axis.
  
  if(branch.lengths == FALSE) tree$edge.length <- rep(1e-6, length(tree$edge.length))
  
  plt <- plot(tree,
              use.edge.length=TRUE,
              direction='downwards',
              show.tip.label=FALSE,
              edge.color=brcol,
              plot=TRUE)
  
  # plot(tree, 
  #      use.edge.length=plt$use.edge.length,
  #      direction=plt$direction,
  #      show.tip.label=plt$show.tip.label, 
  #      edge.color=brcol)
  
  #Plot y-axis: branch length to root? (axis == TRUE):
  if(axis){
    axis(2, at=c(seq(plt$y.lim[2], 0, -1), 0), labels=c(0:floor(plt$y.lim[2]), signif(plt$y.lim[2], 3)), las=2, line=1, cex.axis=0.8)
    
    mtext('Branch Length to Root', 2, line=3.5, cex=0.8)
  }
  
  if('t' %in% test || 'a' %in% test){
    if(Ntip(tree) < 100) cex <- 1 else cex <- 0.5
    
    tiplabels('', 1:Ntip(tree), 
              frame='none',
              pch=16,
              cex=cex,
              #pch=ifelse(br_l[[prev_c]][[i]]$tip_col == 1, 16, 1),
              col=n_col[as.character(br_l$tip_state)],
              adj=c(0.5,0.5)
    )
  }
  
  if('n' %in% test || 'a' %in% test){
    nodelabels('', 
               as.numeric(names(br_l$node_state)),
               frame='none',
               pch=24,
               cex=1.25,
               bg=n_col[as.character(br_l$node_state)],
               col='black')
  }  
}


#plt_cor_tr() - Visualizes The Overlap of Ancestral States For A Selected Pair of Trait Lineages.
#* requires the 'ggtree' package.
plt_cor_tree <- function(tree, br_x, br_y, lab=NULL, colnames.offset=0, sep.col='transparent'){
  #br_x - annotation state list (tip, node, branch) for one trait distribution
  #br_y - annotation state list (tip, node, branch) for a second trait distribution
  
  #lab - Labels for the presence/absence strips (traits)
  
  
  #Bind together tip locus presence absence state
  pa <- cbind(as.character(ifelse(br_x$tip_state == 1, 1, 0)),
              as.character(ifelse(br_y$tip_state == 1, 1, 0)))
  rownames(pa) <- tree$tip.label
  
  #Add column labels, if present:
  if(length(lab) != 0) colnames(pa) <- lab
  
  #rownames(pa) <- rowSums(pa)
  #pa <- apply(pa, 1, function(x) ifelse(x != 0, 'Present', 'Absent'))
  
  #bind together the edge mappings:          
  b <- rbind(ifelse(br_x$branch_state == 1, 1, 0), 
             ifelse(br_y$branch_state == 1, 1, 0))
  
  b_t <- data.frame(cbind(node=tree$edge[,2], branch_col=apply(b, 2, sum)))
  
  tr <- full_join(as_tibble(tree), b_t)
  
  #Assign phylogenetic tree branch lengths into bins (by quantiles):
  #br_bins <- cut(tr$branch.length, seq(floor(min(signif(tr$branch.length, 1), na.rm=TRUE)), ceiling(max(signif(tr$branch.length, 1), na.rm=TRUE)), 0.1), include.lowest=TRUE)
  
  br_bins <- cut(tr$branch.length, signif(quantile(tr$branch.length, na.rm=TRUE), 2), include.lowest=TRUE)
  
  #Format the branch length bins to be more legible.
  bin_level <- str_match_all(levels(br_bins), '([\\(\\[])(.*),(.*)(])')
  names(bin_level) <- levels(br_bins)
  relab <- unlist(lapply(bin_level, function(x) paste0(x[2], formatC(as.numeric(x[3]), format='e', digits=2), ', ', formatC(as.numeric(x[4]), format='e', digits=2), x[5])))
  
  br_bins <- factor(as.vector(relab[br_bins]), levels=relab)
  
  
  #Extract the lower bound of the branch length bins for size mapping.
  br_size <- sapply(sapply(br_bins, function(x) as.numeric(unlist(strsplit(gsub('[\\(\\)\\[\\]]', '', as.character(x), perl=TRUE), split=',')))), function(x) x[1])
  
  #Branch colouring if they are shared between the two locus distributions
  branch_cols <- brewer.pal(3, 'Set1')
  names(branch_cols) <- c('0', '1', '2')
  branch_labs <-  c('None', 'Unique', 'Shared')
  names(branch_labs) <- c('0', '1', '2')
  
  tr$branch_col <- factor(tr$branch_col, levels=c(0,1,2), exclude=NA)
  
  #Normalize branch lengths:
  tr <- tr %>% mutate(branch.length.std=1e-6)
  
  
  #Make the tree:
  gt <- ggtree(as.treedata(tr), 
               aes(color=branch_col, size=br_bins), #Note: exponentially transform branch lengths...?
               branch.length='branch.length.std',
               #lineend='square',
               layout='slanted') +
    scale_color_manual(values=branch_cols, 
                       labels=branch_labs,
                       na.translate=FALSE) +
    scale_size_manual(breaks=levels(br_bins),
                      values=(1/2)*1:length(levels(br_bins)),
               labels=levels(br_bins)) +
    labs(size='Branch Length',
         color='Trait Branch Overlap') +
    theme(line=element_line(lineend='square'))
  
  
  #Append the presence/absence strips for the pair of traits:
  
  ##Plot column labels if they are supplied:
  if(is.null(lab)) cn <- FALSE else cn <- TRUE
  
  gheatmap(gt, data.frame(pa), 
           width=0.2, 
           color=sep.col, 
           colnames_position='top', colnames=cn, colnames_offset_y=colnames.offset, colnames_angle=45, hjust=0) + 
    scale_fill_manual(name='Tip State',
                      labels=c('Absent', 'Present'), 
                      values=c('white', 'black')) + 
    ylim(0, NA) +
    labs(
      size='Branch Length',
      fill='Trait Presence')
  
  #Note: column names might not fit in the plot for large trees...
  #Can be adjusted by changing colnames_offset_y and expanding plot limits.
  
}


###
#plt_tree_faith() - tree plotting function for faith's PD examples.
plt_tree_faith <- function(tree, node_state, include.root=TRUE){
  #Get the tips presence/absence states:
  t <- node_state$tip_state
  
  #Corresponding normalized Faith's PD values:
  f <- pd(matrix(t, 1, Ntip(tree), dimnames=list(NULL, tree$tip.label[as.numeric(names(t))])), tree, include.root)
  
  f <- f/sum(tree$edge.length)
  
  #Faith branch length mappings - if include.root=TRUE for the PD calculation:
  n <- unique(unlist(nodepath(tree)[which(t == 1)]))
  
  #If include.root=FALSE, then remove nodes between the root and the MRCA of the present tips.
  if(include.root == FALSE){
    m <- getMRCA(tree, which(t == 1))
    #If the MRCA is not the root
    if(m != tree$edge[1,1]){
      nr <- nodepath(tree, n[1], m)
      nr <- nr[-length(nr)]
      n <- n[-which(n %in% nr)]
    }
  }
  
  #Make a temporary node and edge mapping list, with branch lengths coded as those which are included for Faith's PD calculation.
  ns_tmp <- list()
  ns_tmp$tip_state <- node_state$tip_state
  ns_tmp$branch_state <- ifelse(tree$edge[,1] %in% n & tree$edge[,2] %in% n, 1, 0)
  ns_tmp$node_state <- array(rep(NA, length(node_state$node_state)), dimnames=list(names(node_state$node_state)))
  
  #return(ns_tmp)
  
  gt <- plt_tree_f(tree, ns_tmp) + labs(title=paste0('Faith\'s PD = ', signif(f[1,1], 3)))
  
  #labs(title=paste0('Faith\'s PD\n', 'With Root = ', signif(f[1,1], 3), '\n', 'Without Root = ', signif(f[2,1], 3)))
  
  gt
}


  
```
