## Supplementary material for "RecPD: A Recombination-Aware Measure of Phylogenetic Diversity": RecPD Tutorial: S1 File - recpd_tutorial.html


### RecPD Tutorial

###### Dr. Cedoljub Bundalovic-Torma

#### 25/09/2021

- 1 Packages Required
- 2 Overview of RecPD
  - 2.1 Randomized Trait Distribution Generation - Version 1
  - 2.2 Randomized Trait Distribution Generation - Version 2

### 1 Packages Required

```
library(ape)
library(phangorn)
library(picante)
library(phytools)

library(dplyr)
library(tidyr)
library(tidytree)
library(ggplot2)
library(ggtree)
library(ggtreeExtra)
library(ggnewscale)
library(ggpubr)
library(RColorBrewer)
library(tibble)
library(tidyr)
library(stringr)


#Set the working directory (linux), relative to where this tutorial is located:
setwd('/media/cedatorma/Seagate Expansion Drive/psy_project/t3se_clusters/recpd/recpd_tutorial/')
```

Load the RecPD functions:

```
#Load RecPD Functions
source('./recpd_functions.R')
```

### 2 Overview of RecPD

RecPD requires two components (user provided), a species tree, and a presence/absence matrix of traits mapped to the tips of the tree.

To illustrate the different functionalities of RecPD and how to process the metrics it generates, we’ll be using randomly generated phylogenetic trees and presence/absence tip state matrices.

#### 2.1 Randomized Trait Distribution Generation - Version 1

The first approach is to begin with a randomly generated species tree of 10 tips.

```
r_tr <- rtree(10)
```

The following code chunk provides a function which will generate a matrix of all possible presence/absence states on a given tree. Note that this works for a tree of 10 tips, however, this will be computationally unfeasable for trees of larger size (the number of possible presence/absence states combinatorially explodes).

```
#A function which generates a matrix of permuted locus distributions with given prevalence across N tips:

locus_rand <- function(tree, prevalence){
  #Prevalence (P) indicates the total number of tips with a state present
  
  #Steps: 
  #1) Set the tip states as an array of length equal to the number of tips in the tree, initialized to 0.
  #2) Generate a set of starting arrays with a single presence states assigned to successive indicies, 
  ##stopping after reaching an index position P steps away from the end of the array.
  #3) From these starting arrays, repeat the procedure above and iteratatively add an additional presence state until reaching maximum prevalence.
  
  #This function should be defined outside of locus_rand, so it does not have to be initialized every time locus_rand is called.
  array_return <- function(a){
    a_tmp <- NULL
    a_return <- NULL
    i <- which(a == 1)
    
    for(j in i[length(i)]:length(a)){
      a_tmp <- a
      a_tmp[j] <- 1
      a_return <- rbind(a_return, a_tmp, deparse.level=0)
    }
    return(a_return)
  }
  
  l_tmp <- NULL
  a_tmp <- rep(0, Ntip(tree))
  names(a_tmp) <- tree$tip.label
  
  for(i in 1:(Ntip(tree) - prevalence + 1)){
    a <- a_tmp
    a[i] <- 1
    l_tmp <- rbind(l_tmp, a, deparse.level=0)
  }
  
  n <- 2
  while(n <= prevalence){
    for(i in 1:nrow(l_tmp)){
      l_tmp <- rbind(l_tmp, array_return(l_tmp[i,]), deparse.level=0)
    }
    
    l_tmp <- l_tmp[-which(rowSums(l_tmp) < n),]
    n <- n + 1
  }
  
  if(is.null(nrow(l_tmp))){
    l_tmp <- as.matrix(t(l_tmp))
  }
  
  return(l_tmp)
}
```

Next, assign randomized presence/absence trait states to the tips of the species tree.

```
#Generate a list of randomized tip state/locus presence/absence patterns for a given tree:
#Do not include prevalence == Ntip(tree)?

pa_l <- data.frame()

for(i in 1:(Ntip(r_tr)-1)){
  pa_l <- rbind.data.frame(pa_l, data.frame(locus_rand(r_tr, i)))
}
```

To illustrate the output of RecPD, we’ll calculate RecPD for a single each randomized trait distribution.

```
res <- recpd_calc(r_tr,   #The species tree
             pa_l[1,],    #The trait presence/absence state matrix, rows = trait stat, columns = corresponding tips of the species tree.
             option='nn', #Ancestral state reconstruction approach to use ('nn', 'mpr', or 'ace')
             calc=TRUE    #Calculate RecPD derived metrics? (default = FALSE)
             )
```

For each trait, RecPD will produce a list of results: - “metrics” : the RecPD metric and RecPD-derived metrics (if calc == TRUE) - “node\_state” : the final trait evolutionary history reconstructions, with states assigned to tips, internal nodes, and branches of the species phylogenetic tree. These will be used for phylogenetic tree visualizations. - “ns\_old” : preliminary ancestral state reconstructions.

```
names(res[[1]])
```

```
## [1] "metrics"    "node_state" "ns_old"
```

Let’s look inside “metrics”:

```
unlist(res[[1]]$metrics)
```

```
##      recpd       span    cluster  longevity   lability 
## 0.06365753 0.00000000 0.00000000 0.10713412 0.12500000
```

The following code consolidates the RecPD metric results for all of the trait distributions.

```
#For each tree, calculate RecPD (nn, mpr ace) and faith's PD of each set of randomized trait distributions.
test_res1 <- data.frame()

{
  #Calculate RecPD for randomized trait distributions using NN, MPR, and ACE methods:
  r_nn <- recpd_calc(r_tr, pa_l, option='nn')
  r_mpr <- recpd_calc(r_tr, pa_l, option='mpr')
  r_ace <- recpd_calc(r_tr, pa_l, option='ace')
  
  #Also calculate Faith's PD:
  faith <- pd(pa_l, r_tr, include.root=FALSE)
  
  
  #Store the results of the RecPD analysis
  ntip <- Ntip(r_tr)

  test_res1 <- rbind.data.frame(test_res1, 
                               data.frame(ntip=ntip,
                                          prevalence=faith[,2],
                                          faith=faith[,1]/sum(r_tr$edge.length),
                                          recpd_nn=unlist(lapply(r_nn, function(x) x$metrics$recpd)),
                                          recpd_mpr=unlist(lapply(r_mpr, function(x) x$metrics$recpd)),
                                          recpd_ace=unlist(lapply(r_ace, function(x) x$metrics$recpd)))
                               )
  
  
}
```

Now, visualize the RecPD vs. Faith’s PD metrics calculated for each randomized trait distribution:

```
#Plot the results:

#RecPD (NN, MPR, ACE) and Faith's PD:
plt1 <- ggplot(test_res1 %>% pivot_longer(3:6),
       aes(factor(prevalence), value, color=name)) +
  geom_abline(intercept=0, slope=0.1, lty=2, lwd=0.5) +
  geom_boxplot(outlier.size=1) +
  labs(title='RecPD (NN, MPR, ACE) vs. Faith\'s PD',
       #subtitle='Tree Size - 10 Tips, 1022 Randomized Gene-Family Distributions',
       x='Prevalence',
       y='Phylogenetic Diversity',
       color='Metric') + 
  theme(plot.title=element_text(size=12),
        axis.text.x=element_text(size=10),
        axis.text.y=element_text(size=10),
        axis.title.x=element_text(size=12),
        axis.title.y=element_text(size=12),
        legend.title=element_text(size=12),
        legend.text=element_text(size=12),
        plot.margin=unit(c(5,10,20,10), 'points'))


#All metrics - normalized to Faith's PD:
plt2 <- ggplot(test_res1 %>% pivot_longer(4:6), 
       aes(factor(prevalence), value/faith, color=name)) + 
  geom_hline(yintercept=1, lty=2, lwd=0.5) +
  geom_boxplot(outlier.size=0.5) +
  labs(title='RecPD - NN, MPR, and ACE Normalized by Faith\'s PD',
       #subtitle='Tree Size - 10 Tips, 1022 Randomized Gene-Family Distributions',
       x='Prevalence',
       y='Normalized RecPD\n(RecPD / Faith\'s PD)',
       color='Metric') + 
  theme(plot.title=element_text(size=12),
        axis.text.x=element_text(size=10),
        axis.text.y=element_text(size=10),
        axis.title.x=element_text(size=12),
        axis.title.y=element_text(size=12),
        legend.title=element_text(size=12),
        legend.text=element_text(size=12),
        plot.margin=unit(c(5,10,20,10), 'points'))

plts1 <- ggarrange(plt1, plt2, ncol=1, legend='right')

plts1
```

Now lets use the results of the RecPD to visualize the evolutionary histories of a given trait mapped onto the species tree.

```
#For example, trait distribution 427
i <- 427

#plt_tr_faith() can be used to visualize evolutionary histories based on vertical ancestry, i.e. that which is inferred using Faith's PD:
plt3 <- plt_tree_faith(r_tr, r_nn[[i]]$node_state)  + 
  theme(plot.title=element_text(size=12),
        plot.margin=unit(c(25,10,25,10), 'points'),
        legend.title=element_text(size=12),
        legend.text=element_text(size=12))


#plt_tree_f() is used for visualizing RecPD evolutionary histories, providing the species tree and the node_state list produced by recpd_calc() for a specific trait distribution in the presence/absence matrix.

#RecPD using Nearest-Neighbours:
plt4 <- plt_tree_f(r_tr, r_nn[[i]]$node_state) +
  labs(title=paste0('RecPD_nn = ', signif(test_res1$recpd_nn[i], 3))) + 
  theme(plot.title=element_text(size=12),
        plot.margin=unit(c(25,10,25,10), 'points'),
        legend.title=element_text(size=12),
        legend.text=element_text(size=12))

#RecPD using MPR:
plt5 <- plt_tree_f(r_tr, r_mpr[[i]]$node_state) +
  labs(title=paste0('RecPD_mpr = ', signif(test_res1$recpd_mpr[i], 3))) + 
  theme(plot.title=element_text(size=12),
        plot.margin=unit(c(25,10,25,10), 'points'),
        legend.title=element_text(size=12),
        legend.text=element_text(size=12))

#RecPD using ACE
plt6 <- plt_tree_f(r_tr, r_ace[[i]]$node_state) +
  labs(title=paste0('RecPD_ace = ', signif(test_res1$recpd_ace[i], 3))) + 
  theme(plot.title=element_text(size=12),
        plot.margin=unit(c(25,10,25,10), 'points'),
        legend.title=element_text(size=12),
        legend.text=element_text(size=12))


plts2 <- ggarrange(plt3, plt4, 
                   #ggplot() + theme_void(), 
                   plt5, plt6, 
                   #ggplot() + theme_void(), 
                   common.legend=TRUE, legend='right')


plts2
```

Using RecPD to calculate correlations between the evolutionary histories for pair of trait distributions.

```
#Calculation, will output a list of metrics for comparison:
##recpd branch length correlations (recpd_cor), 
##unweighted recpd branch length correlations (recpd_jacc),
##tip state Jaccard similarity (tip_jacc).

recpd_cor_calc(r_tr, #Species tree.
               r_nn[[100]]$node_state, #RecPD node state mappings for trait distribution 1.
               r_nn[[101]]$node_state  #RecPD node state mappings for trait distribution 2.
)
```

```
## $recpd_cor
## [1] 0.622565
## 
## $recpd_jacc
## [1] 0.625
## 
## $tip_jacc
## [1] 0.5
```

```
#Plotting the RecPD evolutionary histories for a pair of traits on a species phylogeny:
plt_cor_tree(r_tr,
           r_nn[[100]]$node_state,
           r_nn[[101]]$node_state,
          sep.col='grey50')
```

#### 2.2 Randomized Trait Distribution Generation - Version 2

For larger trees, the alternative is to randomly sample a number of presence/absence tip assignments, at different levels of prevalence.

```
#A new version of locus_rand2 which allows trees of different sizes to be examined through sampling a certain number of locus distributions:
#using sample() to select which tips will be present:

locus_rand2 <- function(r_tr, prevalence=1, nsamp=10){
  tip <- r_tr$tip.label
  
  dist <- data.frame()
  
  i <- 1
  check <- NULL
  while(i <= nsamp){
    d <- ifelse(tip %in% sample(tip, prevalence), 1, 0)
    
    #check if the trait distribution has previously been generated:
    if(i != 1) check <- which(apply(dist, 1, function(x) identical(as.numeric(x), d)) == TRUE)
    
    if(length(check) == 0){
      dist <- rbind.data.frame(dist, d)
    
      i <- i + 1
      check <- NULL
    }
    
    #Will produce inf if the size of tree tips are <= 200
    if(length(tip) < 200 & i > factorial(length(tip))/(factorial(length(tip)-prevalence)*factorial(prevalence))) break
    
    if(length(tip) >= 200 & prevalence  == length(tip)) break
    
  }
  
  colnames(dist) <- tip
  return(dist)
}


#A function to generate corresponding randomized trait distributions for each tree:
dist_gen <- function(tree_l, nsamp=10){
  #tree_l - a list of randomly generated trees
  #nsamp - the number of randomly generated locus distributions to sample
  
  #A list to store the randomized distributions generated for each tree topology:
  dist_l <- list()
  
  for(i in 1:length(tree_l)){
    dist_l[[i]] <- list()
      
   #To make prevalence comparable between trees of different sizes,
    #Select the prevalence cutoffs using percentile ranges:
    prev <- trunc(quantile(1:Ntip(tree_l[[i]]), probs=seq(0.1, 1, 0.1)))
    
    
    for(j in prev){
      dist_l[[i]] <- rbind.data.frame(dist_l[[i]], 
                                locus_rand2(tree_l[[i]], prevalence=j, nsamp=nsamp))

    }
  }
  
  return(dist_l)
}
```

Generate sets of randomized trait distributions for each tree:

```
#Generate a list of trees of different size:
tree_l <- lapply(c(100, 500, 1000), rtree)

#Generate random trait distributions for each tree:
dist_l <- dist_gen(tree_l, nsamp=10)

#For each tree, calculate RecPD (nn, mpr ace) and faith's PD of each set of randomized trait distributions.
test_res2 <- data.frame()
j <- 1 

for(i in 1:length(dist_l)){
  #Note when just the tree toplogy doesn't change, but the branchlength distributions do, 
  #do not change the tip state distributions!
  r_nn <- recpd_calc(tree_l[[i]], dist_l[[j]], option='nn')
  r_mpr <- recpd_calc(tree_l[[i]], dist_l[[j]], option='mpr')
  r_ace <- recpd_calc(tree_l[[i]], dist_l[[j]], option='ace')

  faith <- pd(dist_l[[j]], tree_l[[i]], include.root=FALSE)

  ntip <- Ntip(tree_l[[i]])

  
  test_res2 <- rbind.data.frame(test_res2, 
                               data.frame(ntip=ntip,
                                          ntree=j,
                                          prevalence=faith[,2],
                                          faith=faith[,1]/sum(tree_l[[i]]$edge.length),
                                          #faith=faith[,1]/sum(tree_l[[i]][[j]]$edge.length),
                                          recpd_nn=unlist(lapply(r_nn, function(x) x$metrics$recpd)),
                                          recpd_mpr=unlist(lapply(r_mpr, function(x) x$metrics$recpd)),
                                          recpd_ace=unlist(lapply(r_ace, function(x) x$metrics$recpd)))
                               )
  j <- j + 1
    
}

test_res2 <- test_res2 %>% mutate(ntip=as.numeric(as.character(ntip)))


#RecPD normalized to Faith's using different tree sizes:
ggplot(test_res2 %>% pivot_longer(5:7), 
       aes(factor(prevalence/ntip), value/faith, color=name)) + 
  geom_hline(yintercept=1, lwd=0.3, lty=2) +
  geom_boxplot() + 
  facet_wrap(~ntip, nrow=1, labeller=labeller(ntip=function(x) paste('Tree Size =', x, 'tips')), scales='free_x') +
  scale_y_continuous(breaks=c(0.5, 1, 1.5)) +
  labs(title='RecPD - NN, MPR, ACE Normalized by Faith\'s PD',
       subtitle='Varying Tree Sizes',
       x='Trait Prevalence (Proportion of Tips)',
       y='Normalized RecPD\n(RecPD / Faith\'s PD)',
       color='Metric') +
  theme(legend.position='bottom')
```
