## Supplementary material for "RecPD: A Recombination-Aware Measure of Phylogenetic Diversity": Evolved Trait Histories: S2 File - evolved_trait_histories.html


### Evolved Trait Histories

###### Dr. Cedoljub Bundalovic-Torma

#### 25/09/2021

- 1 Packages Required
- 2 Gene-Family Evolution Simulation
  - 2.1 Functions
    - 2.1.1 lineage\_get()
    - 2.1.2 locus\_evolve()
    - 2.1.3 event\_label()
    - 2.1.4 plot\_evo\_tree()
- 3 Simulate trait evolutionary histories
  - 3.1 RecPD of simulated trait distributions

### 1 Packages Required

```
library(ape)
library(phangorn)
library(picante)
library(phytools)

library(dplyr)
library(tidyr)
library(tidytree)
library(ggplot2)
library(ggtree)
library(ggtreeExtra)
library(ggnewscale)
library(ggpubr)
library(RColorBrewer)
library(tibble)
library(tidyr)
library(stringr)


#Set the working directory (linux), relative to where this tutorial is located:
setwd('/media/cedatorma/Seagate Expansion Drive/psy_project/t3se_clusters/recpd/recpd_tutorial/')
```

Load the RecPD functions:

```
#Load RecPD Functions
source('./recpd_functions.R')
```

### 2 Gene-Family Evolution Simulation

This is the latest version of the locus/state evolution modelling code. This approach adapts some ideas from the generation of phylogenetic trees using the Birth-Death process, see: https://lukejharmon.github.io/pcm/chapter10\_birthdeath/

The essence of the method models a poisson process of events (locus/state losses and recombinations) occurring along the different lineages of a provided species phylogenetic tree. The tree can be either ultra-metric or non-ultrametric, with branch-lengths representing either time since emergence from a common ancestor, or a molecular evolutionary distance (average expected nucleotide/aminoacid substitutions per site).

The model requires two parameters, specifying the rate exponents for each type of event:

- Extinction/Death/Loss Rate (Er): losses of a locus/state
- Recombination Rate (Rr: gains of a locus/state from one species lineage to another

These rates can be thought of as a summary of the evolutionary selective pressures acting to maintain a locus/state or its selective advantage helping to propagate it. Note that these rates remain constant throughout the evolutionary history of the species tree, however in reality they are likely to vary under different population bottlenecks or changing environments.

Using these rates the interarrival time distribution for any event (loss or transfer) can be calculated using the exponential distribution: P(ti) = (Er + Rr) \* exp(-(Er + Rr)\*ti).

In addition, probabilities of events occurring is given by:

- Extinction Probability : Er / (Er + Rr)
- Recombination Probability : Rr / (Er + Rr)

First, the emergence time of the locus/trait (birth\_time) is randomly drawn from a uniform distribution and then randomly assigned to a species lineage existing at that time (locus/state origination event). Next, the occurrence of events is modelled using a poisson process by randomly drawing a sequence of interarrival times from the inverse cumulative probability function of the exponential distribution: e\_t = -log(1-P(i))/(Er + Rr + Lr)), where i is a random variable sampled from a uniform distribution taking values from [0-1].

The inter-arrival time sequence is cumulatively summed and then added to the emergence time of the first event (cumsum(e\_t) + birth\_time) which gives a sequence of the the event occurrence times to be randomly mapped upon the species tree lineages. Note, only those event occurrence times time of emergence for the locus/state until the time when the species tips are observed (= 1) are considered.

Iterating successively through the event timings, a number betweeen [0-1] is randomly drawn from a uniform distribution (prob\_event) and used to determine whether the given event is a loss or recombination:

- Extinction/Loss: if prob\_event <= Er / (Er + Rr); otherwise
- Recombination: prob\_event > Er / (Er + Rr)

In addition, a locus/state longevity rate (Lr) parameter can be incorporated, which will result in the inter-arrival event distribution of exp(-(L + Er + Rr)), but also add the possibility of no events occurring in the evolution of the locs/state:

- No Event (Locus/State Maintained) : if prob\_event <= Lr / (Lr + Er + Rr); otherwise
- Extinction/Loss : prob\_event <= Er / (Lr + Er + Rr); otherwise
- Recombination : prob\_event <= Rr / (Lr + Er + Rr)

If the event is a locus/state loss, species lineages possessing the locus/state at the given time are extracted and assigned as a loss event, and all descendant lineages occurring after the event time are also assigned as losses. If the event is a recombination, then a species lineage lacking the locus/state at the given time (if it exists) is randomly selected and assigned as a locus gain event, and its descendants are assigned as a locus gain event, in distinction to the initial locus/state origination event. In effect this generates a locus/state distribution presence/absences for the tips of the species tree, as well as the ancestral evolutionary histories of these traits. There may also be tips which never possessed the locus in their evolutionary history (absence). Using this approach we can examine how frequently locus/state distributions overlap between different evolutionary regimes, e.g. loss dominated vs. recombination dominated.

#### 2.1 Functions

##### 2.1.1 lineage\_get()

First step: Identify species lineages by traversing the species tree chronogram from the root (t = 0) to the tips (present time - t = 1) at a pre-defined intervals (defined time-step = delta). At each time step, find the coextant branches and their corresponding ancestral nodes they are descended from.

```
#lineage_get() : Identify all species (and thus potential locus/state lineages) present at a given timestep (time from the tree root).

lineage_get <- function(tree){
  
  #Get the nodepaths of all tips to the root:
  np <- nodepath(tree)
  
  #Get all the distances between tips and nodes
  dn <- dist.nodes(tree)
  
  #Get cumulative distances from root to each node in a path leading to a tip, these will be the time from the tree root when each node first appears: 
   dp <- lapply(np, function(x){
      dp <- NULL
      
      #initialize for descendants of the root:
      dp <- 0
      #Get the time at which the ancestor of a child node appears from its ancestor node:
      #Note, exclude the root and tips as they don't have ancestors or children nodes, respectively.
      
      for(i in 1:(length(x)-2)){
          #If there is more than one child node to the root, traverse down the nodepath to the tip, and assign emergence times
          #Of descendant nodes from their ancestors.
          if(length(x) > 2){
            dp <- append(dp, dn[x[i], x[i+1]])
          }
      }
      dp <- signif(cumsum(dp), 4)
      names(dp) <- x[-1]
      return(dp)
  })
   
  
  #Next, use the array of successive emergence times of ancestor -> descendant node pairs (dp) to the  get the duration times of the descendant/children nodes of each ancestral node:
  #The time at which the ancestor node split, and the time, i.e. branchlengths, of its descendant child nodes until they in turn bifrucate into 
  #their respective descendant nodes.
   
  lin_time <- data.frame(matrix(0, length(unique(names(unlist(dp)))), 2, dimnames=list(unique(names(unlist(dp))), c('begin', 'end'))))

  for(i in 1:length(dp)){
    for(j in 1:length(dp[[i]])){
      n <- names(dp[[i]][j])
      if(length(dp[[i]]) > 1 & j < length(dp[[i]])){
        lin_time[n,]$begin <- dp[[i]][j]
        lin_time[n,]$end <- dp[[i]][j+1]
      }
      else{
        lin_time[n,]$begin <- dp[[i]][j]
        lin_time[n,]$end <- dn[Ntip(tree)+1, n]
      }
    }
  }
  
  #Also, add the root node with begin and end times [-1, 0]:
  
  lin_time <- rbind.data.frame(root=c(0,0), lin_time)
  rownames(lin_time)[1] <- Ntip(tree) + 1
  
  return(lin_time)
}


#An illustration of the lineges identified:
####

#Generate a tree?:
tree_pb <- pbtree(b=0.5, n=10, scale=1)


lineage <- lineage_get(tree_pb)

delta <- 1e-1
plot(tree_pb,
    label.offset=0.05)

title(paste0('Species Lineages Identified', 'Time-Step = ', delta))


nodepos <- with(.PlotPhyloEnv$last_plot.phylo,
     cbind.data.frame(node=seq(1, Ntip+Nnode), x=xx, y=yy))


delta <- 1e-1
ts <- seq(0, 1, delta)
axis(1, ts, labels=ts)
mtext('Time From Root', 1, line=3)


#Label descendant branches/lineages present at given time slices.
for(i in 1:nrow(lineage)){
  n <- rownames(lineage)[i]
  if(i == 1){
     ti <- -2e-2
  }
  else{
    ti <- lineage[n,]$begin + (lineage[n,]$end - lineage[n,]$begin)/2
  }
  
  text(ti,
       nodepos[rep(n, length(ti)),]$y,
       nodepos[rep(n, length(ti)),]$node, 
       pos=3,
       cex=0.5)
}
```

##### 2.1.2 locus\_evolve()

Second step: simulate trait evolutionary histories on phylogenetic tree ancestral species lineages with a poisson process to model loss and recombination.

```
###Code for modelling locus/trait evolution on a species tree using a poisson process.

#Starting with the species tree. Draw from the uniform distribution the time when the gene lineage first appears. Assign it randomly to an extant species lineage at that given time. Get the descendant lineages from that time-point.

#Randomly draw locus/state lineage extinction times from the poisson distribution (with a defined death_rate). Randomly assign these losses to the descendant lineages present at each extinction time.


#Define a function to model gains, losses, and transfers of a locus/state on species lineages of a given species tree:
locus_evolve2 <- function(tree, lineage, death_r=0, recomb_r=0, long_r=1, root=FALSE){
  ###
  ###Step 1: Initialize Parameters and Variables:
  ###
  
  #Get the maximum tree length (root to tip) from the lineage matrix:
  tree_len <- max(lineage[,2])
  
  #The longevity rate (no event occurring):
  #long_r <- 0.5
  
  #The extinction rate:
  #death_r <- 1
  
  #The recombination rate:
  #recomb_r <- 0.5
  
  
  #The time when the locus/state lineage emerges:
  birth_time <- NULL
  
  #The corresponding species lineage where the gene arises at birth_time:
  birth_lin <- NULL
  
  #The species descendants from the birth_lineage:
  birth_descend <- NULL
  
  #The times when the locus/state lineage is lost:
  loss_time <- NULL
  
  #The corresponding species lineages where the locus/state is lost/goes extinct:
  loss_lin <- NULL
  
  #The species descendants of the loss_lineage:
  loss_descend <- NULL
  
  
  #The donor/recpient species lineages where a locus/state is acquired through recombination:
  donor_lin <- NULL
  recip_lin <- NULL
  
  #The species descendants of the recip_lineage:
  recip_descend <- NULL

  
  #Tally of tips descended from loss lineages, this doesn't seem to be used for anything.
  lt <- 0
  
  ###
  ###Step 2: Randomly select the time where the locus/state emerges:
  ###
  
  #Randomly draw a value between 0-1 from a uniform distribution:
  if(!root){
    #Now can set the birth of the locus/state originating at the root, i.e. if the time of its appearance is before time = 0.
    birth_time <- signif(runif(1, min=-0.1, max=1), 4)
    if(birth_time < 0){
      birth_time <- 0
    }
  }
  else{
    #Or set locus/state as present at the species tree root (0):
    birth_time <- 0
  }
  
  #Get the lineages existing within the given time:
  birth_lin <- rownames(lineage)[which(lineage$begin <= birth_time & lineage$end >= birth_time)]
  birth_lin <- as.numeric(birth_lin)
  
  #If multiple lineages exist at a given time, randomly select one:
  if(length(birth_lin) > 1){
    birth_lin <- sample(birth_lin, 1)
  }
  
  #print('birth')
  #print(birth_time)
  #print(birth_lin)
  
  #Get descendants of the birth lineage:
  birth_descend[[as.character(birth_time)]] <- unique(c(birth_lin, Descendants(tree, birth_lin, type='all')))
  
  
  ###
  ###Step3 : Model Locus/State Loss and Recombintion Events Using the Poisson Process.
  ###
  
  #Sample the Inverse Cumulative Probability Function of the Exponential Probability Density Distributions for Event Interrarival Times:
  eve_time <- -log(1-runif(20))/(death_r + long_r + recomb_r)
  
  #Cumulative sum the event inter-arrival times following the time when the locus/state originates. S
  eve_time <- birth_time + cumsum(eve_time)
  
  #Select only those event arrival times which occurr within the birth_time to the tip/present date of the tree
  eve_time <- signif(eve_time[which(eve_time > birth_time & eve_time <= tree_len)], 4)
  
  if(length(eve_time) != 0){
    
    #Iterate through the event timings, and determine whether losses or recombinations will occur.
    for(e_t in eve_time){
      
      #Check the probability of an event occurring: runif(1) >= long_r / (long_r + death_r + recomb_r);
      #If so, then check whether the event is a loss:
      #Probability of loss <= death rate / (death_rate + recomb_rate);
      #Otherwise it is a recombination.
      if(runif(1) > long_r / (long_r + death_r + recomb_r)){
        if(runif(1) <= death_r / (death_r + recomb_r)){
          #Get the lineage where a loss occurrs:
          loss_lin <- rownames(lineage)[which(lineage$begin <= e_t & lineage$end >= e_t)]
          loss_lin <- as.numeric(loss_lin)
          #loss_lin <- lineage[[as.character(signif(birth_time + ext_time[i], 1))]]
          
          #Check to see if the loss lineages haven't already been selected and are present in the birth and potential transfer lineages:
          #Keep from overwriting loss_lin when there is no overlap with the birth_descend lineages, i.e. when the locus only arises in a single outgroup... 
          loss_lin2 <- loss_lin[which(!loss_lin %in% unlist(loss_descend) & loss_lin %in% unlist(birth_descend))]
          
          #Have to check if any recipent transfer lineages have been selected:
          if(length(unlist(recip_descend)) != 0){
            loss_lin2 <- append(loss_lin2, loss_lin[which(loss_lin %in% unlist(recip_descend))])
          }
          
          loss_lin <- loss_lin2
          
          #If a potential loss lineage has been found:
          if(length(loss_lin) != 0){
            if(length(loss_lin) > 1){
              loss_lin <- sample(loss_lin, 1)
            }
          
            ld <- Descendants(tree, loss_lin, type='all')
          
            #Only add loss lineages which haven't been previously selected:
            if(length(which(ld %in% unlist(loss_descend))) == 0){
              #print('loss')
              #print(e_t)
              #print(loss_lin)
              
              loss_descend[[as.character(e_t)]] <- unique(c(loss_lin, ld))
              
              #Tally the number of tips lost?
              #lt <- lt + length(which(ld <= n_tip))
            }
          }
        }
        else{

          #Otherwise recombination occurs:
          
          #Get the lineage where a recombination event occurrs.
          #Have to make sure there are extant lineages at that given time to represent the donor:
          
          #Get lineages present at the current time:
          lin <- rownames(lineage)[which(lineage$begin <= e_t & lineage$end >= e_t)]
          lin <- as.numeric(lin)
          
          #Check if the donor lineages are present in the birth lineage, but have not yet been lost.
          i <- which(lin %in% unlist(birth_descend) & !lin %in% unlist(loss_descend))
          
          #If a donor lineage exists at the given event time, then select a potential recipient lineage present at the same time:
          if(length(i) != 0){
            if(length(i) == 1){
              donor_lin <- lin[i]
            }
            else{
              donor_lin <- sample(lin[i], 1)
            }
            
            #Choose a recipient lineage from the non-birth descendant lineages, make sure to exclude revious recipient lineages if they exist.
            
            #####What about reaquisition into a loss lineage? 
            i <- which(!lin %in% unlist(birth_descend))
            j <- which(lin %in% unlist(recip_descend))
            
            if(length(j) != 0){
              i <- c(i, j)
            }
            
            recip_lin <- lin[i][which(!lin[i] %in% unlist(recip_descend))]
            
            if(length(recip_lin) != 0){
              if(length(recip_lin) != 1){
                recip_lin <- sample(recip_lin, 1)
              }
              
              #Get descendants of the recipient lineage:
              rd <- Descendants(tree, recip_lin, type='all')
              
              recip_descend[[as.character(e_t)]] <- unique(c(recip_lin, rd))
              
              #print('transfer')
              #print(e_t)
              
              #print(recip_lin)
            }
          }
        }
      }
    }
  }
  
  return(list(birth_descend=birth_descend, loss_descend=loss_descend, recip_descend=recip_descend, params=list(long_r=long_r, death_r=death_r, recomb_r=recomb_r)))
}
```

##### 2.1.3 event\_label()

Third step: summarize the events observed for each simulated trait evolutionary history.

```
#Define a function for assigning final locus/state presence/absence annotations to tips:
event_label <- function(tree, evolve_res){
  #1) Assign all tips and internal nodes as absent (i.e not possessing the locus in an ancestor)
  #2) Assign presence to any tip nodes found in the birth_descend and recip_descend lists
  #3) Assign loss to any tip nodes found in the loss_descend list
  
  #Extract the birth, loss, and recipient lineages assigned to the species tree given by locus_evolve2()
  birth_descend <- evolve_res$birth_descend
  loss_descend <- evolve_res$loss_descend
  recip_descend <- evolve_res$recip_descend
  
    
  #Iterate through all nodes in the tree, and extract the times associated with 
  #Birth (first emergence of locus/trait), transfer, and loss.
  
  #For each node assign:
  #Gain - if the latest event prior to the node is a birth (locus/state origination) with no subsequent losses
  #gain-transfer - if the latest event prior to the node is a transfer (no subsequent losses)
  #Loss - if the latest event prior to a node is a loss (with no subsequent transfers)
  #Absent - if there are no birth, or recipient events occurring.
  
  node_label <- NULL
  
  for(i in 1:(Ntip(tree) + Nnode(tree))){
    t_b <- NA
    t_l <- NA
    t_r <- NA
    
    for(t in names(c(birth_descend, recip_descend, loss_descend))){
      if(i %in% birth_descend[[t]]){
        t_b <- as.numeric(t)
      }
      
      if(i %in% recip_descend[[t]]){
        t_r <- as.numeric(t)
      }
      
      if(i %in% loss_descend[[t]]){
        t_l <- as.numeric(t)
      }
    }
    
    type <- NA
    #Birth lineage without losses:
    if(!is.na(t_b) & is.na(t_l)){
      type <- 'gain'
    }
    #Transfer lineage without losses:
    else if(is.na(t_b) & is.na(t_l) &!is.na(t_r)){
      type <- 'gain-transfer'
    }
    #Loss lineage in birth lineage:
    else if(!is.na(t_b) & !is.na(t_l) & is.na(t_r)){
      type <- 'loss'
    }
    #Gains via transfers in loss lineage or losses in tranfer lineages:
    else if(!is.na(t_l) & !is.na(t_r)){
      if(t_l > t_r){
        type <- 'loss'
      }
      else if(t_l < t_r){
        type <- 'gain-transfer'
      }
    }
    #Lineage with no locus/state acquired:
    else if(is.na(t_b) & is.na(t_l) & is.na(t_r)){
      type <- 'absent'
    }
    
    node_label <- rbind.data.frame(node_label, data.frame(node=i, birth_time=t_b, trans_time=t_r, loss_time=t_l, node_type=type))
    #print(c(i, t_b, t_r, t_l, type))
  }
  
  return(node_label)
}
```

##### 2.1.4 plot\_evo\_tree()

A plotting function to visualize evolved trait distributions.

```
#Plotting function for evolved locus/state lineages on chronograms or phylograms:
plot_evo_tree <- function(tree, evolve_res, node_label, lin=FALSE, cex=1){

  #Color branches by their locus/state type:
  #Green indicates the ancestral node and descendant lineages where the locus/state was first acquired (birth_descend)
  #Red indicates the ancestral node and descendant lineages where the locus/state was lost (loss_descend)
  #Purple indicates the ancestral node and descendant lineages where the locus/state was acquired through transfer (recip_descend)
  
  #e_col <- rep('black', nrow(tree_pb$edge))
  
  #Assign branch colours according to the times they appear.
  
  # e_col[which(tree_pb$edge[,1] %in% unlist(birth_descend) & tree_pb$edge[,2] %in% unlist(birth_descend))] <- 'green'
  # e_col[which(tree_pb$edge[,1] %in% unlist(loss_descend) & tree_pb$edge[,2] %in% unlist(loss_descend))] <- 'red'
  # e_col[which(tree_pb$edge[,1] %in% unlist(recip_descend) & tree_pb$edge[,2] %in% unlist(recip_descend))] <- 'purple'
  
  
  #Another way for annotating edges:
  {
    e_col <- rep('black', nrow(tree$edge))
    
    #Gains (by initial birth):
    gain_n <- node_label$node[grep('^gain$', node_label$node_type)]
    e_col[which(tree$edge[,1] %in% gain_n & tree$edge[,2] %in% gain_n)] <- 'green'
    
    #Gains (by recombination):
    trans_n <- node_label$node[grep('gain-transfer', node_label$node_type)]
    e_col[which(tree$edge[,1] %in% trans_n & tree$edge[,2] %in% trans_n)] <- 'purple' 
    
    #Losses:
    loss_n <- node_label$node[grep('loss', node_label$node_type)]
    e_col[which(tree$edge[,1] %in% loss_n & tree$edge[,2] %in% loss_n)] <- 'red' 
  }
  
  #Plot the tree annotated with locus distributions
  
  plot(tree, edge.color=e_col, show.tip.label=FALSE, edge.width=2)
  #nodelabels(node_label[(1+Ntip(tree_pb)):(Nnode(tree_pb) + Ntip(tree_pb)),5])
  #tiplabels(node_label[1:Ntip(tree_pb),5])
  
  #Get node x and y plotting coordinates:
  nodepos <- with(.PlotPhyloEnv$last_plot.phylo,
       cbind.data.frame(node=seq(1, Ntip+Nnode), x=xx, y=yy))
  
  #X-axis: time from tree root.
  
  delta <- 1e-1
  
  
  axis(1, seq(0, max(nodepos$x), delta), labels=seq(0, max(nodepos$x), delta))
  mtext('Distance/Time From Root', 1, line=3, cex=cex/2)
  
  
  #Annotate lineages?
  if(lin){
    for(n in rownames(lineage)){
      ti <- lineage[n,]$begin + (lineage[n,]$end - lineage[n,]$begin)/2
      text(ti,
           nodepos[rep(n, length(ti)),]$y,
           nodepos[rep(n, length(ti)),]$node,
           pos=3,
           cex=0.5)
    }
  }  
  
  #Add additional branchlengths beginning from the time slice on a given descendant branch where the loss occured, and ending #at the decendant node position in the tree. Use nodepos to get the appropriate y position and horizontal end position.
  
  birth_descend <- evolve_res$birth_descend
  loss_descend <- evolve_res$loss_descend
  recip_descend <- evolve_res$recip_descend
  
  
  #Gains:
  for(i in names(birth_descend)){
    j <- birth_descend[[i]][1]
    points(c(as.numeric(i), nodepos[j,]$x), c(nodepos[j,]$y, nodepos[j,]$y), type='l', col='green', lwd=2)
    points(c(as.numeric(i), as.numeric(i)), c(0, nodepos[j,]$y), type='l', lty=2, col='green', xpd=TRUE)
    points(as.numeric(i), nodepos[j,]$y, type='p', pch='X', lty=2, col='green', xpd=TRUE)
  }
  
  #Losses:
  for(i in names(loss_descend)){
    j <- loss_descend[[i]][1]
    points(c(as.numeric(i), nodepos[j,]$x), c(nodepos[j,]$y, nodepos[j,]$y), type='l', col='red', lwd=2)
    points(c(as.numeric(i), as.numeric(i)), c(0, nodepos[j,]$y), type='l', lty=2, col='red', xpd=TRUE)
    points(as.numeric(i), nodepos[j,]$y, type='p', pch='X', lty=2, col='red', xpd=TRUE)
  }
  
  #For transfers:
  for(i in names(recip_descend)){
    j <- recip_descend[[i]][1]
    points(c(as.numeric(i), nodepos[j,]$x), c(nodepos[j,]$y, nodepos[j,]$y), type='l', col='purple', lwd=2)
    points(c(as.numeric(i), as.numeric(i)), c(0, nodepos[j,]$y), type='l', lty=2, col='purple', xpd=TRUE)
    points(as.numeric(i), nodepos[j,]$y, type='p', pch='X', lty=2, col='purple', xpd=TRUE)
  }
  
  #Calculate the actual phylogenetic distance for the evolved locus distribution:
  
  #Get all of the nodes which have gained a locus/state (through original generation or transfer, with subsequent lossess occurring):
  gain_n <- node_label$node[grep('gain', node_label$node_type)]
  
  #presence/absence of branches, for comparison with recpd results:
  br_pa <- ifelse(tree$edge[,1] %in% gain_n & tree$edge[,2] %in% gain_n, 1, 0)
  
  #The adjusted phylogenetic diversity of the locus distribution:
  pd_actual <- sum(tree$edge.length[which(br_pa == 1)]) / sum(tree$edge.length)

  title(paste('Actual PD = ', signif(pd_actual, 3)), cex=cex)
}


tr_norm <- rtree(50)
tr_norm$edge.length <- tr_norm$edge.length/max(dist.nodes(tr_norm)[1:Ntip(tr_norm), Ntip(tr_norm)+1])

lineage <- lineage_get(tr_norm)

par(mfrow=c(3,3))

for(i in 1:9){
  
  evolve_res <- locus_evolve2(tr_norm, lineage, death_r=2, recomb_r=3, long_r=1, root=TRUE)
  
  node_label <- event_label(tr_norm, evolve_res)
  
  plot_evo_tree(tr_norm, evolve_res, node_label, cex=1)
}
```

### 3 Simulate trait evolutionary histories

All the basic pieces are in place. Wrap this up to simulate locus/state distributions on the species tree using the poisson process, altering death, recombination, and longevity rates. For each run, calculate the prevalence of a locus/state, number originating initially through birth, number originating through transfer, and number lost.

Using the tip locus/state presence/absence patterns resulting from the simulated poisson process, calculate RecPD vs. Faith’s PD and compare against against the actual PD given for simulated locus/state histories.

```
#Make a function to generate the evolved locus/state distributions and calculate PDs:

run_evo <- function(tip_range, n_tree, rate_mat, n_iter, type=c('chronogram', 'phylogram')){
  #Initialize a data.frame for storing results:
  test_res <- list()
  n <- 1
  
  #First iterate through trees of different sizes (# of tips), single trees, but can generate multiple trees with a given number of tips...
  
  #By storing the the evolve_locus2() results and branches for simulated locus/state histories, one can also perform correlation calculations.
  for(n_tip in tip_range){#seq(25, 100, 25)){
    
    if(type[1] == 'chronogram'){
      #Generate a birth-death species tree:
      tree_l <- pbtree(b=0.5, n=n_tip, scale=1, nsim=n_tree)
    }
    else{
      #Or a random distance based tree, midpoint rooted?:
      tree_l <- rmtree(n_tree, n_tip)
    }
    
    #For each simulated tree:
    for(t_i in 1:length(tree_l)){
      
      tree_pb <- tree_l[[t_i]]
      
      if(type[1] == 'phylogram'){
        #Normalize tree branch lengths so that tree depth == 1
        tree_pb$edge.length <- tree_pb$edge.length/max(dist.nodes(tree_pb)[1:n_tip, n_tip+1])
        
        #Midpoint root the tree (if a phylogram)?:
        #tree_pb <- midpoint.root(tree_pb)
      }
      
      #Generate the locus lineages for the species tree, required for the locus_evolve2() function:
      lineage <- lineage_get(tree_pb)
      
      #For the recpd calculations:
      #Generate a node_sp list of tree nodes and their nearest-neighbour tip descendant:
      nn <- nn_nodes(tree_pb)
  
      #print(paste(n_tip, t_i))
      
      for(i in 1:nrow(rate_mat)){
        #Set rates:
        l_r <- rate_mat[i,1]
        d_r <- rate_mat[i,2]
        r_r <- rate_mat[i,3]
        
        #Number of simulated locus/state histories to perform:
        for(j in 1:n_iter){
          
          evolve_res <- locus_evolve2(tree_pb, lineage, death_r=d_r, recomb_r=r_r, long_r=l_r, root=FALSE)
          
          node_label <- event_label(tree_pb, evolve_res)
          
          #Gains from birth: 
          #Number of events based on times mapped to the tree:
          gain_n <- length(evolve_res$birth_descend)
          
          #(tips which originated from initial birth lineage):
          gain_nt <- node_label$node[grep('^gain$', node_label$node_type)]
          gain_nt <- length(which(gain_nt <= Ntip(tree_pb)))
          
          #Gains from transfer:
          #Number of events based on times mapped to the tree:
          trans_n <- length(evolve_res$recip_descend)
          
          #(tips which originated from transfer lineages):
          trans_nt <- node_label$node[grep('gain-transfer', node_label$node_type)]
          trans_nt <- length(which(trans_nt <= Ntip(tree_pb)))
          
          #Losses 
          loss_n <- length(evolve_res$loss_descend)
          
          #(tips lost from both origination and transfer lineages):
          loss_nt <- node_label$node[grep('loss', node_label$node_type)]
          loss_nt <- length(which(loss_nt <= Ntip(tree_pb)))
          
  
             
          #Tip Presence/Absence
          pa <- data.frame(
            matrix(ifelse(1:Ntip(tree_pb) %in% grep('gain', node_label[1:Ntip(tree_pb),]$node_type), 1, 0), 
                 1, Ntip(tree_pb), 
                 dimnames=list(1, tree_pb$tip.label))
          )
          
          ####
          #Phylogenetic Diversity based on evolutionary history:
          
          #Only calculate pd if at least two tips exist:
          if(sum(pa) > 1){
            
            #Branches present in the simulated locus distribution:
            pres <- node_label$node[grep('gain', node_label$node_type)]
            
            #Include ancestors of the gained nodes? 
            pres <- unique(c(pres, Ancestors(tree_pb, pres, 'parent')))
          
            #Or, only if the children of present nodes are both descenants of the root:
            # if(length(which(pres %in% Descendants(tree_pb, Ntip(tree_pb) +1, type='children')))){
            #   pres <- append(pres, Ntip(tree_pb)+1)
            # }
  
            #Actual PD based on the simulated evolutionary history of the tree:
            br_pa <- ifelse(tree_pb$edge[,1] %in% pres & tree_pb$edge[,2] %in% pres, 1, 0)
            pd_orig <- sum(tree_pb$edge.length[which(br_pa == 1)]) / sum(tree_pb$edge.length)
          
          
            #Phylogenetic Diversity of the tip presence/absence pattern using RecPD
            #Calculate RecPD, using the 'nn', 'mpr', and 'ace' methods for node state assignments:
            r_nn <- recpd_calc(tree_pb, pa, 'nn')
            r_mpr <- recpd_calc(tree_pb, pa, 'mpr')
            r_ace <- recpd_calc(tree_pb, pa, 'ace')
            
            r_nn <- r_nn[[1]]$metrics$recpd
            r_mpr <- r_mpr[[1]]$metrics$recpd
            r_ace <- r_ace[[1]]$metrics$recpd
            
            #What about comparing how successful the different approaches are at identifying potential recombination and loss internal nodes?
            
            #Check if any transfers are present:
            #Get the gain-transfer nodes from the simulated locus/state history:
            t_n <- which(node_label$node_type == 'gain-transfer')
            
            #Get the recpd node_state assignments:
            #MPR method, for example...
            ####Expand this for other RecPD methods (nn, and ace)...
            ns <- node_state_mpr(tree_pb, pa[1,])
            
            #Number of matches between transfer nodes and split nodes predicted by RecPD:
            t_a <- 0
            t_m <- 0
            
            if(length(t_n) != 0){
              #Get the time(s) and lineages where the transfered occurred, then grab the ancestral node(s) of the lineages where it occurred:
              t_t <- unique(node_label$trans_time[t_n])
              t_d <- as.numeric(unlist(lapply(evolve_res$recip_descend[as.character(t_t)], function(x) x[1])))
              t_a <- unlist(Ancestors(tree_pb, t_d, type='parent'))
              
              #Check if any split nodes exist:
              if(length(which(ns == 2)) != 0){
              
                #Get the descendants of the split node (=2), select the one which is present as the likely transfer node:
                ns_t <- unlist(lapply(Descendants(tree_pb, which(ns == 2), type='children'), function(x) x[which(ns[x] == 1)]))
                
                #Check if the ancestor node(s) of the transfer-gain event(s) is(are) also the descendants of the split nodes
                t_m <- length(which(ns_t %in% t_a))
              }
              
              #Convert t_a to the number of transfer nodes:
              t_a <- length(t_a)
            }
            
            #Losses which occurr at internal nodes only....
            
            #Calculate Faith's PD:
            faith <- pd(pa, tree_pb, include.root=TRUE)$PD
            
            faith <- faith/sum(tree_pb$edge.length)
            
            test_res[[n]] <- c(n_tip=n_tip, n_tree=t_i,
                                          long_r=l_r, 
                                          death_r=d_r, 
                                          recomb_r=r_r, 
                                          prev=sum(pa), 
                                          num_birth_event=gain_n,
                                          num_trans_event=trans_n,
                                          num_loss_event=loss_n,
                                          num_birth_tip=gain_nt, 
                                          num_trans_tip=trans_nt, 
                                          num_loss_tip=loss_nt,
                                          pd_orig=pd_orig,
                                          num_split=length(which(ns == 2)),
                                          t_m=t_m,
                                          recpd_nn=r_nn,
                                          recpd_mpr=r_mpr,
                                          recpd_ace=r_ace,
                                          faith=faith)
            
            n <- n + 1
          }
        }
      }
    }
  }
  
  #Format the output table into a data.frame:
  test_res <- data.frame(matrix(unlist(test_res), length(test_res), length(test_res[[1]]), by=2, dimnames=list(1:length(test_res), names(test_res[[1]]))))
  
  #Add an additional column to classify parameter sets:
  param_lab <- apply(test_res, 1, function(x){
      if(x[4] == x[5]){
        return('loss = recomb')
      }
      else if(x[4] > x[5]){
        return('loss > recomb')
      }
      else if(x[4] < x[5]){
        return('recomb > loss')
      }
    }
  )
  
  test_res <- cbind(test_res, param_set=param_lab)
  
  return(test_res)
}
```

Perform locus/state evolution simulations and PD calculation comparisons:

```
#Generate different sets of input rates:
rate_mat <- expand.grid(long=seq(0), death=seq(0, 10, 2), recomb=seq(0, 10, 2))

#Number of iterations of locus_evolve2() to perform for a given set of rates and tree:
n_iter <- 5

#Generate a range of tree sizes:
tip_range <- 50 #seq(50, 200, 50)

#Or perform these analyses separately for trees of different size...

#Number of trees to generate:
n_tree <- 1

#Note, these parameters are selected to compute in a reasonable time and as you will notice in the following code chunks, are not exhaustive. For more robust simulations, you would want to increase the range of tree sizes (tip_range), number of distinct tree topologies generated (n_tree), the number of simulations performed per tree (n_iter).

#Simulate trait distributions:
test_res <- run_evo(tip_range, n_tree, rate_mat, n_iter, type='phylogram')
```

Evaluate the number of loss and recombination events identified for simulations evolved under increasing loss and recombination rates.

```
#Show the average number of loss and transfer events predicted for each tree...

test_event <- test_res %>% 
  #mutate(param_set=param_lab) %>%
  pivot_longer(c(num_loss_event, num_trans_event), names_to='event', values_to='num_event') %>%
  select(n_tip, prev, long_r, death_r, recomb_r, param_set, event, num_event)


#For Increasing Recombination Rate:

event_bp_recomb <- ggplot(test_event, aes(x=factor(recomb_r), y=num_event, color=event)) +
  geom_boxplot(width=0.5, outlier.size=0.5) +
  facet_grid(param_set ~.) +
  scale_color_discrete(name='Event Type', labels=c('Loss', 'Transfer')) + 
  labs(title='Loss and Recombination Events by Rate Regime',
       x='Recombination Rate',
       y='Number of Events')

#count the number of simulations performed for each parameter set...
sim_count_recomb <- test_res %>% group_by(param_set, recomb_r) %>% summarize(count=n())

n_sims_recomb <- ggplot(sim_count_recomb, aes(x=factor(recomb_r), y=count, color=param_set)) +
  geom_point() +
  ylim(0,NA) +
  facet_grid(vars(param_set), scales='fixed') +
  labs(title='Number of Simulations by Rate Regime',
       x='Recombination Rate',
       y='Number of Simulations',
       color='Rate Regime')

ggarrange(n_sims_recomb, event_bp_recomb, ncol=2, legend='bottom')
```

```
#For Increasing Loss Rate:
event_bp_loss <- ggplot(test_event, aes(x=factor(death_r), y=num_event, color=event)) +
  geom_boxplot(width=0.5, outlier.size=0.5) +
  facet_grid(param_set ~.) +
  scale_color_discrete(name='Event Type', labels=c('Loss', 'Transfer')) + 
  labs(title='Loss and Recombination Events by Rate Regime',
       x='Loss Rate',
       y='Number of Events')

#count the number of simulations performed for each parameter set...
sim_count_loss <- test_res %>% group_by(param_set, death_r) %>% summarize(count=n())

n_sims_loss <- ggplot(sim_count_loss, aes(x=factor(death_r), y=count, color=param_set)) +
  geom_point() +
  ylim(0,NA) +
  facet_grid(vars(param_set), scales='fixed') +
  labs(title='Number of Simulations by Rate Regime',
       x='Loss Rate',
       y='Number of Simulations',
       color='Rate Regime')

ggarrange(n_sims_loss, event_bp_loss, ncol=2, legend='bottom')
```

#### 3.1 RecPD of simulated trait distributions

Plot out the differences between actual PD vs. calculated RecPD of simulated trait distributions.

```
#Group together runs by tree size (n_tip) and prevalence (prev), and calculate  mean and standard deviation values for each pd metric: pd_orig, faith, recpd_nn, recpd_mpr, repc_ace, and identical split node (transfers) identified (only calculated for recpd_mpr method at this point).


test_parse <- test_res %>% 
  #mutate(param_set=param_lab) %>%
  pivot_longer(c(faith, recpd_nn, recpd_mpr, recpd_ace), names_to='method', values_to='PD') %>% 
  select(n_tip, long_r, param_set, num_trans_event, num_loss_event, prev, pd_orig, method, PD)

#Long-format table for phylogenetic diversity estimates:
pd_tab <- test_res %>% 
  #select(prev, num_trans, num_loss, num_birth, long_r, death_r, recomb_r, pd_orig, recpd, faith) %>% 
  pivot_longer(cols=c(recpd_nn, recpd_mpr, recpd_ace, faith))

  

#Plotting results of the simulations:


###
#A labelling array for tree size (n_tip):
n_tip_lab <- array(paste('Ntip = ', unique(pd_tab$n_tip)), dimnames=list(unique(pd_tab$n_tip)))

#Labelling for loss rates:
death_r_lab <- array(paste('Loss Rate = ', unique(pd_tab$death_r)), dimnames=list(unique(pd_tab$death_r)))

#Labelling for recombinatino rates:
recomb_r_lab <- array(paste('Recomb Rate = ', unique(pd_tab$recomb_r)), dimnames=list(unique(pd_tab$recomb_r)))


#Estimated vs. Actual PD differences for the recomb > loss regime:
plt1 <- ggplot(test_parse %>% filter(param_set == 'recomb > loss'), aes(pd_orig, PD, color=PD/pd_orig)) + #color=factor(n_tip))) + #group=cut(pd_orig, breaks=seq(0, 1, 1e-1), include.lowest=TRUE))) +
  #geom_boxplot() +
  geom_point(size=0.2) +
  geom_abline(slope=1, intercept=0, lty=2, size=0.2) +
  #geom_smooth() +
  facet_grid(~method, scales='free') +
  #facet_grid(param_set ~ method) +
  scale_color_gradient2(name='Estimated /\nActual PD', 
                        midpoint=1, 
                        low='red', mid='blue', high='red',
                        na.value='red3',
                        limits=c(0,2)) +
  labs(title='Phylogenetic Diversity Comparisons: Actual Vs. Estimated PD by Method', 
      subtitle=paste0('Recombination > Loss (', 
                    nrow(test_res %>% filter(param_set == 'recomb > loss')), 
                    ' Simulations)'),
       x='Actual Phylogenetic Diversity',
       y='Estimated Phylogenetic Diversity') +
  theme(legend.key.size = unit(0.5, 'cm'),
        legend.text=element_text(size=10),
        axis.text.x=element_text(hjust=1, angle=45, size=10),
        axis.text.y=element_text(size=10),
        axis.title.x=element_text(size=12),
        axis.title.y=element_text(size=12),

        strip.text.x=element_text(size=12),
        plot.margin=unit(c(20,0,0,20), 'points'))


#Density plot distribution of Estimated - Actual PD differences for the recomb > loss regime:
plt2 <- ggplot(pd_tab %>% filter(param_set == 'recomb > loss'),
       aes(x=value/pd_orig, 
           lty=factor(recomb_r), 
           color=factor(name), 
           fill=factor(name)
           )
       ) +
  geom_vline(xintercept=1, lty=2, lwd=0.5) +
  geom_density(aes(y=..density..), alpha=0.1) +
  facet_wrap(vars(factor(name)),
             nrow=1, ncol=4,
             scales='free',
             labeller=labeller(recomb_r=recomb_r_lab, n_tip=n_tip_lab)) +
  labs(title='Actual / Estimated PD Density Distribution by Method',
      x='Estimated / Actual PD',
      y='Density',
      lty='Recombination\nRate',
      color='Method',
      fill='Method') + 
  theme(legend.key.size = unit(0.5, 'cm'),
        legend.text=element_text(size=10),
        axis.text.x=element_text(size=10),
        axis.text.y=element_text(size=10),
        axis.title.x=element_text(size=12),
        axis.title.y=element_text(size=12),
        strip.text.x=element_text(size=12),
        plot.margin=unit(c(20,0,0,20), 'points'))


#Combine Actual vs. Estimate Scatterplot (see code chunk above - plt1) with PD Difference Density Plot for Recomb > Loss Regime:

ggarrange(plt1, plt2, nrow=2)
```

```
## Warning in grid.Call.graphics(C_polygon, x$x, x$y, index): semi-transparency is
## not supported on this device: reported only once per page
```

```
#All actual PD - estimated PD differences by rate regime:

ggplot(test_parse, aes(pd_orig, PD, color=PD/pd_orig)) +
  #geom_boxplot() +
  geom_point(size=0.2) +
  geom_abline(slope=1, intercept=0, lty=2, size=0.2) +
  #geom_smooth() +
  facet_grid(param_set ~ method, scales='free') +
  #facet_grid(param_set ~ method) +
    scale_color_gradient2(name='Estimated /\nActual PD', 
                        midpoint=1, 
                        low='red', mid='blue', high='red',
                        na.value='red3',
                        limits=c(0,2)) +
  labs(title='Phylogenetic Diversity Comparisons\nActual Vs. Estimated',
       subtitle=paste(nrow(test_res), 'Simulations'),
       x='Actual Phylogenetic Diversity',
       y='Estimated Phylogenetic Diversity') +
  theme(axis.text.x=element_text(hjust=1, angle=45),
        legend.key.size = unit(0.5, 'cm'))
```

```
#boxplot summaries?

ggplot(pd_tab, aes(x=name, y=value-pd_orig, fill=name))  +
  geom_boxplot(outlier.size=0.5)
```

```
# #Calculate the mean and stdev of estimated / actual PD values - 1 by PD method.
# pd_tab %>% 
#     group_by(name) %>% 
#     summarize(mean(value/pd_orig - 1), sd(value/pd_orig -1))
# 
# #wilcox text P-value difference of NN vs. MPR:
# 
# wilcox.test(test_res$recpd_nn/test_res$pd_orig - 1, test_res$recpd_mpr/test_res$pd_orig -1)
# 
# #wilcox test p-value difference of NN with pd_orig
# wilcox.test(test_res$recpd_nn, test_res$pd_orig)
```
